# The Muscle Tissue Environment Limits Muscle Stem Cells in Aged Mice

**DOI:** 10.64898/2026.04.10.717808

**Authors:** Alicia A. Cutler, Tenaya K. Vallery, Thomas O. Vogler, Jesse Kurland, Thea S. Zlatklov, Tiffany Antwine, Nicole Dalla Betta, Tze-Ling Chang, Brad Pawlikowski, Carson H. Butcher, Kory J. Lavine, David M. Ornitz, Kristi S Anseth, Bradley B. Olwin

## Abstract

Frailty arising from loss of muscle function and mass is a significant health concern impacting quality of life and dramatically increasing health care costs as our population ages. Ameliorating frailty derived from reduced muscle function is thus a critical research priority to improve health span. Cell intrinsic defects in muscle stem cells (MuSC), or satellite cells, occur as skeletal muscle ages, reducing the capacity of MuSCs to maintain and repair skeletal muscle and are accompanied by cell nonautonomous changes. Although rejuvenating stem cells in aged tissues or organs has potential to improve muscle aging phenotypes, we found that the extracellular environment in aged mice abrogates rejuvenated muscle stem cell potential. MuSCs from young mice were unable to grow on extracellular matrix derived from aged mice that contains elevated collagen protein levels, establishing a critical role for the environment in contributing to muscle phenotypes in aging. Combining an inducible FGF receptor 1 (FGFR1) to rescue MuSC intrinsic aging defects with a drug to reduce fibrosis partially rescued muscle mass loss in aged mice. We conclude that aging affects tissues, and particularly skeletal muscle tissue, via complex multifactorial processes requiring multifaceted interventions to improve aging phenotypes.

## Introduction

Skeletal muscle mass, function, and regenerative capacity decline with age, leading to increased frailty and morbidity in the elderly^1–4^. By 2050, 20% of the world population will be over the age of 60^5^, resulting in a predicted 18% increase in overall health care expenditures^6^. Increasing health span and quality of life for the elderly by ameliorating aging related muscle declines will decrease societal burden and improve quality of life for the increasingly aging population.

Skeletal muscle in elderly individuals has reduced mass, fewer MuSCs, and increased fibrosis compared to younger individuals^3,7–9^. Skeletal muscle regenerates poorly in the elderly, in part, due to cell intrinsic aging defects in MuSCs that have inherently reduced regenerative capacity in aged mammals^10,11^, reduced self-renewing asymmetric divisions^12,13^, and increased spontaneous premature differentiation^12,14^. Although exposing aged muscle to a young environment improves regeneration^15,16^, transplanting MuSCs from aged muscle into young mice does not rescue the age-associated self-renewal deficit^12^. Moreover, transplanting MuSCs from young mice fails to rescue muscle phenotypes in aged mice, suggesting both cell autonomous and cell nonautonomous changes occurring during aging affect muscle maintenance and repair. MuSCs integrate and respond to extracellular cues, including fibroblast growth factor (FGF) signaling, which is essential for determining cell fate and function during regeneration, diminishes with age^12,17^. Consistent with this idea, a drug-activated FGFR1 partially rescues self-renewal in MuSCs from aged mice and partially rescues MuSC engraftment for transplants from aged mice to young mice^12^. MuSC cell nonautonomous signaling declines with age as activating ß1-integrin improves FGF signaling in muscle of aged mice and rescues myofiber size and strength in a mouse model of Duchenne muscular dystrophy^17^. Based on data from these findings, we found that partially rejuvenating MuSCs *in vivo* while concomitantly improving the muscle environment in aged mice improved the regenerative capacity of muscle in aged mice, restoring muscle weight size to a more youthful phenotype.

## Results

Mice were bred for temporally inducible caFGFR1^18^ expression specifically in MuSCs to rejuvenate MuSCs (Fig. 1A). The caFGFR1 transgene is a chimeric receptor fusing the mutant FGFR3c(R248C) extracellular and transmembrane domains^19^ to the intracellular FGFR1 tyrosine kinase domain with a C-terminal MYC epitope tag^20,21^. To determine the efficiency of the genetic system including three transgenes, *Pax7^CreERT2^*; *LSL-rtTA*; *tetO:caFGFR^+^* (caFGFR1+) mice and *Pax7^CreERT2^*; *LSL-rtTA* (caFGFR1−), control mice were tamoxifen treated for 5 days and fed standard chow or doxycycline-containing chow for 5 days. Myoblasts were then isolated and cultured in the presence or absence of doxycycline. The myoblasts were scored for immunoreactivity for PAX7, MYC, and FGFR3 (the extracellular domain of caFGFR1 that is not expressed in MuSCs or their progeny)^22^ (Fig. 1B). In caFGFR1+ cultures treated with doxycycline, 58% of the PAX7+ cells were immunoreactive for MYC or FGFR3 (Fig. 1C). While *Gapdh* transcripts were detectable by RT-PCR under all conditions, caFGFR1 transgene expression was maintained only in the presence of doxycycline (Fig. 1D). Thus, caFGFR1 expression is inducible and reversible.

**Figure 1.**
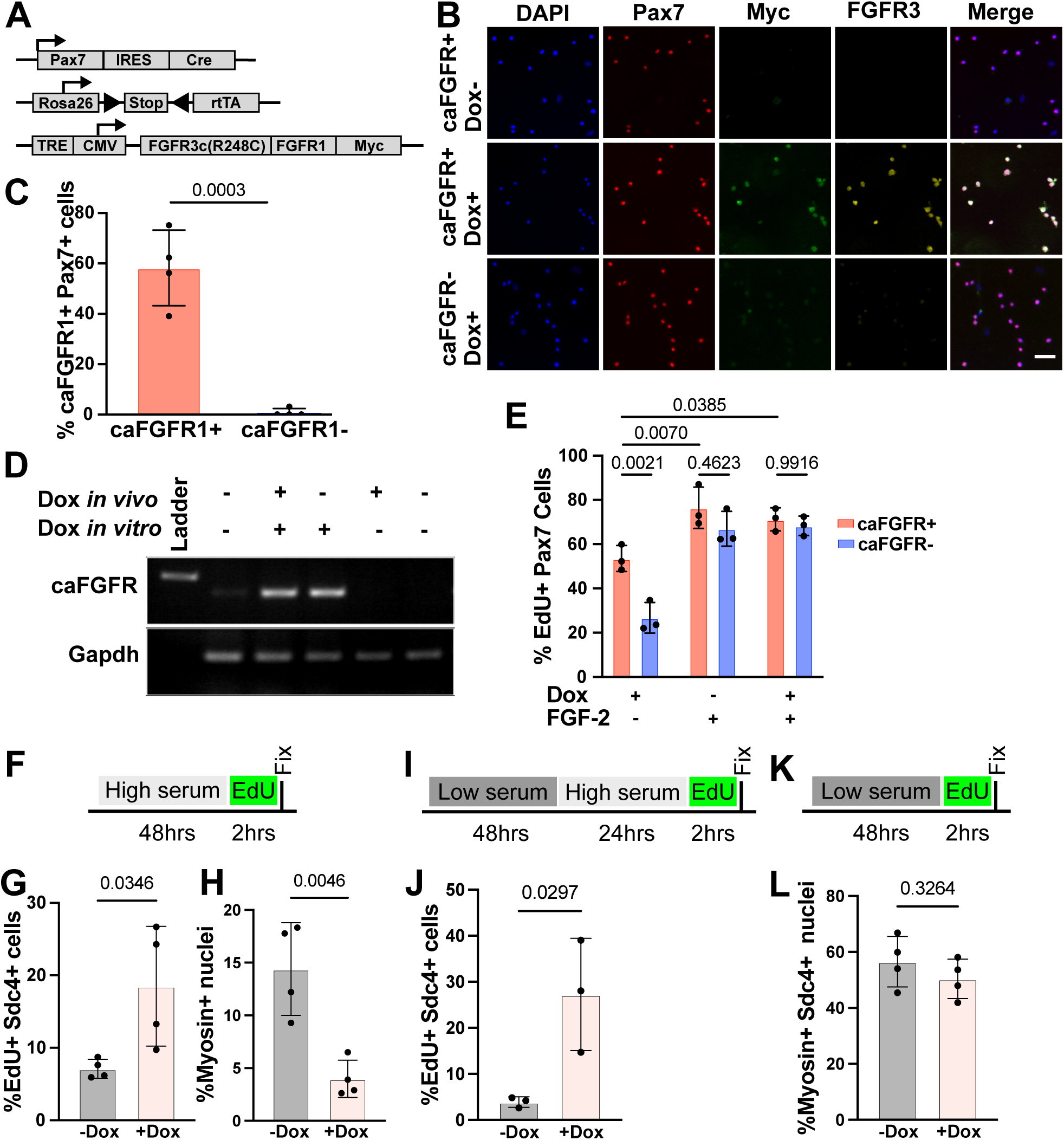
Constitutively active FGFR1 expression rescues MuSC cell numbers in aged mice. A) Schematic of caFGFR1 mouse genetics. B) Immunofluorescence images of MuSCs isolated from caFGFR1+ and caFGFR1− mice treated with doxycycline or vehicle and tested for immunoreactivity to PAX7 (MuSC marker), MYC, and FGFR3 (the extracellular domain of caFGFFR1). Scale bar = 50 μm. C). Quantification of percentage of Pax7+ cells positive for FGFR3 (n=4 primary myoblast isolates from young mice), comparison by t-test. D) RT-PCR gels of transcripts isolated from caFGFR1+ cells treated with doxycycline *in vivo* and/or *in vitro*. Primers to the chimeric caFGFR1 transcript or GAPDH were provided. E) Quantification of precent EdU positive Pax7+ cells treated with doxycycline (Dox), FGF2, or both. n=3 primary myoblast isolates from young mice, comparisons by one-way ANOVA. F) Experimental approach schematic for primary myoblasts isolated from old (20–24-month-old) caFGFR1+ mice. G) EdU+ myoblasts, identified by Syndecan-4 immunoreactivity quantified under growth conditions. H) Nuclei in myosin+ cells quantified under growth conditions. I) Schematic of stress recovery experimental design. J) EdU+ myoblasts, identified by Syndecan-4 immunoreactivity quantified under growth conditions following stress from serum withdrawal. K) Schematic of serum withdrawal experimental design. L) Nuclei in myosin+ cells quantified under differentiation conditions. For G-L, n=4 primary myoblast isolates, comparisons by t-tests.

If caFGFR1 signaling replaces FGF2, then recombined MuSCs should be unresponsive to added FGF2. MuSCs explanted from tamoxifen treated caFGFR1+ and caFGFR1− mice were cultured in the presence and absence of doxycycline and with or without FGF2, followed by EdU addition 2 hours prior to fixation. In cultures without FGF2, only 26% of vehicle treated caFGFR1+ PAX7+ cells were EdU+, but ∼53% were EdU+ with doxycycline treatment. With FGF2 addition, ∼70% of PAX7+ cells were EdU+ in both vehicle and doxycycline treated conditions, an average of 2.7-fold increase of caFGFR1− PAX7+ cells and only a 1.3-fold increase of caFGFR1+ PAX7+ cells (Fig. 1E). Thus, caFGFR1 signaling replaces FGF2 addition in recombined cells.

MuSCs isolated from aged mice are less proliferative, differentiate prematurely, and are less resilient following stress compared to MuSCs isolated from young mice^12,17^. Primary myoblasts isolated from 24-month-old caFGFR1+ mice were cultured in growth media with or without doxycycline, followed by a 2-hour pulse of EdU (Fig. 1F). Proliferating MuSCs increased 3-fold by doxycycline addition, and differentiation was reduced 4-fold compared to MuSCs cultured with vehicle only, demonstrating that caFGFR1 expression rescues age-associated proliferation deficits and prevents age-associated premature differentiation (Fig. 1G-H). In low serum and the presence of FGF2, MuSCs from young mice remain in a quiescent, undifferentiated state, and re-introducing high serum promotes cell cycle re-entry. In contrast, MuSCs from aged mice differentiate in low serum, but this failure of MuSCs from aged mice to enter quiescence in low serum is rescued by FGF2 addition^12,23^.

MuSCs isolated from aged caFGFR1+ mice were cultured in the presence of doxycycline or vehicle control and subjected to serum withdrawal and reintroduction (Fig. 1I). Less than 3% of the vehicle treated mononuclear MuSCs reentered the cell cycle upon serum restoration, while ∼25% of mononuclear doxycycline treated cells recovered from the stress and reentered the cell cycle 24h after reintroducing serum (Fig. 1J). Expression of caFGFR1 does not prevent differentiation in the presence of pro-differentiation signals as caFGFR1+ myoblasts cultured in low serum differentiation media with doxycycline or vehicle (Fig. 1K) are equally capable of differentiating (Fig. 1L). Thus, signaling from caFGFR1 expression rescues cell intrinsic MuSC aging phenotypes.

Aged mice have half of the number of MuSCs compared to young mice. To attempt restoring MuSC numbers in aged mice, we treated aged (20–26-month-old) caFGFR1+ and caFGFR1− mice with tamoxifen for 5 days and fed the mice doxycycline chow for 30 days prior to collection of the tibialis anterior (TA) muscle (Fig. 2A). Cross-sections of TA muscles from aged caFGFR1+ mice possessed 1.7-fold more MuSCs, identified by immunoreactivity to PAX7, than aged caFGFR1− mice (Fig.2B, S1A). Since decreased MuSC self-renewal is a hallmark of MuSCs from aged mice^12^, single myofibers from aged caFGFR1+ and caFGFR1− mice were explanted and assayed for asymmetric division by parD3 distribution (Fig. 2C)^27^. We observed a >2-fold increase in asymmetrically dividing MuSCs on myofibers explanted from aged caFGFR1+ mice compared to aged caFGFR1− mice (Fig. 2D), indicating that caFGFR1 expression rescues age-related declines in asymmetric division.

**Figure 2.**
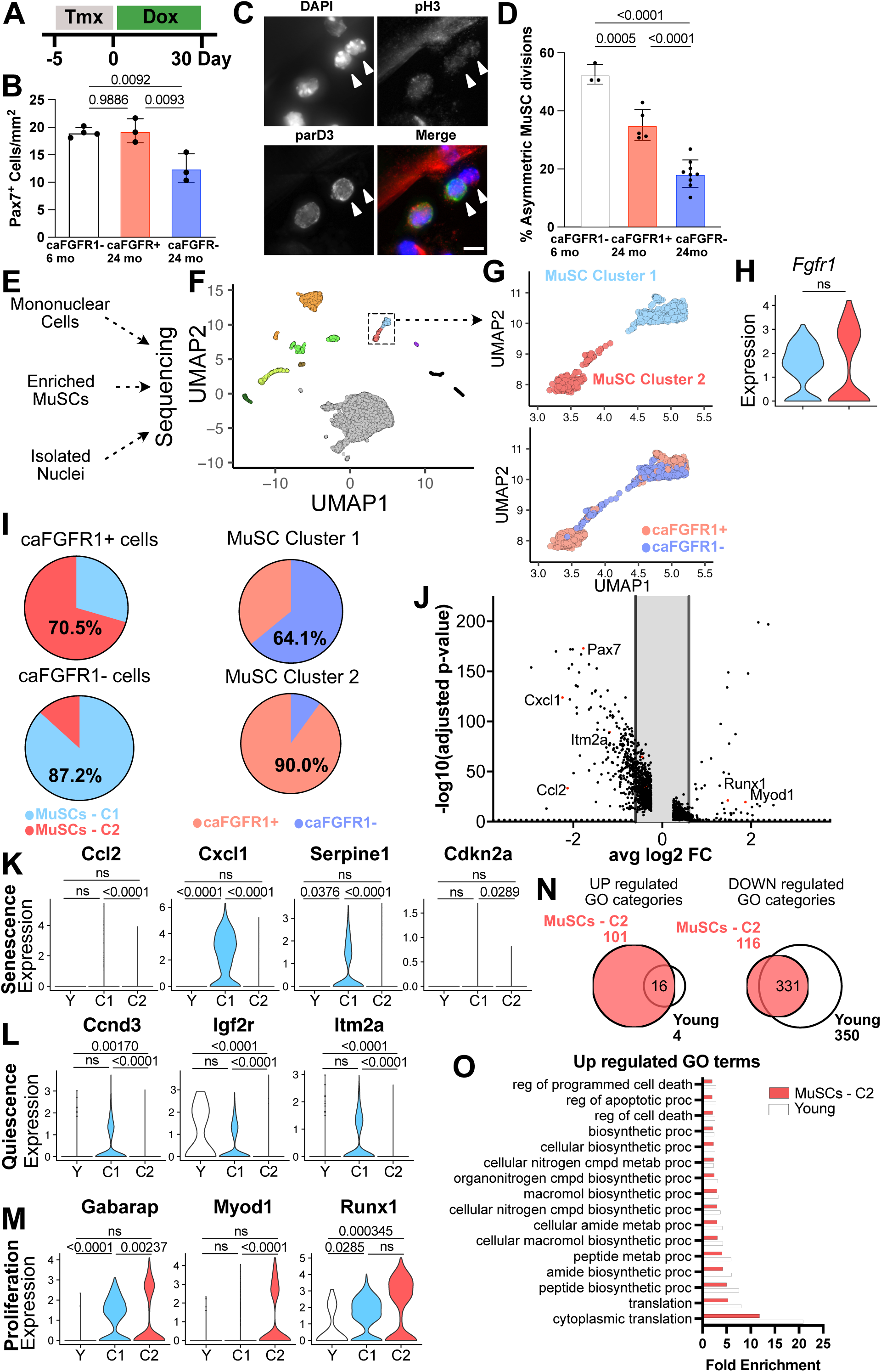
caFGFR1 expression partially rejuvenates MuSCs in old mice. A) Treatment of old mice (20–24-month-old) with tamoxifen for 5 days followed by 30 days of doxycycline chow. B) PAX7+ cells/mm2 quantified from n=3-4 mice per genotype, comparisons by one-way ANOVA. C) Immunofluorescence images of MuSCs on myofibers isolated from aged mice. Phospho-histone 3 (pH3) identifies cells in the cell cycle, and ParD3 is asymmetrically distributed between daughter cells. Arrowheads indicate asymmetric daughter cells. Scale bar = 10 μm. D) The percentage of asymmetric MuSC divisions quantified on isolated myofibers. n=3-9 mice with >100 MuSCs quantified per mouse, comparisons by one-way ANOVA. E) Schematic of single cell and single nuclear sequencing experiments from old caFGFR1+ and caFGFR1−mice. F) UMAP plot of all cells and nuclei from the 3 sequencing experiments. G) Inset of the two MuSC clusters colored by clusters and by genotypes. H) Violin plot of *Fgfr1* transcript levels in MuSC Clusters 1 and 2. I) The percent of MuSCs isolated from caFGFR1+ and caFGFR1− mice in MuSC Clusters 1 and 2 and the percent of the MuSC Clusters 1 and 2 composed of each genotype. J) Volcano plot comparing transcripts between the MuSC Clusters 1 and 2 from the combined data set. Transcripts on the left are depleted from caFGFR1+ MuSCs and transcripts on the right are enriched in caFGFR1+ MuSCs. Transcripts of special note with greater than 1.5-fold change compared to control are marked in red and labeled. Violin plots of transcript levels between young (Y) and MuSC Clusters 1 and 2 for transcripts associated with K) senescence, L) quiescence, and M) proliferation. N) Venn diagrams of GO terms of the up-regulated transcripts and down-regulated transcripts for the comparison of MuSC Cluster 2 (salmon) and young (white) cells. O) The fold enrichment relative to background of the concordant GO categories of up-regulated transcripts.

Expressing caFGFR1 in MuSCs from aged mice rescues cell intrinsic aging phenotypes *in vitro* and *in vivo*, suggesting caFGFR1 expression rejuvenates MuSCs in aged mice. If MuSCs from aged mice are rejuvenated by caFGFR1 expression, then transcriptomes of caFGFR1+ MuSCs would exhibit a more youthful signature than caFGFR1− MuSCs. To assess the transcriptomes of mononuclear cells and myonuclei in skeletal muscle, we employed three distinct sequencing strategies: 1) single cell RNA-sequencing (scRNA-seq) of all mononuclear cells isolated from muscle, 2) scRNA-seq of enriched MuSCs, and 3) single nuclear RNA sequencing (snRNA-seq) optimized to capture myonuclei from myofibers (Fig. 2E, S1B)^24^. Clustering individual cells and nuclei together revealed cell types expected from skeletal muscle, identified by cell-type specific markers (Fig. 2F, S1C-E). Two distinct MuSC clusters were identified, one enriched for cells isolated from caFGFR1+ mice and the other enriched for cells from caFGFR1− mice (Fig. 2G), and express similar levels of Fgfr1 and caFGFR1 transcripts, indicating that the caFGFR1 gene is not overexpressed *in vivo* (Fig. 2H). When quantified, 71% of MuSCs isolated from caFGFR1+ mice segregate into MuSC Cluster 2 (Fig. 2I), consistent with the recombination frequencies of caFGFR1+ cultured myoblasts (Fig. 1D). Comparing the two MuSC clusters identified 127 differentially regulated transcripts (Fig. 2J, Table S1). Transcripts associated with senescence^25,26^ (Fig. 2K, S1F) and quiescence^27^ (Fig. 2L, S1F) were depleted in MuSC Cluster 2 while transcripts associated with MuSC activation^23,28,29^ (Fig. 2M, S1F) were enriched. Thus, MuSC Cluster 2 contains the rejuvenated MuSCs from aged caFGFR1+ mice and any MuSCs from aged caFGFR1− mice that have not adopted an aged phenotype.

To examine the extent to which caFGFR1+ MuSCs were rejuvenated, we incorporated published scRNA-seq datasets of young (3-6 month old) and aged (24 month old) mice^30–32^. Comparing the gene ontology (GO) biological process terms of the differentially expressed transcripts from the MuSC Cluster 2 of both aged caFGFR1+ mice and aged caFGFR1− mice to the GO terms obtained from differentially expressed transcripts between wild-type young (3-6 month old) and aged (24 month old) mice^30–32^ revealed that 16 out of 20 total GO terms upregulated in caFGFR1+ MuSCs were also upregulated in MuSCs from young mice when compared to old mice (Fig. 2N). The most significant GO process categories enriched in caFGFR1+ MuSCs and young mice were cytoplasmic protein translation, protein translation, peptide biosynthetic process, as well as altered metabolism and negative regulation of apoptosis (Fig. 2O). A total of 350 GO categories were depleted in MuSCs from young mice when compared to MuSCs from aged mice. Of those, 331 were depleted in caFGFR1+ MuSCs compared to caFGFR1−MuSCs obtained from aged mice (Fig. 2N). The depleted GO categories common across these comparisons are associated with ATP production, response to growth factors, and positive regulation of apoptosis (Fig. S1G). The striking agreement of enriched GO processes (80%) and of depleted GO processes (95%) detected when comparing MuSC Cluster 2 with MuSCs isolated from young and aged mice further demonstrates that caFGFR1 signaling rejuvenates MuSC from aged mice as they acquire transcriptional signatures like MuSCs from young mice.

MuSCs are optimally positioned to affect surrounding cell types, occupying a niche against the myofiber close to capillaries^33^ and enriched at the neuromuscular junction^34^ (NMJ). We queried the sc/snRNA-seq data sets of caFGFR1+ and caFGFR1−mice that were fed doxycycline chow for 30 days and asked whether rejuvenated MuSCs altered the transcriptomes of myonuclei and other mononuclear cells in skeletal muscle (Fig. 3A). Among 730 differentially expressed transcripts between myonuclei isolated from caFGFR1+ and caFGFR1− mice (Fig. 3B, Table S2), 42% of transcripts enriched in caFGFR1+ myonuclei were related to GO processes associated with proteostasis and denervation response (Fig. S2A). Transcripts depleted from caFGFR1+ myonuclei were associated with GO processes linked to metabolism, including differences in electron transport, glycogen biosynthesis, mitochondria, and focal adhesions (Fig. S2B). Because proteostasis-linked transcripts were altered, we asked if MuSCs expressing caFGFR1 induced muscle hypertrophy, but found no differences in TA muscle wet weight (Fig. 3C), myofiber number (Fig. 3D), or myofiber diameter (Fig. 3E, F) between aged caFGFR1+ and caFGFR1− mice. Increased MuSC contribution to NMJs improves innervation in old mice^34^, thus increased MuSC content of caFGFR1+ muscles may improve myofiber innervation. We isolated myofibers from caFGFR1+ and caFGFR1− mice and evaluated NMJ complexity (Fig. 3G, S2C) and innervation (Fig. 3H, S2D); neither measure was significantly different, nor was the final number of Pax7 cells at 30 dpi (Fig. S2E).

**Figure 3.**
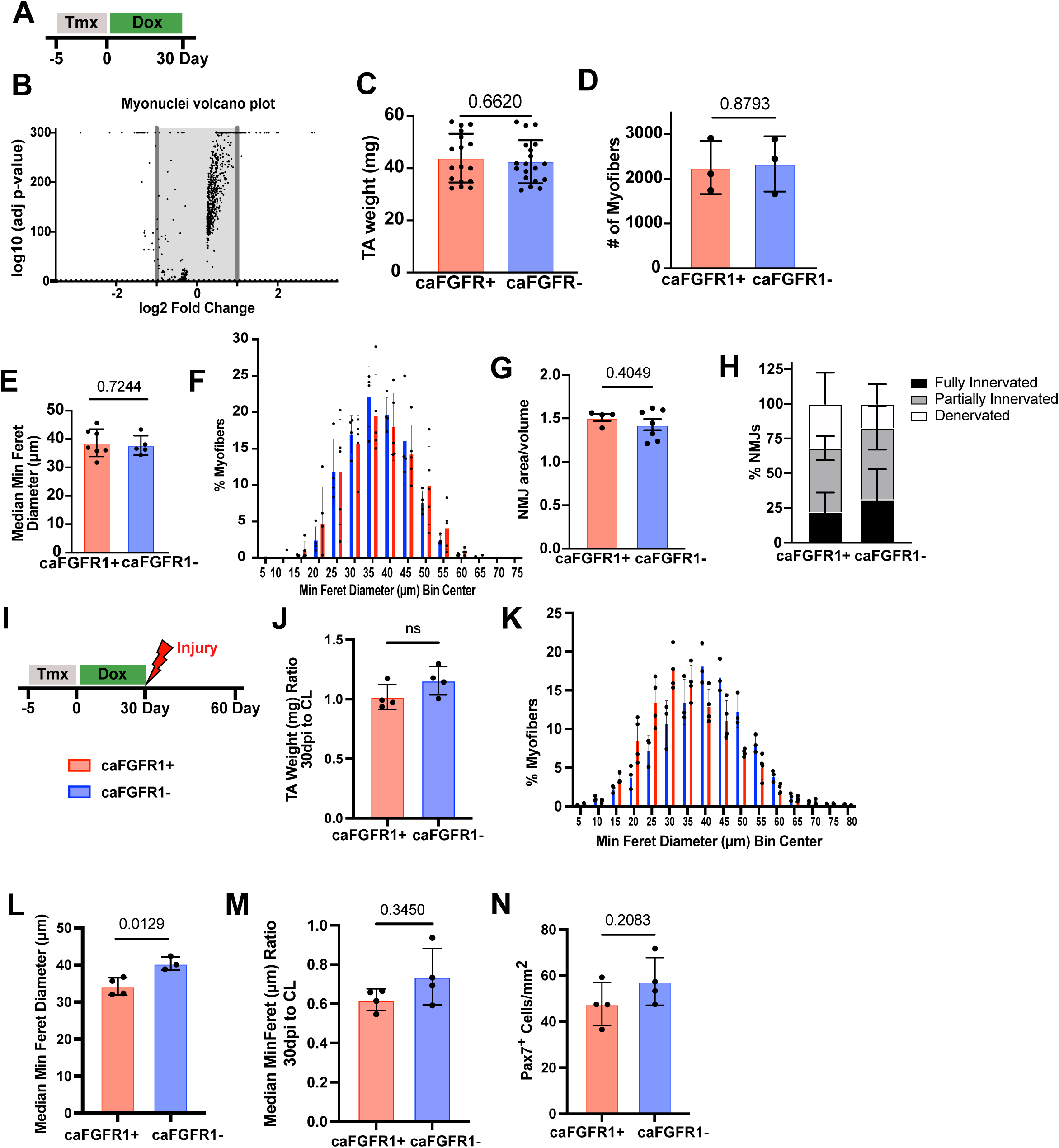
Rejuvenating MuSCs does not improve muscle morphology and regeneration in aged mice. A) Treatment of old mice with tamoxifen for 5 days followed by 30 days of doxycycline. B) Volcano plot comparing myonuclear transcripts from caFGFR1+ and caFGFR1− mice. Transcripts on the left are depleted from myonuclei from caFGFR1+ mice and transcripts on the right are enriched in myonuclei from caFGFR1+ mice. C) TA muscle wet weight from old caFGFR1+ and caFGFR1− mice quantified from n=17-20 mice, comparison by t-test. D) The number of myofibers in a full TA muscle section from caFGFR1+ and caFGFR1− old mice. n=3 mice with 3 full TA sections quantified per mouse, comparison by t-test. E) Distribution of minimum Feret diameters of myofibers in the TA. n=4 mice with >500 myofibers quantified per mouse. F) Median minimum Feret diameter for TA muscles from caFGFR1+ and caFGFR1− old mice. n=5-7 mice, comparison by t-test. G) Neuromuscular junction complexity (represented as Bungarotoxin+ area/endplate volume) on individual myofibers isolated from the extensor digitorum longus muscle of caFGFR1+ and caFGFR1− mice. n=4-7 mice with >100 NMJs quantified per mouse, comparison by t-test. H) NMJ innervation on individual myofibers. n=4-7 mice with >100 NMJs quantified per mouse, comparison by one-way ANOVA. No values were significantly different. I) Schematic of muscle injury following 5 days of tamoxifen and 30 days of doxycycline treatment with regeneration in the absence of doxycycline. J) Ratio of 30 dpi TA muscle wet weight compared to contralateral (CL) from old caFGFR1+ and caFGFR1− mice quantified from n=4 mice, comparison by t-test. K) Distribution of minimum Feret diameters of regenerated myofibers. L) The median minimum Feret diameter of regenerated myofibers. n=3-4 mice with >500 myofibers quantified per mouse, comparison by t-test. M) The ratio of median minimum Feret diameters of 30 dpi regenerated myofibers to CL myofibers of the same mouse (n=4), with >500 myofibers quantified per mouse, comparison by t-test. N) The number of Pax7+ MuSC per mm^2^ in regenerated TA muscle. n=4 mice, comparison by t-test.

Aging compromises the ability of skeletal muscle to regenerate. Since caFGFR1 expression partially rejuvenates MuSCs from aged mice, muscle repair may be enhanced when MuSCs express caFGFR1. Therefore, we injured TA muscles by BaCl_2_ injection, maintaining aged caFGFR1+ and caFGFR1− mice on doxycycline chow for 30 days (Fig. S2F), collected the TA muscle, and quantified muscle weights (Fig. S2G), myofiber diameter (Fig. S2H, I), and MuSC content (Fig. S2J, K). The TA muscle weights, diameters, and MuSC numbers were not significantly different between muscles with MuSCs expressing caFGFR1 or MuSCs lacking the transgene. Since caFGFR1 represses myogenesis and might interfere with MuSC progeny differentiating and fusing, aged caFGFR1+ mice and caFGFR1− mice were recombined, fed doxycycline for 30 d, injured with BaCl_2_, doxycycline removed, and assessed at 30 d post-injury (Fig. 3I). Quantifying the normalized TA weights (Fig. 3J), minimum feret diameters (Fig. 3 K, L) and normalizing diameters to contralateral TA muscles (Fig. 3M) revealed no significant differences between aged control groups and those expressing caFGFR1. No differences in MuSC numbers were observed in regenerated muscle, whether MuSCs expressed caFGFR1 for 30d prior to injury (Fig. 3N). Since the myofiber diameters and TA muscle weights vary significantly between aged mice due to variability in age-related phenotypes, normalizing data from injured TA muscle to the contralateral limb TA muscle of the same mice reduces variability and provides a more accurate assessment of interventional effects.

Rejuvenating SCs failed to improve muscle aging phenotypes in the absence of an induced injury, failed to improve muscle phenotypes following an injury, and failed to improve the regenerative response. Thus, we asked if cell nonautonomous age-related changes affected the capability of caFGFR1 expressing MuSCs to improve muscle in aged mice. To identify cell nonautonomous signaling pathways in skeletal muscle, we compared signaling pathways between young mice, aged caFGFR1+ mice, and caFGFR1− mice by Seurat dimensional reduction and cell type clustering of sc/snRNAseq data using CellChat network centrality analyses. We identified significant collagen signaling network expression where fibroblasts were the dominant senders. Endothelial cells and smooth muscle cells were the only receivers in young mice, while MuSCs were the receivers in aged mice, regardless of caFGFR1 expression (Fig. 4A). MuSCs in Cluster 1 were dominant receivers, as well as influencers and mediators, while Cluster 2 MuSCs were moderate receivers (Fig. 4A). Syndecan-4, is expressed at low levels in MuSCs from young mice and at higher levels in Cluster 1 from aged mice vs. lower levels in Cluster 2 MuSCs from aged mice (Fig. 4B). Syndecan-4 in Cluster 2 is the receiver for caFGFR1+ MuSCs and syndecan-4 in Cluster 1 is the receiver for caFGFR1− MuSCs (Fig. 4C). The primary collagens expressed by the sender fibroblasts are expressed at much greater levels in TA muscle of aged mice compared to TA muscle in young mice (Fig. 4D, S3A) where ≤40% of the fibroblasts in muscle from young mice express collagens 1, 4 and 6 (Fig. 4E, S3B). In contrast, ≥70 of the fibroblasts in muscle from aged caFGFR1+ or caFGFR1− mice express collagens 1, 4 and 6 (Figs. 4D S3A). Collagen protein levels were assessed in uninjured TA muscles and at 7 dpi in injured skeletal muscle from young mice and aged mice (Fig. 4F). Aged caFGFR1+, aged caFGFR1−, and young mice were injected with tamoxifen and fed doxycycline chow for 30 d. Mice were then injured by BaCl_2_ injection and 7 dpi TA muscles harvested (Fig. 4F). In the uninjured contralateral TA muscle of young mice, low level variable protein immunoreactivity is visible for Col 1, Col6a1, and Col 4 (Fig. 4G, H) while much higher levels are present in muscle from aged caFGFR1+ mice as well as caFGFR1− mice (Fig. 4G, H). The collagen proteins dramatically increased at 7 dpi in young mice but remained relatively constant in aged mice whether MuSCs had expressed caFGFR1 (Fig. 4I, S3C).

**Figure 4.**
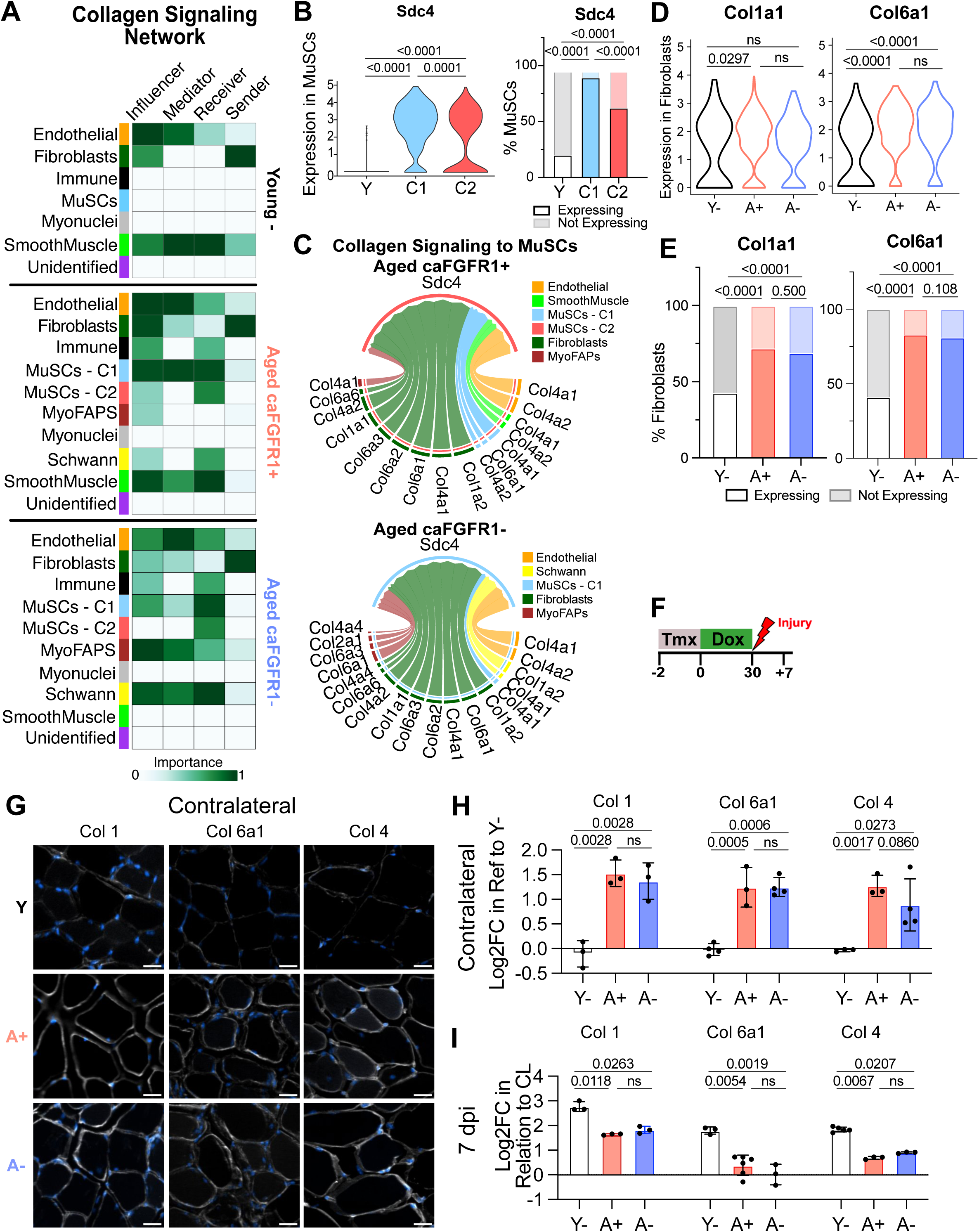
caFGFR1 expression in aged mice does not rescue the age-related elevation of collagen signaling. A) Cell-Chat network centrality analysis of the collagen signaling pathway in young caFGFR1− mice, aged caFGFR1+ mice, and aged caFGFR1− mice. MuSCs are additionally separated by clusters (MuSC Cluster 1 and Cluster 2) in aged caFGFR1+ and caFGFR1− mice. B) Expression of Sdc4 in aged mice compared to young mice, separated by MuSC Clusters 1 and 2. Violin plot of *Sdc4* transcript levels in the MuSC Clusters 1 and 2, comparisons by Seurat adjusted p-values (FindMarkers). Percentage of MuSCs expressing Sdc4, comparisons by contingency Fisher’s exact tests. C) Chord diagram displaying individual cell-type groups that express collagen ligands which may interact with receptors on MuSCs from aged caFGFR1+ and aged caFGFR1− mice. D) Violin plots of transcript levels between young caFGFR1− (Y), aged caFGFR1+ (A+), and aged caFGFR1− (A-) fibroblasts for transcripts associated with collagen. Adjusted p-values from Seurat FindMarkers’ tables. E) Percentage of fibroblasts expressing collagen genes, comparisons by contingency Fisher’s exact tests. F) Schematic of muscle injury following 2 days of tamoxifen and 30 days of doxycycline treatment with regeneration in the absence of doxycycline. G) Immunoreactivity for Col1, Col6a1, and Col4 on cross-sections from the contralateral TA muscle of 7-days post-injury (dpi) young caFGFR1− (Y-), aged caFGFR1+ (A+), and aged caFGFR1− (A-) mice (n=3, scale bar = 20 μm). Collagen immunoreactivity is white while DAPI is blue. H-I) Log 2 fold change (Log2FC) of collagen immunoreactivity shown in G and Fig. S3C in relation to young signal for the contralateral samples and in relation to the contralateral signal for the 7 dpi samples (n=3), comparisons by nested one-way ANOVA.

MuSCs reside in an asymmetric niche interacting with the myofiber plasma membrane and the basement membrane where ECM proteins including collagen, fibronectin, laminin, and heparan sulfate proteoglycans influence MuSC behavior. The MuSC niche regulates MuSC function, is dynamically remodeled following a muscle injury, and is altered in diseased muscle^17,35,36^. Since collagen proteins and Syndecan-4 are elevated in muscle from aged mice and unaffected by MuSCs expressing caFGFR1, we asked if the collagens present in muscle from aged mice influence the ability of MuSCs to proliferate and differentiate. Hindlimb muscle from young mice and aged mice was isolated, decellularized and resuspended for coating tissue culture plates (Fig. 5A). MuSCs isolated from young mice, recombined aged caFGFR1− mice and recombined caFGFR1+ mice were cultured with doxycycline on gelatin as a control, on ECM from young mice (ECM-Y), or aged mice (ECM-A), monitoring cell growth for 3 days (Fig. 5B). MuSCs from young mice and aged mice did not preferentially adhere to gelatin, ECM-Y or ECM-A, but differences in total number of adherent cells between mouse age and genotype groups were apparent at one or two days of culture (Fig. S4). MuSCs were assessed after 3 days of culture and MuSC numbers were normalized to their respective day 1 cultures to compare growth curves. MuSCs from young mice expanded on ECM-Y but failed to expand on ECM-A (Fig. 5B, C), while MuSCs from aged caFGFR1− mice failed to expand on either ECM-Y or ECM-A, consistent with cell autonomous aging deficits (Fig. 5B, C)^10,11,24^. In contrast, MuSCs expressing caFGFR1 expanded on ECM-Y, like MuSCs from young mice, but grew poorly on ECM-A (Fig. 5B, C). Thus, ECM from aged mice inhibits MuSC expansion, demonstrating a cell nonautonomous aging deficit that could affect the capability of rejuvenated MuSCs to improve muscle in aged mice.

**Figure 5.**
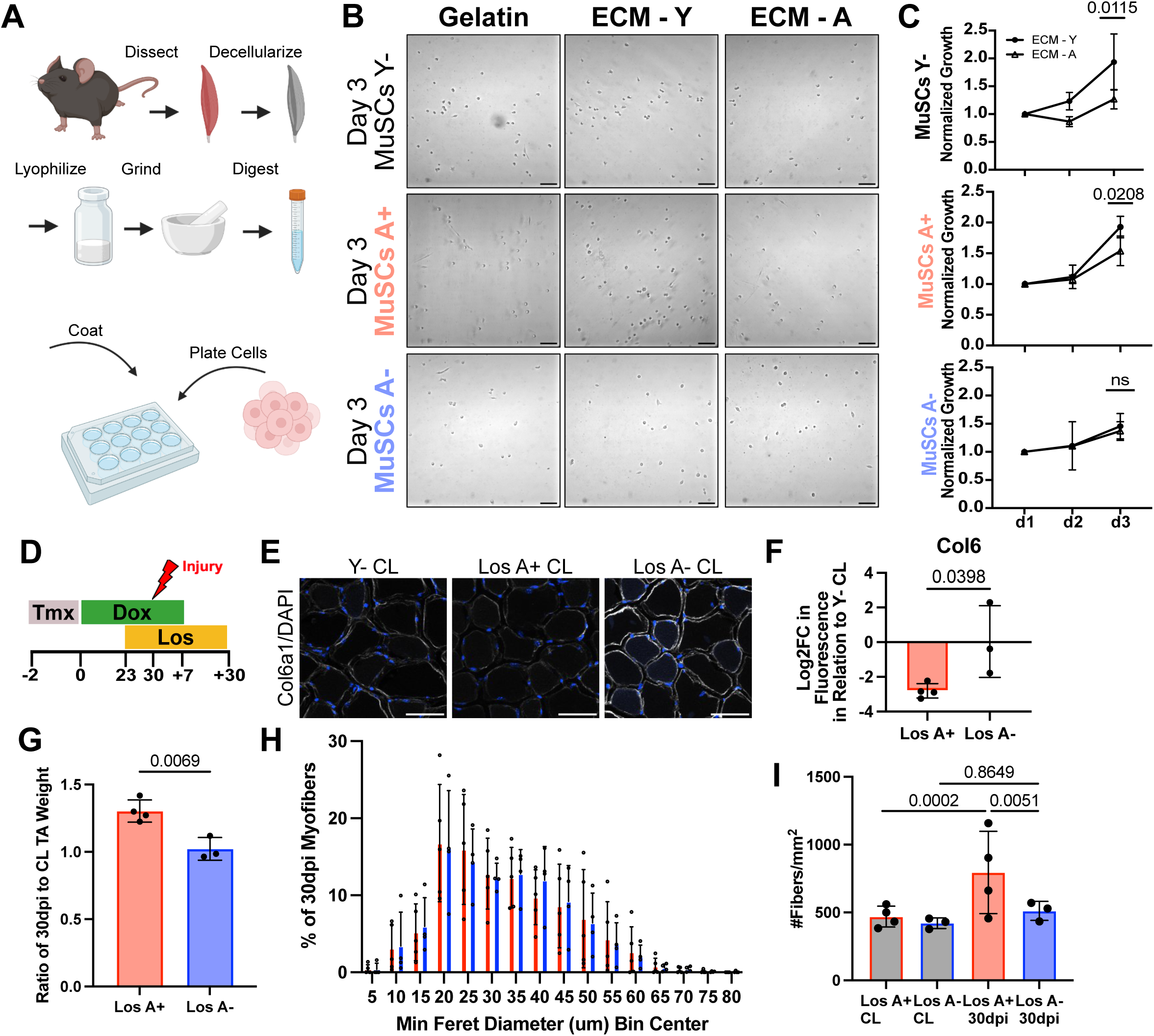
Extracellular matrix from aged mice limits MuSC growth and combinatory anti-fibrotic treatment and caFGFR1 expression improve muscle regeneration in aged mice. A) Schematic of the extracellular matrix (ECM) growth experiment. ECM was extracted from both young and aged wildtype mice, decellularized with 1% SDS, lyophilized, ground to a fine powder, and dissolved in 0.1M HCl. ECM solution was then diluted 1:5 with 0.1M acetic acid and incubated with gelatin-coated plates overnight before MuSCs were plated. B) Brightfield images of Day 3 growth of the Y-, A+, and A+ MuSCs on either gelatin control, ECM components from a young wildtype mouse (ECM-Y), or ECM components from an aged wildtype mouse (ECM-A) (n=3, scale bar = 100 μm). C) Growth curves of MuSCs on ECM components normalized to growth on the gelatin control for three days (d1, d2, d3), comparisons by two-way ANOVA (n=3). D) Schematic of muscle injury following 2 days of tamoxifen, 30 days of doxycycline treatment, and 7 days of Losartan (Los) administration with regeneration in the presence of Losartan and the first 7 days of doxycycline. E) Ratio of 30 dpi TA muscle wet weight compared to CL from old caFGFR1+ and caFGFR1− mice quantified from n=3-4 mice, comparison by t-test. F) Representative images of Col6a1 immunohistochemistry in cross-sections of the contralateral TA muscle from 30 dpi mice. Scale bar = 50 μm. G) Log 2-fold change (Log2FC) of Col6a1 immunoreactivity shown in F in relation to young signal for the contralateral samples (n=3), comparison by t-test. H) Distribution of minimum Feret diameters of regenerated myofibers. I) The number of fibers per mm^2^ of CL and 30 dpi TAs for Losartan-treated, aged caFGFR1+ and caFGFR1− mice (n=3), comparisons by one-way ANOVA.

Since aging disrupts the MuSC niche, in part by altering the ECM^7,37,38^, and anti-fibrotic drugs reduce fibrosis when muscle is injured in aged mice, we asked if treating mice with Losartan would improve the regenerative capacity of MuSCs expressing caFGFR1. Mice possessing the caFGFR1 transgene and caFGFR1− controls were recombined, fed doxycycline chow, and then treated with Losartan prior to an induced BaCl_2_ injury and maintained on Losartan until 30 dpi when the injured and contralateral TA muscles were harvested (Fig. 5D). Collagen 6a1 protein was assessed by immunoreactivity in uninjured (contralateral) TA muscle sections (Fig. 5E). Compared to young caFGFR1− mice, similar levels of Col6a1 protein were observed in Losartan-treated aged caFGFR1− mice, demonstrating that Losartan reduces age-associated fibrosis with the caveat that Col6a1 protein levels in individual mice were highly variable (Fig. 5F). In aged mice when MuSCs express caFGFR1, Losartan treatment reduced Col6a1 levels dramatically below those in young mice and aged caFGFR1− control mice with little variability between mice (Fig. 5F), identifying a synergistic effect on fibrosis. If the age-associated ECM changes limited the ability of caFGFR1+ MuSCs to repair muscle, then injuring TA muscles in Losartan-treated caFGFR1+ aged mice should improve regeneration. Comparing the TA muscle weights of Losartan-treated caFGFR1−mice with caFGFR1+ mice at 30 dpi identifies a significant gain in TA mass (Fig. 5G) with no significant change in myofiber minimum feret diameter (Fig. 5H). The increase in muscle mass arises from a dramatic, nearly 2-fold increase in myofiber number (Fig. 5I), demonstrating that rejuvenated MuSCs were limited by the skeletal muscle environment.

## Discussion

Skeletal muscle progressively loses force, mass, and regenerative capacity with age in part due to deficits in MuSCs^10,11^. MuSCs in aged mice undergo reduced self-renewal and differentiate prematurely or fail to differentiate^11–14^. Among the signaling pathways compromised in MuSCs of aged mice are autophagy^39^, JAK/STAT signaling^14,40,41^, and FGF signaling^12,14,17^. Activating an ectopic, drug inducible FGFR1 to re-establish FGF signaling rescued aging associated defects in cultured MuSCs and restored the transplantability of MuSCs from aged mice^12^. Furthermore, rescuing asymmetric division in MuSCs from a mouse model of Duchenne Muscular Dystrophy rescues disease phenotypes in MuSCs as well as muscle histology and function^42^. We thus predicted that ectopic FGFR1 signaling, which also rescues asymmetric division^12^, would partially rescue phenotypes attributed to skeletal muscle aging. We constructed mice to regulate spatial and temporal expression of caFGFR1 in MuSCs to test whether forced FGFR1 signaling could rescue some or all the muscle aging phenotypes *in vivo*. MuSCs expressing FGFR1 for 30d appear rejuvenated as MuSC numbers are restored to those of young mice, asymmetric division is restored, stress resistance is improved, and premature differentiation is prevented. Transcriptomic changes between MuSCs from aged caFGFR1+ and caFGFR1− mice compared MuSC transcriptome differences between young and aged mice significantly overlap, providing further evidence that MuSCs from aged mice expressing caFGFR1 appear rejuvenated. Reduced senescence gene expression in caFGFR1+ MuSCs are also consistent with a rejuvenated phenotype. However, we detected no changes in aging-associated muscle phenotypes or improvements in muscle repair upon rejuvenating MuSCs by caFGFR1 expression.

Accompanying the MuSC deficits are changes in the extracellular environment, which appears perpetually inflamed^7,43^ and contains ^7,43^ accumulated collagen, perturbing the MuSC niche^8,37^ and affecting^8,37^ the mechanical properties of the ECM^44^. The inflamed environment leads to accumulating muscle fibrosis^7,45^, compromising the MuSC niche, MuSC quiescence and ligand sensitivity^9,38^. The aging phenotypes in skeletal muscle ECM are thus likely to interfere with the ability of MuSCs to regenerate and maintain skeletal muscle tissue. If MuSCs can be partially rejuvenated, as our data demonstrate and based on several independent observations, the skeletal muscle environment may limit MuSC function via cell nonautonomous mechanisms. Losartan, an inhibitor of the Renin-Angiotensin-Aldosterone System^46^, effectively reduces collagen I deposition and fibrils in tumors^47^ and improves the MuSC niche by reducing fibrosis following a muscle injury^48^. Injuring TA muscles of young mice transiently increased collagen transcripts and collagen protein by 7 dpi but failed to increase collagen levels in aged mice as collagen transcript and protein levels in aged caFGFR1+ mice and caFGFR1− mice are already high in uninjured muscle. In aged caFGFR1+ and caFGFR1− mice, Col I levels increase moderately by 7 dpi with little change in Col 6a1 and Col 4a1 that are indistinguishable whether MuSCs express caFGFR1.

Collagen VI in skeletal muscle, secreted predominately from the fibroblasts, regulates TGF-ß bioavailability in skeletal muscle^48,49^. Collagen VI mutants cause progressive muscular dystrophies^50^, identifying Col VI as a critical regulator of skeletal muscle. Like Col I and Col4a1, Col6a1 transcripts and protein are elevated in caFGFR1+ and caFGFR1− muscle from aged mice. Although administering Losartan reduced average Col6a1 levels in caFGFR1− mice, individual mice exhibited wide variability in Col6a1 levels. In contrast, in mice expressing caFGFR1 in MuSCs Col6a1 is reduced by over 4-fold in uninjured TA muscles with Losartan treatment, compared to caFGFR1− mice, suggesting that the rejuvenated MuSCs and Losartan act synergistically. Following an injury, the TA muscle mass in caFGFR1+ mice was significantly greater than TA muscle mass in injured Losartan-treated caFGFR1− mice, demonstrating elevated collagen levels are primarily responsible for the cell nonautonomous age-related effects limiting muscle repair by caFGFR1 rejuvenated MuSCs. Surprisingly, the increase in muscle mass at 30 dpi is not arising from hypertrophy of caFGFR1+ mouse muscle as the minimum feret diameter of myofibers was unchanged in Losartan-treated caFGFR1+ mice compared to caFGFR1− mice. Instead, the increase in muscle mass occurred from hyperplasia, consistent with a nearly 2-fold increase in myofiber number. Thus, combinatorial treatment to reduce fibrosis and rejuvenate MusCs dramatically improved muscle repair in aged mice, yielding muscle mass and myofiber numbers similar to those in young mice.

MuSCs maintain and repair muscle throughout life, making them an attractive target for gene or cell-based therapies intended to improve muscle health in aging and disease. Expressing caFGFR1 in MuSCs rejuvenates them by restoring a more youthful transcriptome, rescuing cell-intrinsic age associated deficits, restoring asymmetric division, restoring MuSC numbers to youthful levels, and reducing senescence associated transcript levels. Suprisingly, the rejuvenated MuSCs elicited no detectable changes in skeletal muscle phenotypes of aged mice. Targeting a cell population that comprises only 2-7% of nuclei in adult skeletal muscle^51^ and contributes to myofibers at notably low levels in the absence of injury^52^ appears insufficient to improve overall age-associated muscle phenotypes without additional experimental interventions. We demonstrate that reducing fibrosis is one effective intervention. Other interventions that reduce muscle aging phenotypes also involve broad systemic treatments including ablating all senescent cells^53,54^, caloric restriction^55–57^, exercise^58^, or heterochronic parabiosis^16,59–61^. Therefore, developing methods to maintain muscle function and mass during aging will likely require combinatorial interventions that target multiple cell types in the skeletal muscle environment.

## Acknowledgements

We acknowledge the BioFrontiers Sequencing Core, the Light Microscopy Core Facility, and the Stem Cell Research and Technology Resource Center at the University of Colorado Boulder for use of the facilities and expertise in collecting imaging and sequencing data.

## Author Contributions

AAC, TKV, TOV, TLC, KSA, and BBO conceived the experiments. AAC, TKV, TOV, TSZ, NDB, TA, TLC, BP, and CHB performed the experiments, analyzed data, and made Figures. KJL and DMO developed and characterized the caFGFR1 transgenic mice. JK, TSZ, and TKV performed bioinformatic analysis of single cell sequencing. AAC, TKV, and BBO wrote the manuscript. BBO and KSA acquired funding for this study. KSA and BBO supervised the research. All authors read and approved the manuscript.

## Methods Details

### Mice

Mice were bred and housed according to National Institutes of Health (NIH) guidelines for the ethical treatment of animals in a pathogen-free facility at the University of Colorado at Boulder. The University of Colorado Institutional Animal Care and Use Committee (IACUC) approved all animal protocols and procedures. Pax7^iresCre^; ROSA26^LSLrtTA^; caFGFR1 mice were generated by crossing C57Bl/6 (Jackson Labs, ME, USA), Pax7^iresCre^ mice^21^, ROSA26-rtTA mice^20^ (Jackson Labs), and conditional caFGFR1 transgenic mice^18^. Mice from 20-24 months of age were considered old. Initial observations in a small cohort of aged mice suggested a sex difference, consequently, unless otherwise noted experiments were restricted to male mice.

### Tamoxifen and doxycycline treatment

To reduce stress on aged mice, tamoxifen was administered in tamoxifen chow (Envigo TD.130856) for five days. For both age groups, doxycycline was administered in doxycycline chow (Envigo TD.01306) for indicated times.

### Tibialis anterior muscle injuries

Mice were anesthetized with isoflurane, received sustained release Buprenorphine (ZooPharm) and meloxicam as analgesics, and supportive warmed saline, and the left TA muscle was injected with 50μL of 1.2% BaCl_2_. TA muscles were harvested at the indicated time points post injury.

### Losartan Treatment

Pharmaceutical grade Losartan purchased from Millipore Sigma was dissolved at 0.1 g/L in sterile water and fed to mice ad libitum for the indicated durations.

### Primary muscle stem cell isolation

Hindlimb muscles were dissected, minced, and digested in Hams F12 (Gibco) containing 400U/ml collagenase (Worthington) for 60 minutes at 37°C. Digest was quenched with serum containing media and cells filtered through 100µm, 70µm and 40µm. Cells were maintained in Hams F12 media supplemented with Myocult (StemCell Technologies) and 0.8 mM CaCl_2_ at 37°C in 5% CO_2_. Doxycycline was included at 1 µg/ml as experimentally indicated. If required, cells were incubated with 10uM 5-ethynyl-2’-deoxyuridine (EdU – Life Technologies) for two hours prior to fixation. Cells were fixed in 4 % paraformaldehyde for 10 minutes and then processed for immunocytochemistry.

### Primary myofiber isolation

Individual extensor digitorum longus myofibers were isolated, cultured, and immunostained according to previous protocol^62^.^62^. Extensor digitorum longus muscles were dissected and digested in Hams F12 containing 400 U/mL collagenase at 37 °C for 1.5 h. Collagenase was inactivated by the addition serum containing media. Individual myofibers were gently isolated by trituration, maintained in Ham’s F-12 supplemented with 15 % horse serum, 0.8mM CaCl_2_, and 50nM FGF2 at 37°C in 5% CO_2_. Doxycycline was included at 1 µg/ml as indicated. Myofibers were fixed in 4 % paraformaldehyde for 10 minutes and then processed for immunocytochemistry.

### ECM collection and coating

Hindlimb muscles were dissected and chopped into TA muscle-like sizes. The tissue was washed with PBS for 30 minutes at room temperature (RT) and then transferred to 1% (w/v) sodium dodecyl sulfate (SDS) in PBS and rotated for 72 hours at RT, with the solution refreshed every 24 hours. Decellularized skeletal muscle was washed four times in deionized water for 30 minutes at RT to remove the detergent. Following decellularization, the ECM was lyophilized for two days and ground using a mortar and pestle and subsequently digested with pepsin. Pepsin (Sigma, P7012) was dissolved in 0.1 M hydrochloric acid to a concentration of 1 mg/ml, and the ECM was digested in a 10 mg/ml pepsin solution under vortex for 48 hours at RT. The matrix was diluted using 0.1 M acetic acid to obtain a final ECM concentration of 2 mg/ml. These solutions were used to coat tissue culture plates for 1 day at 4°C, followed by rinsing three times with PBS. The coated coverslips were fixed with 4% PFA in PBS for 10 minutes and then washed three times with PBS until further use.

### Immunohistochemistry

TA muscles were dissected and either immediately embedded in O.C.T. (Tissue-Tek) and flash frozen in liquid nitrogen (for fiber type staining) or fixed on ice for 2hrs with 4% paraformaldehyde, and then transferred to PBS with 30% sucrose at 4°C overnight. Muscle was mounted in O.C.T. (Tissue-Tek) and cryo-sectioning was performed on a Leica cryostat to generate 10μm thick sections. Tissue sections were post-fixed in 4% paraformaldehyde for 10 minutes at room temperature (RT) and washed three times for 5 min in PBS. Immunostaining with anti-Pax7, anti-Laminin antibodies required heat-induced epitope retrieval where post-fixed slides were placed in citrate buffer, pH 6.0, and subjected to 6 min of high pressure-cooking in a Cuisinart model CPC-600 pressure cooker. All other antibodies did not require antigen retrieval and so proceeded directly to immunostaining. For immunostaining, tissue sections were permeabilized with 0.25% Triton-X100 (Sigma) in PBS containing 2% bovine serum albumin (Sigma) for 60 min at RT. Incubation with primary antibody occurred at 4°C overnight followed by incubation with secondary antibody at room temperature (RT) for 1hr. Primary antibodies included mouse anti-Pax7 (Developmental Studies Hybridoma Bank, University of Iowa, USA) at 1:750, rabbit anti-laminin (Sigma-Aldrich) at 1:250, rabbit anti-myc-tag (CellSignaling) at 1:200, mouse anti-eMHC (Developmental Studies Hybridoma Bank, University of Iowa, USA) at 1:5, and fiber type antibodies clones SC-71, BF-F3, BA-D5-s (Developmental Studies Hybridoma Bank, University of Iowa, USA). Alexa secondary antibodies (Molecular Probes) were used at a 1:1000 dilution. For analysis that included EdU detection, EdU staining was completed prior to antibody staining using the Click-iT EdU Alexa fluor 488 detection kit (Molecular Probes) following manufacturer protocols. Sections were incubated with 1 μg/mL DAPI for 10 min at room temperature then mounted in Mowiol supplemented with DABCO (Sigma-Aldrich) or ProLong Gold (Thermo) as an anti-fade agent.

### Immunocytochemistry

Fixed cells or fibers were then permeabilized with 0.25% Triton-X100 (Sigma) in PBS containing 2% bovine serum albumin (Sigma) for 60 min at RT. Incubation with primary antibody occurred at 4°C overnight followed by incubation with secondary antibody at room temperature (RT) for 1hr. When staining for alpha-bungarotoxin, fixed myofibers were permeabilized with 0.25% Triton-X100 (Sigma) in PBS containing 2% goat serum (Sigma) for 60 min at RT. Primary antibody incubation was preceded by incubation with Ready Probes^TM^ Mouse on Mouse IgG Blocking Solution (30x) for 60min at RT following the manufacturer’s protocol. Incubation with primary antibody occurred at 4°C overnight in PBS containing 5% goat serum, followed by incubation with secondary antibody in PBS containing 5% goat serum at RT for 1hr. Primary antibodies included mouse anti-Pax7 (Developmental Studies Hybridoma Bank, University of Iowa, USA) at 1:750, rabbit anti-myc-tag (CellSignaling) at 1:400, chicken anti-syndecan4 (Developmental Studies Hybridoma Bank, University of Iowa, USA) at 1:1000, mouse anti-phosphoFGFR1 (CellSignaling) at 1:100, phospho H3 (Millipore Sigma) at 1:500, myog F5D (abcam) at 1:1, emhc F1.652 (DSHB) at 1:5, anti-Synaptic vesicle glycoprotein 2A (Developmental Studies Hybridoma Bank, University of Iowa, USA), and anti-2H3 neurofilament (Developmental Studies Hybridoma Bank, University of Iowa, USA). Alexa secondary antibodies (Molecular Probes) and alpha-bungarotoxin (Invitrogen, Alexa-Fluor 594nm) were used at a 1:1000 dilution. For analysis that included EdU detection, EdU staining was completed prior to antibody staining using the Click-iT EdU Alexa fluor 488 detection kit (Molecular Probes) following manufacturer protocols. Cells were incubated with 1 μg/mL DAPI for 10 min at room temperature then mounted in Mowiol supplemented with DABCO (Sigma-Aldrich) or ProLong Gold (Thermo) as an anti-fade agent.

### Microscopy and image analyses

Images were captured on a Nikon inverted spinning disk confocal microscope or an inverted fluorescence scanner microscope (Olympus IX83). Objectives used on the Nikon were: 10x/o.45NA Plan Apo, 20x/0.75NA Plan Apo and 40x/0.95 Plan Apo. Confocal stacks were projected as maximum intensity images for each channel and merged into a single image. Brightness and contrast were adjusted for the entire image as necessary. Both muscle stem cell numbers and average myofiber diameter were counted manually using Fiji ImageJ. Images were processed using Fiji ImageJ and the analysis package myosoft^63^. Bungarotoxin area was determined using Fiji ImageJ where maximum intensity projections were converted into 8-bit images, the bungarotoxin channel extracted, threshold adjusted, and a binary mask created to determine bungarotoxin area^64^.

### Primers for caFGFR1 detection by PCR

FGFR3 TMD – FGFR1 TKD spanning primers:
FR3-FR1-FW: AGCTACGGGGTGGTCTTCTT
FR3-FR1-RV: AACCAGGAGAACCCCAGAGT

### Software Packages

- Cellranger Software Suite/3.0.1
- FastQC 0.11.8
- R 4.1.1
- Seurat 4.1.1
- SoupX 1.5.2

### Quality control, read alignment, and expression quantification

To assess the quality of Fastq files from sequencing, FastQC was used, evaluating depth and quality of each replicate. Cellranger was then used to process Fastq files and aggregate technical replicates, creating gene-count matrices for each sequencing experiment. For transcriptome alignment of the snRNA-seq datasets, a custom pre-mRNA mm10 reference package was used as previously described for nuclei^22,65^. For the scRNA-seq experiments, the reference package recommended by Cellranger was used.

### Accounting for experimental noise and doublets

Cellranger-aligned feature matrices were then loaded into R using the load10X() function from SoupX. Ambient RNA contamination was predicted using default settings of autoEstCont() and counts were normalized to an estimated noise parameter using adjustcounts().In Seurat, doublets and debris were removed as per their recommended metrics. Cells or nuclei expressing greater than 2,500 or less than 200 features were removed.

### Normalization, dimensional reduction, and nuclear clustering

The tutorial provided by the Mayaan lab was used to guide data normalization (satijalab.org/seurat/articles/pbmc3k_tutorial). Mitochondria and low-quality nuclei were removed. Seurat objects were then passed through NormalizeData(), FindVariableFeatures(), and ScaleData() to scale and log normalize gene counts within each sample. Data were then integrated using the FindIntegrationAnchors() and IntegrateData() functions in Seurat using the reciprocal PCA (rPCA) algorithm.

While performing dimensional reduction of the integrated Seurat object using UMAP and clustering using the shared nearest neighbor (SNN) modularity optimization based clustering algorithm, minimum number of neighbors, minimum distance, and resolution parameters were adjusted to achieve adequate separation of nuclear clusters^66^. Identification of nuclear clusters was done manually, using differential gene expression from the FindMarkers() function between clusters combined with manual literature curation. The Myoatlas database and webtool (research.cchmc.org/myoatlas/) was of particular help in identifying clusters of nuclei^22^.

### Comparison to Kimmel data

To compare the recombined and non-recombined clusters of MuSCs to existing data, data from Kimmel et al., 2020^30^ were downloaded from GSE143476 and processed identically as described above.

### Differential expression tests

Differential gene expression was assessed by the Wilcoxon Rank Sum Test in the Seurat FindMarkers() function. Genes were assessed as significant with an adjusted p-value < 0.05. GO analyses identifying differentially expressed genes were conducted using Panther ^67,68^. A background gene set was used comprising all genes expressed in myogenic nuclei detected from sequencing. GO Biological Processes Complete hierarchy was used to organize results. Example categories in Figures were chosen as contained within ranked clusters. Cell-cell communication analysis was performed on Seurat clusters using the CellChat workflow (Jin et al 2021). ComputeCommunProb() and was used with default trimean to identify significant expression of ligands and receptors across cell type groups.

## REFERENCES

1. Narici, M. V. & Maffulli, N. Sarcopenia: characteristics, mechanisms and functional significance. Br. Med. Bull. 95, 139–159 (2010).

2. Freemont, A. J. & Hoyland, J. A. Morphology, mechanisms and pathology of musculoskeletal ageing. J Pathol 211, 252–9 (2007).

3. Ryall, J. G., Schertzer, J. D. & Lynch, G. S. Cellular and molecular mechanisms underlying age-related skeletal muscle wasting and weakness. Biogerontology 9, 213–28 (2008).

4. Thompson, L. V. Age-related muscle dysfunction. Exp Gerontol 44, 106–11 (2009).

5. Miljkovic, N., Lim, J.-Y., Miljkovic, I. & Frontera, W. R. Aging of Skeletal Muscle Fibers. Ann. Rehabil. Med. 39, 155–162 (2015).

6. Martini, E. M., Garrett, N., Lindquist, T. & Isham, G. J. The Boomers Are Coming: A Total Cost of Care Model of the Impact of Population Aging on Health Care Costs in the United States by Major Practice Category. Health Serv. Res. 42, 201–218 (2007).

7. Mann, C. J. et al. Aberrant repair and fibrosis development in skeletal muscle. Skelet. Muscle 1, 21 (2011).

8. Kanazawa, Y., Miyachi, R., Higuchi, T. & Sato, H. Effects of Aging on Collagen in the Skeletal Muscle of Mice. Int. J. Mol. Sci. 24, 13121 (2023).

9. Wood, L. K. et al. Intrinsic stiffness of extracellular matrix increases with age in skeletal muscles of mice. J. Appl. Physiol. 117, 363–369 (2014).

10. Cosgrove, B. D. et al. Rejuvenation of the muscle stem cell population restores strength to injured aged muscles. Nat. Med. 20, 255–264 (2014).

11. Sousa-Victor, P. et al. Geriatric muscle stem cells switch reversible quiescence into senescence. Nature 506, 316–21 (2014).

12. Bernet, J. D. et al. p38 MAPK signaling underlies a cell-autonomous loss of stem cell self-renewal in skeletal muscle of aged mice. Nat Med 20, 265–71 (2014).

13. Sousa-Victor, P., García-Prat, L., Serrano, A. L., Perdiguero, E. & Muñoz-Cánoves, P. Muscle stem cell aging: regulation and rejuvenation. Trends Endocrinol Metab 10.1016/j.tem.2015.03.006 (2015) doi:10.1016/j.tem.2015.03.006.

14. Chakkalakal, J. V., Jones, K. M., Basson, M. A. & Brack, A. S. The aged niche disrupts muscle stem cell quiescence. Nature 10.1038/nature11438 (2012) doi:10.1038/nature11438.

15. Conboy, I. M., Conboy, M. J., Smythe, G. M. & Rando, T. A. Notch-mediated restoration of regenerative potential to aged muscle. Science 302, 1575–7 (2003).

16. Conboy, I. M. et al. Rejuvenation of aged progenitor cells by exposure to a young systemic environment. Nature 433, 760–4 (2005).

17. Rozo, M., Li, L. & Fan, C.-M. Targeting β1-integrin signaling enhances regeneration in aged and dystrophic muscle in mice. Nat Med 22, 889–96 (2016).

18. Cilvik, S. N. et al. Fibroblast growth factor receptor 1 signaling in adult cardiomyocytes increases contractility and results in a hypertrophic cardiomyopathy. PLoS One 8, e82979 (2013).

19. Naski, M. C., Wang, Q., Xu, J. & Ornitz, D. M. Graded activation of fibroblast growth factor receptor 3 by mutations causing achondroplasia and thanatophoric dysplasia. Nat Genet 13, 233–7 (1996).

20. Belteki, G. et al. Conditional and inducible transgene expression in mice through the combinatorial use of Cre-mediated recombination and tetracycline induction. Nucleic Acids Res. 33, e51 (2005).

21. Murphy, M. M., Lawson, J. A., Mathew, S. J., Hutcheson, D. A. & Kardon, G. Satellite cells, connective tissue fibroblasts and their interactions are crucial for muscle regeneration. Development 138, 3625–37 (2011).

22. Petrany, M. J. et al. Single-nucleus RNA-seq identifies transcriptional heterogeneity in multinucleated skeletal myofibers. Nat. Commun. 11, 6374 (2020).

23. Jones, N. C. et al. The p38alpha/beta MAPK functions as a molecular switch to activate the quiescent satellite cell. J Cell Biol 169, 105–16 (2005).

24. Kurland, J. V. et al. Aging disrupts gene expression timing during muscle regeneration. Stem Cell Rep. 18, 1325–1339 (2023).

25. Saul, D. et al. A new gene set identifies senescent cells and predicts senescence-associated pathways across tissues. Nat. Commun. 13, 4827 (2022).

26. González-Gualda, E., Baker, A. G., Fruk, L. & Muñoz-Espín, D. A guide to assessing cellular senescence in vitro and in vivo. FEBS J. 288, 56–80 (2021).

27. Cheung, T. H. & Rando, T. A. Molecular regulation of stem cell quiescence. Nat. Rev. Mol. Cell Biol. 14, 10.1038/nrm3591 (2013).

28. Cutler, A. A., et al. The regenerating skeletal muscle niche guides muscle stem cell self-renewal. bioRxiv 635805 (2021) doi:10.1101/635805.

29. Umansky, K. B. et al. Runx1 Transcription Factor Is Required for Myoblasts Proliferation during Muscle Regeneration. PLoS Genet. 11, (2015).

30. Kimmel, J. C. et al. Murine single-cell RNA-seq reveals cell-identity- and tissue-specific trajectories of aging. Genome Res. 29, 2088–2103 (2019).

31. Mi, H., Muruganujan, A. & Thomas, P. D. PANTHER in 2013: modeling the evolution of gene function, and other gene attributes, in the context of phylogenetic trees. Nucleic Acids Res. 41, D377–386 (2013).

32. Thomas, P. D. et al. PANTHER: Making genome-scale phylogenetics accessible to all. Protein Sci. 31, 8–22 (2022).

33. Verma, M. et al. Muscle Satellite Cell Cross-Talk with a Vascular Niche Maintains Quiescence via VEGF and Notch Signaling. Cell Stem Cell 23, 530–543.e9 (2018).

34. Liu, W. et al. Loss of adult skeletal muscle stem cells drives age-related neuromuscular junction degeneration. eLife 6, e26464 (2017).

35. Brack, A. S. & Rando, T. A. Intrinsic changes and extrinsic influences of myogenic stem cell function during aging. Stem Cell Rev. Rep. 3, 226–237 (2007).

36. Dumont, N. A. et al. Dystrophin expression in muscle stem cells regulates their polarity and asymmetric division. Nat. Med. 21, 1455–1463 (2015).

37. Olson, L. C., Redden, J. T., Schwartz, Z., Cohen, D. J. & McClure, M. J. Advanced Glycation End-Products in Skeletal Muscle Aging. Bioengineering 8, 168 (2021).

38. Chang, T.-L., Borelli, A. N., Cutler, A. A., Olwin, B. B. & Anseth, K. S. Myofibers cultured in viscoelastic hydrogels reveal the effects of integrin-binding and mechanosensing on muscle satellite cells. Acta Biomater. 192, 48–60 (2025).

39. White, J. P. et al. The AMPK/p27 Kip1 Axis Regulates Autophagy/Apoptosis Decisions in Aged Skeletal Muscle Stem Cells. Stem Cell Rep. 10.1016/j.stemcr.2018.06.014 (2018) doi:10.1016/j.stemcr.2018.06.014.

40. Price, F. D. et al. Inhibition of JAK-STAT signaling stimulates adult satellite cell function. Nat Med 20, 1174–81 (2014).

41. Tierney, M. T. et al. STAT3 signaling controls satellite cell expansion and skeletal muscle repair. Nat. Med. 20, 1182–1186 (2014).

42. Wang, Y. X. et al. EGFR-Aurka Signaling Rescues Polarity and Regeneration Defects in Dystrophin-Deficient Muscle Stem Cells by Increasing Asymmetric Divisions. Cell Stem Cell 0, (2019).

43. The Muscle Stem Cell Niche in Health and Disease. in Current Topics in Developmental Biology vol. 126 23–65 (Academic Press, 2018).

44. The Muscle Stem Cell Niche in Health and Disease. in Current Topics in Developmental Biology vol. 126 23–65 (Academic Press, 2018).

45. Cellular and Molecular Mechanisms Regulating Fibrosis in Skeletal Muscle Repair and Disease. in Current Topics in Developmental Biology vol. 96 167–201 (Academic Press, 2011).

46. Boffa, J.-J. et al. Regression of Renal Vascular and Glomerular Fibrosis: Role of Angiotensin II Receptor Antagonism and Matrix Metalloproteinases. J. Am. Soc. Nephrol. 14, 1132 (2003).

47. Diop-Frimpong, B., Chauhan, V. P., Krane, S., Boucher, Y. & Jain, R. K. Losartan inhibits collagen I synthesis and improves the distribution and efficacy of nanotherapeutics in tumors. Proc. Natl. Acad. Sci. 108, 2909–2914 (2011).

48. Park, J.-K. et al. Losartan Improves Adipose Tissue-Derived Stem Cell Niche by Inhibiting Transforming Growth Factor-β and Fibrosis in Skeletal Muscle Injury. Cell Transplant. 21, 2407–2424 (2012).

49. Mohassel, P. et al. Collagen type VI regulates TGF-**β** bioavailability in skeletal muscle in mice. J. Clin. Invest. 135, (2025).

50. Bönnemann, C. G. The collagen VI-related myopathies: muscle meets its matrix. Nat. Rev. Neurol. 7, 379–390 (2011).

51. Yin, H., Price, F. & Rudnicki, M. A. Satellite cells and the muscle stem cell niche. Physiol. Rev. 93, 23–67 (2013).

52. Keefe, A. C. et al. Muscle stem cells contribute to myofibres in sedentary adult mice. Nat Commun 6, 7087 (2015).

53. Liu, L. et al. Reduction of senescent fibro-adipogenic progenitors in progeria-aged muscle by senolytics rescues the function of muscle stem cells. J. Cachexia Sarcopenia Muscle 13, 3137–3148 (2022).

54. Moiseeva, V. et al. Senescence atlas reveals an aged-like inflamed niche that blunts muscle regeneration. Nature 613, 169–178 (2023).

55. Lee, C.-K., Klopp, R. G., Weindruch, R. & Prolla, T. A. Gene Expression Profile of Aging and Its Retardation by Caloric Restriction. Science 285, 1390–1393 (1999).

56. Cerletti, M., Jang, Y. C., Finley, L. W. S., Haigis, M. C. & Wagers, A. J. Short-term calorie restriction enhances skeletal muscle stem cell function. Cell Stem Cell 10, 515–9 (2012).

57. Mizunoe, Y. et al. Prolonged caloric restriction ameliorates age-related atrophy in slow and fast muscle fibers of rat soleus muscle. Exp. Gerontol. 154, 111519 (2021).

58. Brett, J. O. et al. Exercise rejuvenates quiescent skeletal muscle stem cells in old mice through restoration of Cyclin D1. Nat. Metab. 10.1038/s42255-020-0190-0 (2020) doi:10.1038/s42255-020-0190-0.

59. Brack, A. S. et al. Increased Wnt signaling during aging alters muscle stem cell fate and increases fibrosis. Science 317, 807–10 (2007).

60. Rebo, J. et al. A single heterochronic blood exchange reveals rapid inhibition of multiple tissues by old blood. Nat. Commun. 7, 13363 (2016).

61. Sinha, M. et al. Restoring systemic GDF11 levels reverses age-related dysfunction in mouse skeletal muscle. Science 344, 649–52 (2014).

62. Vogler, T. O., Gadek, K. E., Cadwallader, A. B., Elston, T. L. & Olwin, B. B. Isolation, Culture, Functional Assays, and Immunofluorescence of Myofiber-Associated Satellite Cells. Methods Mol. Biol. Clifton NJ 1460, 141–162 (2016).

63. Encarnacion-Rivera, L., Foltz, S., Hartzell, H. C. & Choo, H. Myosoft: An automated muscle histology analysis tool using machine learning algorithm utilizing FIJI/ImageJ software. PLOS ONE 15, e0229041 (2020).

64. Jones, R. A. et al. NMJ-morph reveals principal components of synaptic morphology influencing structure–function relationships at the neuromuscular junction. Open Biol. 6, 160240 (2016).

65. Cui, M. et al. Dynamic Transcriptional Responses to Injury of Regenerative and Non-regenerative Cardiomyocytes Revealed by Single-Nucleus RNA Sequencing. Dev. Cell S1534580720301465 (2020) doi:10.1016/j.devcel.2020.02.019.

66. Waltman, L. & van Eck, N. J. A smart local moving algorithm for large-scale modularity-based community detection. Eur. Phys. J. B 86, 471 (2013).

67. Ashburner, M. et al. Gene Ontology: tool for the unification of biology. Nat. Genet. 25, 25–29 (2000).

68. The Gene Ontology Consortium et al. The Gene Ontology resource: enriching a GOld mine. Nucleic Acids Res. 49, D325–D334 (2021).

