## Supplemental information for "The Muscle Tissue Environment Limits Muscle Stem Cells in Aged Mice"

Figure S1 Related to Figure 2. A) Immunohistochemistry images of the tibialis anterior of old caFGFR1+ and caFGFR1- mice treated with doxycycline for 30 days. Scale bar = 50 µm. Pax7 white, laminin green, DAPI blue. B) UMAP of pooled results from single cell and single nuclear sequencing experiments with events colored by isolation methods, (C) genotypes, or (D) cell types. E) Violin plots of transcript levels of cell type specific transcripts to identify the cell type of each cluster. F) Violin plots of quiescence associated transcripts in MuSCs. G) The fold-enrichment relative to background of the concordant GO categories of down-regulated transcripts in MuSCs.

Figure S2 Related to Figure 3. The top ten biological process GO terms by fold change of transcripts. (A) significantly enriched in and (B) significantly depleted from caFGFR1+ myonuclei relative to background are presented. C) Representative images of NMJ staining on isolated myofibers and quantification of NMJ complexity. Scale bar = 25 µm. D) Representative images of immunofluorescence for NMJ innervation. Scale bar = 50 µm. E) Representative images of regenerated TA muscles from old caFGFR1+ and caFGFR1- mice pretreated with doxycycline and then with regeneration in the absence of doxycycline. Scale bar = 50 µm. F) Schematic of muscle injury ($BaCl_2$) and regeneration in the presence of doxycycline. G) TA muscle wet weight from old caFGFR1+ and caFGFR1- mice quantified from n=4 mice, comparisons by two-way ANOVA. H) Distribution of minimum Feret diameters of 30 dpi regenerated myofibers in TA muscle of old caFGFR1+ and caFGFR1- mice. n=4 mice with >500 myofibers quantified per mouse. I) Median minimum Feret diameter of regenerated myofibers from caFGFR1+ and caFGFR1- mice. n=4 mice, comparison by t-test. J) Pax7+ MuSC per

$mm^2$ in regenerated TA muscle quantified from caFGFR1+ and caFGFR1- mice. n=4-5 mice, comparison by t-test. K) Representative images of regenerated TA muscles from old caFGFR1+ and caFGFR1- mice with regeneration in the presence of doxycycline.

Figure S3 Related to Figure 4. A) Violin plots of Col1a2, Col4a1, and Col6a2 transcript levels between young caFGFR1- (Y-), aged caFGFR1+ (A+), and aged caFGFR1- (A-) fibroblasts. Adjusted p-values from Seurat FindMarkers' tables. B) Percentage of fibroblasts expressing collagen genes, comparisons by contingency Fisher's exact tests. C) Immunoreactivity for Col1, Col6a1, and Col4 on cross-sections from the injured TA muscle of 7-day post-injury (dpi) young caFGFR1- (Y-), aged caFGFR1+ (A+), and aged caFGFR1- (A-) mice (n=3, scale bar = 20 μm). Collagen immunoreactivity is white while DAPI is blue.

Figure S4 Related to Figure 5. Growth of MuSCs on ECM components from either young (ECM-Y) or aged (ECM-A) caFGFR1- (wildtype) mice. (A) Day 1 and Day 2 growth in presence of doxycycline to accompany Day 3 growth in Fig. 5B and quantification in Fig. 5C (n=3, scale bar = 100 μm). MuSCs were from cell lines established from primary cells isolated from young caFGFR1- (Y-), aged caFGFR1+ (A+), and aged caFGFR1- (A-) mice.

Figure S1

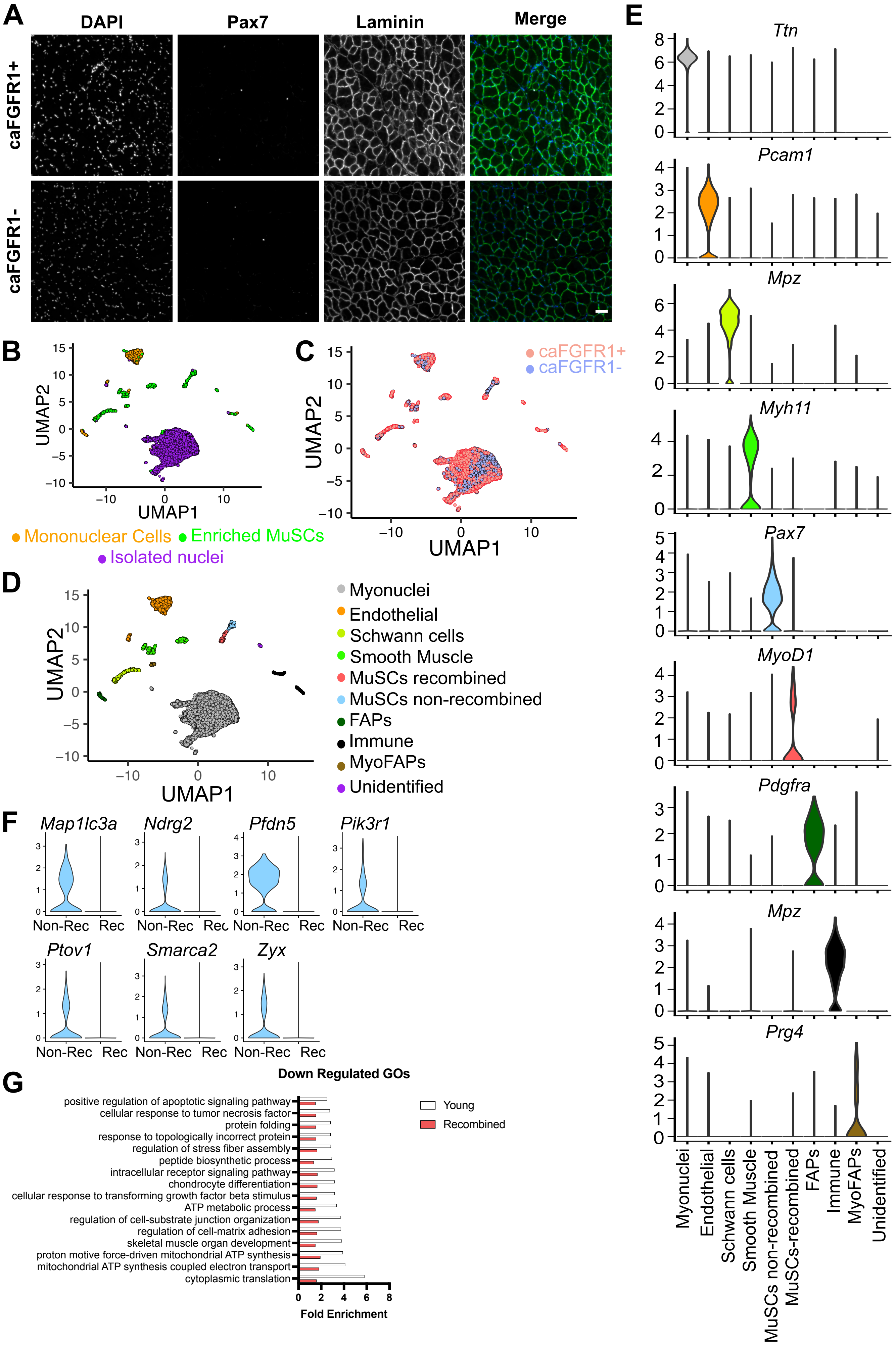

A
DAPI
Pax7
Laminin
Merge
caFGFR1+
caFGFR1-
B
UMAP2
UMAP1
Mononuclear Cells
Enriched MuSCs
Isolated nuclei
C
caFGFR1+
caFGFR1-
UMAP2
UMAP1
D
UMAP2
UMAP1
Myonuclei
Endothelial
Schwann cells
Smooth Muscle
MuSCs recombined
MuSCs non-recombined
FAPs
Immune
MyoFAPs
Unidentified
E
Ttn
Pcam1
Mpz
Myh11
Pax7
MyoD1
Pdgfra
Mpz
Prg4
Myonuclei
Endothelial
Schwann cells
Smooth Muscle
MuSCs non-recombined
MuSCs-recombined
FAPs
Immune
MyoFAPs
Unidentified
F
Map1lc3a
Ndrg2
Pfdn5
Pik3r1
Ptov1
Smarca2
Zyx
Non-Rec
Rec
G
Down Regulated GOs
positive regulation of apoptotic signaling pathway
cellular response to tumor necrosis factor
protein folding
response to topologically incorrect protein
regulation of stress fiber assembly
peptide biosynthetic process
intracellular receptor signaling pathway
chondrocyte differentiation
cellular response to transforming growth factor beta stimulus
ATP metabolic process
regulation of cell-substrate junction organization
regulation of cell-matrix adhesion
skeletal muscle organ development
proton motive force-driven mitochondrial ATP synthesis
mitochondrial ATP synthesis coupled electron transport
cytoplasmic translation
Young
Recombined
Fold Enrichment

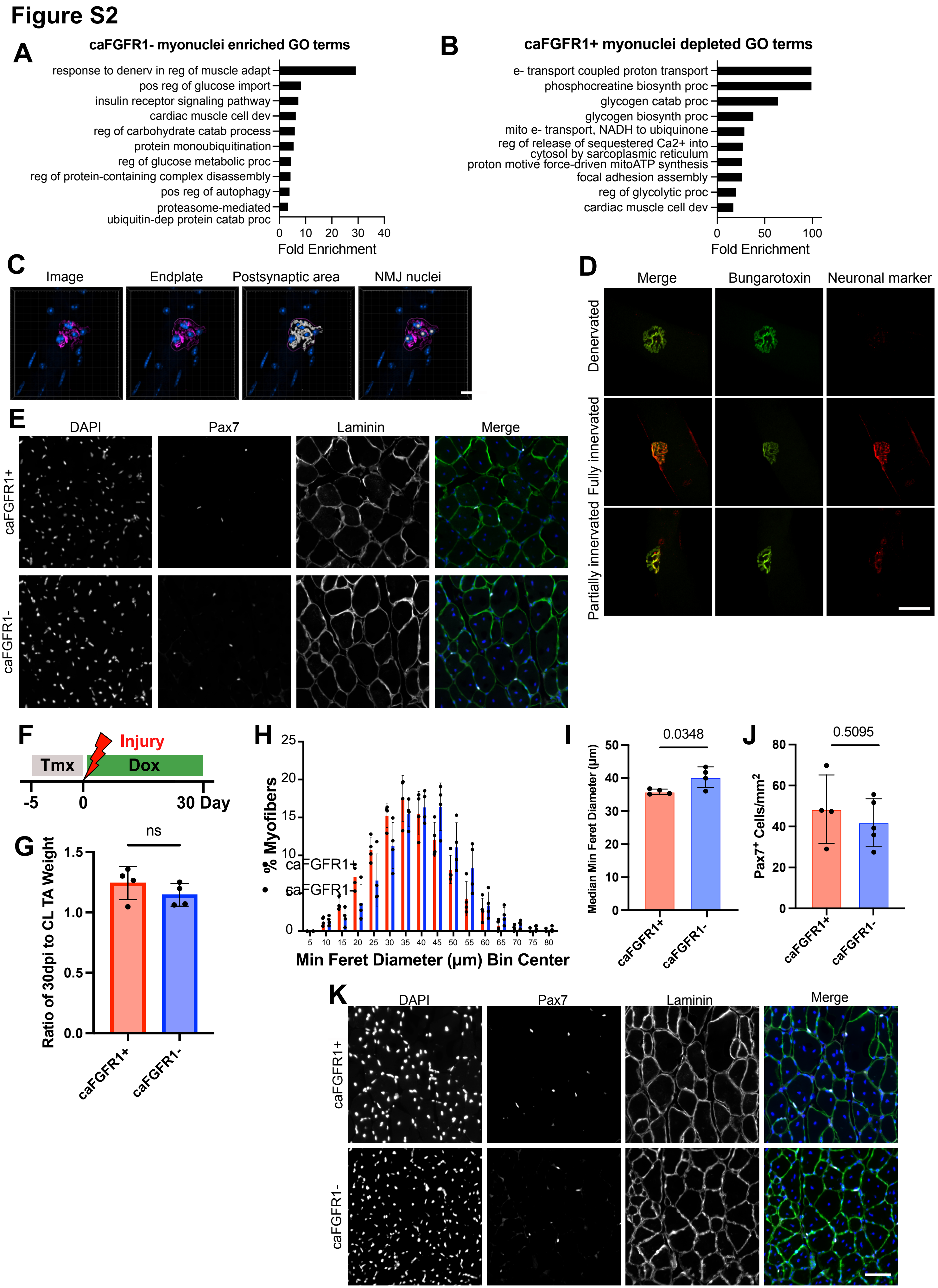
Figure S2
A
caFGFR1- myonuclei enriched GO terms
response to denerv in reg of muscle adapt
pos reg of glucose import
insulin receptor signaling pathway
cardiac muscle cell dev
reg of carbohydrate catab process
protein monoubiquitination
reg of glucose metabolic proc
reg of protein-containing complex disassembly
pos reg of autophagy
proteasome-mediated ubiquitin-dep protein catab proc
0 10 20 30 40
Fold Enrichment
B
caFGFR1+ myonuclei depleted GO terms
e- transport coupled proton transport
phosphocreatine biosynth proc
glycogen catab proc
glycogen biosynth proc
mito e- transport, NADH to ubiquinone
reg of release of sequestered Ca2+ into cytosol by sarcoplasmic reticulum
proton motive force-driven mitoATP synthesis
focal adhesion assembly
reg of glycolytic proc
cardiac muscle cell dev
0 50 100
Fold Enrichment
C
Image
Endplate
Postsynaptic area
NMJ nuclei
D
Merge
Bungarotoxin
Neuronal marker
Denervated
Fully innervated
Partially innervated
E
DAPI
Pax7
Laminin
Merge
caFGFR1+
caFGFR1-
F
Injury
Tmx
Dox
-5
0
30 Day
G
ns
Ratio of 30dpi to CL TA Weight
0.0
0.5
1.0
1.5
caFGFR1+
caFGFR1-
H
% Myofibers
caFGFR1+
caFGFR1-
Min Feret Diameter (μm) Bin Center
I
0.0348
Median Min Feret Diameter (μm)
caFGFR1+
caFGFR1-
J
0.5095
Pax7+ Cells/mm2
caFGFR1+
caFGFR1-
K
DAPI
Pax7
Laminin
Merge
caFGFR1+
caFGFR1-

#### Figure S3

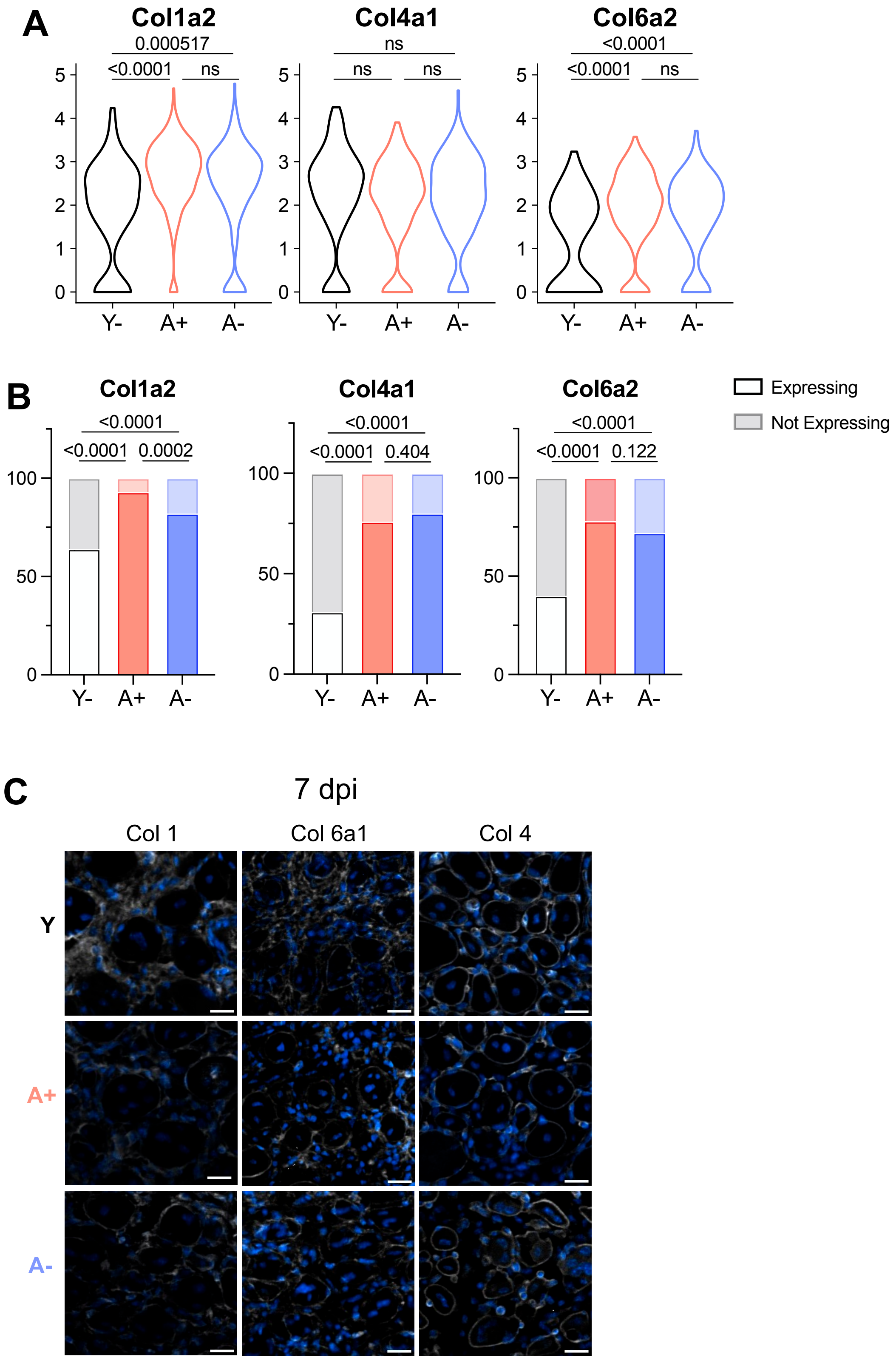

A
Col1a2
0.000517
<0.0001
ns
Col4a1
ns
ns
ns
Col6a2
<0.0001
<0.0001
ns
Y-
A+
A-
B
Col1a2
<0.0001
<0.0001
0.0002
Col4a1
<0.0001
<0.0001
0.404
Col6a2
<0.0001
<0.0001
0.122
Expressing
Not Expressing
C
7 dpi
Col 1
Col 6a1
Col 4
Y
A+
A-

### Figure S4

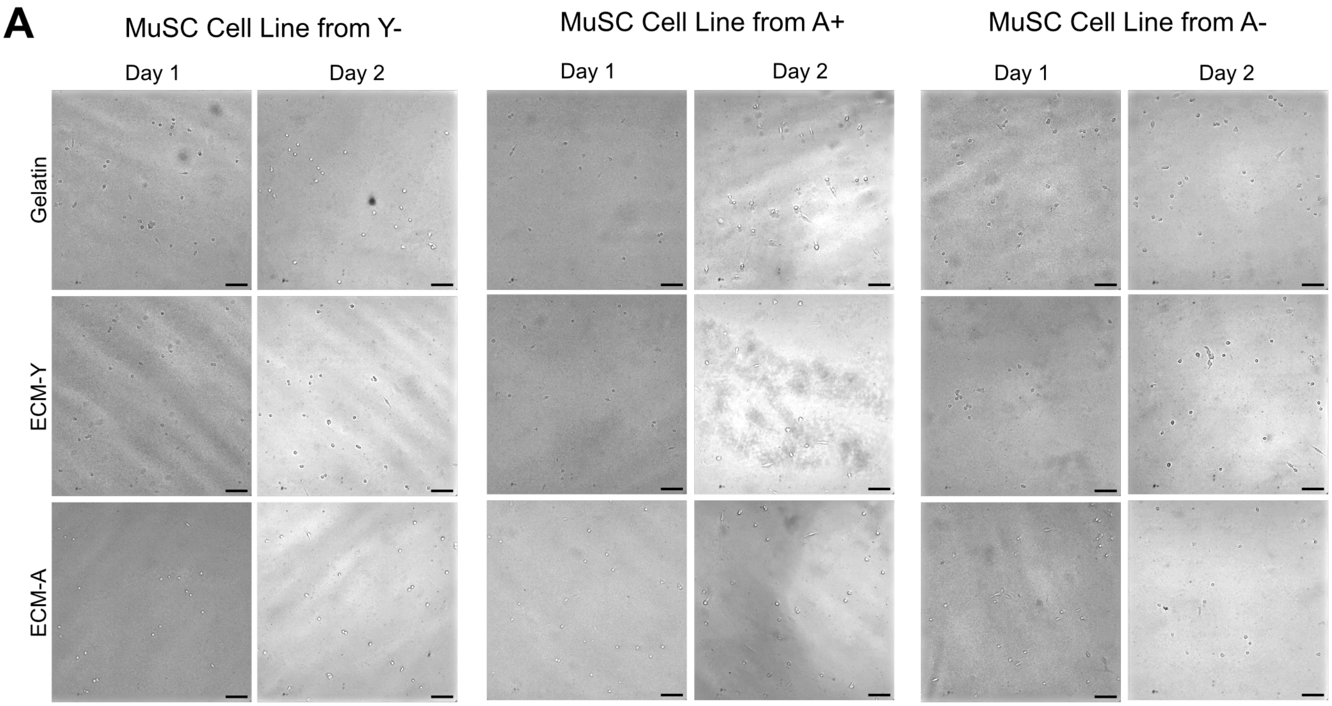
A
MuSC Cell Line from Y-
MuSC Cell Line from A+
MuSC Cell Line from A-
Day 1
Day 2
Day 1
Day 2
Day 1
Day 2
Gelatin
ECM-Y
ECM-A

**Table S1: MuSC differentially expressed transcripts between caFGFR1 recombined and non-recombined clusters**

| | p_val | avg_log2FC | pct.1 | pct.2 | p_val_adj |
|---|---|---|---|---|---|
| Rps27 | 3.46E-204 | 2.16289029 | 0.942 | 0.901 | 9.35E-200 |
| Rps28 | 3.34E-202 | 2.38625357 | 0.93 | 0.873 | 9.02E-198 |
| Pax7 | 3.23E-178 | -1.7733129 | 0.111 | 0.793 | 8.72E-174 |
| Jund | 1.88E-177 | -2.064348 | 0.37 | 0.914 | 5.08E-173 |
| Junb | 8.78E-177 | -2.0112063 | 0.431 | 0.919 | 2.37E-172 |
| Igfbp5 | 8.01E-175 | -2.3880187 | 0.081 | 0.712 | 2.17E-170 |
| Dag1 | 8.14E-172 | -1.8254447 | 0.136 | 0.802 | 2.20E-167 |
| Pdlim4 | 1.77E-162 | -1.6467643 | 0.079 | 0.691 | 4.78E-158 |
| Sparc | 9.03E-162 | -2.0279536 | 0.069 | 0.662 | 2.44E-157 |
| Dnajb1 | 4.07E-159 | -2.946288 | 0.203 | 0.778 | 1.10E-154 |
| Zfp36 | 1.43E-158 | -2.1013255 | 0.15 | 0.773 | 3.86E-154 |
| Crlf1 | 1.98E-158 | -1.4724297 | 0.063 | 0.655 | 5.35E-154 |
| Btg2 | 1.84E-154 | -1.8679911 | 0.283 | 0.903 | 4.99E-150 |
| Rps21 | 8.54E-154 | 1.4876817 | 0.95 | 0.922 | 2.31E-149 |
| Rps29 | 2.49E-153 | 1.93237317 | 0.91 | 0.882 | 6.73E-149 |
| Cebpd | 9.29E-141 | -1.9420619 | 0.305 | 0.854 | 2.51E-136 |
| Nfkbia | 1.45E-140 | -1.9011993 | 0.266 | 0.83 | 3.91E-136 |
| Rpl39 | 3.83E-139 | 1.46132328 | 0.942 | 0.914 | 1.03E-134 |
| Gadd45b | 1.80E-136 | -1.4965545 | 0.185 | 0.793 | 4.87E-132 |
| Gnas | 3.79E-136 | -1.2264763 | 0.644 | 0.975 | 1.03E-131 |
| Meg3 | 8.00E-135 | -1.507766 | 0.284 | 0.872 | 2.16E-130 |
| Notch3 | 3.79E-133 | -1.5775668 | 0.042 | 0.545 | 1.02E-128 |
| Cavin2 | 4.83E-130 | -1.3816983 | 0.066 | 0.581 | 1.31E-125 |
| mt-Atp6 | 1.63E-129 | -0.8249393 | 0.935 | 0.941 | 4.42E-125 |
| Cxcl1 | 4.44E-129 | -2.2420228 | 0.17 | 0.722 | 1.20E-124 |
| Jun | 8.93E-127 | -2.0750232 | 0.387 | 0.883 | 2.42E-122 |
| Lgals1 | 7.84E-122 | -1.0592968 | 0.111 | 0.646 | 2.12E-117 |
| Clmn | 3.08E-120 | -1.3863589 | 0.07 | 0.581 | 8.32E-116 |
| Ier2 | 1.42E-119 | -1.6033121 | 0.157 | 0.699 | 3.83E-115 |
| Fos | 5.33E-119 | -1.9240114 | 0.277 | 0.796 | 1.44E-114 |
| Map1b | 7.89E-118 | -1.3007184 | 0.145 | 0.698 | 2.13E-113 |
| Fosb | 1.73E-117 | -1.6444759 | 0.311 | 0.833 | 4.68E-113 |
| Eif3f | 3.92E-115 | -0.9776187 | 0.133 | 0.689 | 1.06E-110 |
| Sytl2 | 9.46E-115 | -1.1660962 | 0.113 | 0.639 | 2.56E-110 |
| Hspa5 | 2.29E-113 | -1.9177372 | 0.501 | 0.901 | 6.19E-109 |
| Crip2 | 2.23E-111 | -1.1700175 | 0.064 | 0.537 | 6.04E-107 |
| Rpl3 | 3.29E-111 | -1.1159722 | 0.533 | 0.918 | 8.88E-107 |
| Tln2 | 1.53E-110 | -1.0985266 | 0.121 | 0.66 | 4.14E-106 |
| Zbtb20 | 4.78E-109 | -1.1136274 | 0.605 | 0.997 | 1.29E-104 |
| Egr1 | 6.85E-109 | -1.2382974 | 0.73 | 0.934 | 1.85E-104 |
| Ppib | 1.98E-108 | -0.8983862 | 0.08 | 0.561 | 5.36E-104 |

| | | | | | |
|---|---|---|---|---|---|
| Col3a1 | 7.65E-104 | -1.4498145 | 0.032 | 0.441 | 2.07E-99 |
| Maff | 1.24E-103 | -1.0271677 | 0.178 | 0.73 | 3.36E-99 |
| Rpl37a | 1.21E-102 | 1.31463378 | 0.919 | 0.902 | 3.28E-98 |
| Eef1d | 2.02E-102 | -0.8162155 | 0.223 | 0.823 | 5.45E-98 |
| Gapdh | 3.70E-100 | -1.0201533 | 0.171 | 0.699 | 1.00E-95 |
| mt-Co2 | 7.24E-100 | -0.7803056 | 0.81 | 0.931 | 1.96E-95 |
| Col4a1 | 2.29E-98 | -0.9665575 | 0.07 | 0.517 | 6.20E-94 |
| mt-Co3 | 3.26E-98 | -0.6879084 | 0.935 | 0.929 | 8.81E-94 |
| Rps5 | 1.92E-97 | -0.8304738 | 0.812 | 0.927 | 5.18E-93 |
| Atp5g2 | 2.06E-97 | -0.7423773 | 0.147 | 0.675 | 5.56E-93 |
| Cux1 | 3.51E-97 | -0.8581107 | 0.109 | 0.6 | 9.50E-93 |
| Socs3 | 5.04E-97 | -1.4573764 | 0.114 | 0.58 | 1.36E-92 |
| Bag3 | 8.46E-97 | -1.3065736 | 0.237 | 0.773 | 2.29E-92 |
| Atp5d | 2.47E-95 | -0.6799355 | 0.107 | 0.591 | 6.69E-91 |
| Itm2a | 1.13E-94 | -1.1835352 | 0.012 | 0.366 | 3.06E-90 |
| Ppp1r15a | 6.81E-94 | -1.1954472 | 0.347 | 0.861 | 1.84E-89 |
| Nfix | 3.29E-93 | -0.8255336 | 0.128 | 0.621 | 8.90E-89 |
| Gem | 9.24E-93 | -0.9259149 | 0.166 | 0.672 | 2.50E-88 |
| Rpl35 | 1.60E-92 | 1.64860751 | 0.795 | 0.761 | 4.32E-88 |
| mt-Cytb | 1.99E-92 | -0.7291742 | 0.798 | 0.963 | 5.37E-88 |
| Pkm | 5.62E-92 | -0.6778874 | 0.089 | 0.54 | 1.52E-87 |
| Pebp1 | 2.07E-91 | -0.8229389 | 0.052 | 0.455 | 5.61E-87 |
| Rpl10-ps3 | 5.73E-91 | -0.7035776 | 0.133 | 0.63 | 1.55E-86 |
| Ppp1r14b | 1.10E-90 | -0.8740405 | 0.09 | 0.53 | 2.98E-86 |
| Aldoa | 1.33E-90 | -0.659843 | 0.13 | 0.612 | 3.59E-86 |
| Ttc3 | 5.17E-89 | -0.9096077 | 0.11 | 0.568 | 1.40E-84 |
| Rack1 | 1.85E-88 | -0.9215719 | 0.609 | 0.926 | 5.00E-84 |
| Cd81 | 2.24E-88 | -0.8479926 | 0.071 | 0.485 | 6.05E-84 |
| Capns1 | 3.07E-87 | -0.8493718 | 0.033 | 0.399 | 8.29E-83 |
| Clta | 3.09E-87 | -0.7698356 | 0.088 | 0.512 | 8.36E-83 |
| Chodl | 4.60E-87 | -0.7813398 | 0.23 | 0.791 | 1.24E-82 |
| Eef1g | 1.29E-86 | -0.6100669 | 0.184 | 0.722 | 3.48E-82 |
| Oaz1 | 1.64E-86 | -0.7905449 | 0.255 | 0.843 | 4.43E-82 |
| Ctsz | 5.06E-86 | -0.7900815 | 0.057 | 0.448 | 1.37E-81 |
| Creb5 | 1.19E-85 | -0.9317329 | 0.089 | 0.509 | 3.21E-81 |
| Smc6 | 1.07E-84 | -0.6977733 | 0.126 | 0.596 | 2.90E-80 |
| Timp2 | 2.24E-84 | -0.7651668 | 0.087 | 0.507 | 6.05E-80 |
| Aldh2 | 3.32E-84 | -0.6692259 | 0.085 | 0.509 | 8.99E-80 |
| Gsn | 6.28E-84 | -0.7711089 | 0.155 | 0.639 | 1.70E-79 |
| Cox7a2l | 1.03E-83 | -0.5398062 | 0.153 | 0.653 | 2.78E-79 |
| Apoe | 2.50E-83 | -1.1740735 | 0.369 | 0.858 | 6.76E-79 |
| Slc25a4 | 4.65E-83 | -0.6395719 | 0.165 | 0.659 | 1.26E-78 |
| Rpl38 | 4.73E-83 | 1.50536037 | 0.833 | 0.844 | 1.28E-78 |

| | | | | | |
|---|---|---|---|---|---|
| Fgfr4 | 8.18E-83 | -0.8846488 | 0.047 | 0.421 | 2.21E-78 |
| Nfkbiz | 1.17E-82 | -0.8947936 | 0.13 | 0.586 | 3.17E-78 |
| Birc3 | 2.73E-82 | -0.946543 | 0.053 | 0.424 | 7.39E-78 |
| Vcam1 | 3.23E-82 | -0.3702741 | 0.106 | 0.534 | 8.74E-78 |
| Cdh15 | 3.45E-82 | -1.0889096 | 0.028 | 0.36 | 9.33E-78 |
| Dpt | 5.83E-82 | -0.9405105 | 0.024 | 0.361 | 1.58E-77 |
| Actg1 | 5.96E-82 | -1.1969653 | 0.378 | 0.864 | 1.61E-77 |
| Rpl7 | 6.72E-82 | -0.7716807 | 0.689 | 0.924 | 1.82E-77 |
| Eif3k | 7.98E-82 | -0.5316481 | 0.149 | 0.64 | 2.16E-77 |
| Hmcn2 | 9.45E-82 | -0.9229506 | 0.078 | 0.48 | 2.55E-77 |
| Park7 | 1.60E-81 | -0.8570868 | 0.049 | 0.409 | 4.32E-77 |
| Pde1c | 1.82E-81 | -1.1847334 | 0.046 | 0.409 | 4.91E-77 |
| Atp5c1 | 1.13E-80 | -0.689989 | 0.077 | 0.48 | 3.05E-76 |
| Gnb2 | 1.37E-80 | -0.5619624 | 0.064 | 0.439 | 3.71E-76 |
| Rnf122 | 2.39E-80 | -0.8892711 | 0.039 | 0.392 | 6.46E-76 |
| Spry1 | 2.45E-79 | -1.0203273 | 0.03 | 0.365 | 6.63E-75 |
| Cdk14 | 3.04E-79 | -1.256387 | 0.05 | 0.409 | 8.23E-75 |
| Edn3 | 9.24E-79 | -1.0092517 | 0.029 | 0.359 | 2.50E-74 |
| Col15a1 | 2.72E-78 | -1.0908202 | 0.009 | 0.306 | 7.34E-74 |
| Fxyd6 | 8.26E-78 | -0.7836006 | 0.063 | 0.434 | 2.23E-73 |
| Myf5 | 2.21E-77 | -0.8894413 | 0.057 | 0.419 | 5.97E-73 |
| Smdt1 | 4.86E-77 | -0.7441087 | 0.055 | 0.415 | 1.31E-72 |
| Cfl2 | 8.01E-77 | -0.4546108 | 0.113 | 0.544 | 2.17E-72 |
| Id3 | 9.11E-77 | -1.0988394 | 0.313 | 0.782 | 2.46E-72 |
| Edf1 | 1.38E-76 | -0.6132829 | 0.058 | 0.417 | 3.72E-72 |
| Sema6a | 2.19E-76 | -0.9123432 | 0.09 | 0.487 | 5.91E-72 |
| Nenf | 3.00E-76 | -0.6442633 | 0.053 | 0.408 | 8.11E-72 |
| Hspg2 | 1.99E-75 | -0.7448768 | 0.037 | 0.369 | 5.37E-71 |
| Atp5a1 | 4.15E-75 | -0.4819904 | 0.118 | 0.55 | 1.12E-70 |
| Dot1l | 4.95E-75 | -0.8192218 | 0.06 | 0.421 | 1.34E-70 |
| Ltbp1 | 8.23E-75 | -1.0527407 | 0.072 | 0.446 | 2.22E-70 |
| Ubc | 8.58E-75 | -1.2210652 | 0.628 | 0.92 | 2.32E-70 |
| Eif3h | 2.18E-74 | -0.6092523 | 0.213 | 0.726 | 5.91E-70 |
| Tagln2 | 2.98E-74 | -0.5360297 | 0.122 | 0.555 | 8.05E-70 |
| Ltbp4 | 6.52E-74 | -0.6085412 | 0.092 | 0.489 | 1.76E-69 |
| Gas1 | 6.75E-74 | -0.9617323 | 0.089 | 0.475 | 1.83E-69 |
| Coq10b | 1.76E-73 | -0.5570991 | 0.124 | 0.554 | 4.75E-69 |
| Psmb1 | 2.28E-73 | -0.4236343 | 0.072 | 0.448 | 6.17E-69 |
| Anxa6 | 3.09E-73 | -0.7827304 | 0.052 | 0.398 | 8.35E-69 |
| Cnn3 | 5.46E-73 | -0.4511948 | 0.163 | 0.635 | 1.48E-68 |
| Cd82 | 7.15E-73 | -0.6239435 | 0.078 | 0.453 | 1.93E-68 |
| Snhg12 | 7.82E-73 | -0.4765481 | 0.11 | 0.52 | 2.12E-68 |
| Ank3 | 2.25E-72 | -0.8318175 | 0.08 | 0.452 | 6.08E-68 |

| | | | | | |
|---|---|---|---|---|---|
| Nfib | 4.01E-72 | -0.7055039 | 0.295 | 0.847 | 1.08E-67 |
| Airn | 3.38E-71 | -1.8489422 | 0.032 | 0.339 | 9.13E-67 |
| Gnai2 | 3.49E-71 | -0.2973236 | 0.151 | 0.621 | 9.44E-67 |
| Pde4b | 1.38E-70 | -0.8244975 | 0.385 | 0.894 | 3.72E-66 |
| Dtx4 | 1.43E-70 | -0.7718527 | 0.035 | 0.35 | 3.86E-66 |
| Herpud1 | 4.30E-70 | -0.8155049 | 0.072 | 0.429 | 1.16E-65 |
| Rarres2 | 4.94E-70 | -0.4940419 | 0.124 | 0.541 | 1.34E-65 |
| Pfdn5 | 7.30E-70 | -0.4499985 | 0.247 | 0.787 | 1.98E-65 |
| Tanc2 | 9.16E-70 | -0.7216752 | 0.115 | 0.521 | 2.48E-65 |
| Cdon | 1.39E-69 | -0.833419 | 0.061 | 0.404 | 3.76E-65 |
| Clcn5 | 1.88E-69 | -0.9957631 | 0.047 | 0.368 | 5.07E-65 |
| Vgll4 | 2.32E-69 | -0.8518226 | 0.039 | 0.359 | 6.28E-65 |
| Rap1a | 3.09E-69 | -0.7075439 | 0.031 | 0.338 | 8.36E-65 |
| Ifrd1 | 3.96E-69 | -1.216741 | 0.565 | 0.881 | 1.07E-64 |
| Thra | 6.45E-69 | -0.7034867 | 0.035 | 0.344 | 1.75E-64 |
| Ndufb9 | 1.21E-68 | -0.429586 | 0.097 | 0.488 | 3.27E-64 |
| Ndufb8 | 1.70E-68 | -0.430839 | 0.042 | 0.359 | 4.59E-64 |
| Ndufb11 | 1.89E-68 | -0.60831 | 0.057 | 0.39 | 5.10E-64 |
| Prmt1 | 2.24E-68 | -0.6854704 | 0.024 | 0.318 | 6.05E-64 |
| Rpl37 | 3.04E-68 | 1.09895231 | 0.883 | 0.903 | 8.21E-64 |
| Pmp22 | 4.02E-68 | -0.7677236 | 0.056 | 0.383 | 1.09E-63 |
| Nfic | 4.60E-68 | -0.7626595 | 0.075 | 0.417 | 1.24E-63 |
| Spats2l | 5.41E-68 | -0.7883742 | 0.028 | 0.324 | 1.46E-63 |
| Il11ra1 | 1.54E-67 | -0.6415201 | 0.058 | 0.389 | 4.15E-63 |
| Rpl15 | 2.23E-67 | -0.7863833 | 0.592 | 0.913 | 6.03E-63 |
| Rps2 | 2.28E-67 | -0.7003363 | 0.742 | 0.918 | 6.17E-63 |
| Rps4x | 2.92E-67 | -0.6905759 | 0.852 | 0.936 | 7.89E-63 |
| Hspa1a | 4.10E-67 | -1.2572046 | 0.54 | 0.849 | 1.11E-62 |
| Atp5b | 6.86E-67 | -0.4888468 | 0.205 | 0.698 | 1.86E-62 |
| Atxn10 | 1.05E-66 | -0.7346157 | 0.018 | 0.295 | 2.83E-62 |
| H2-Q4 | 1.56E-66 | -0.8649106 | 0.087 | 0.438 | 4.21E-62 |
| Jsrp1 | 1.58E-66 | -0.6680178 | 0.03 | 0.324 | 4.27E-62 |
| Dusp8 | 2.19E-66 | -0.7510963 | 0.044 | 0.356 | 5.93E-62 |
| Zfhx4 | 2.66E-65 | -0.7043921 | 0.095 | 0.463 | 7.18E-61 |
| Rps3 | 3.21E-65 | -0.6909288 | 0.71 | 0.928 | 8.68E-61 |
| Mafk | 3.97E-65 | -0.5558474 | 0.129 | 0.531 | 1.07E-60 |
| Tspo | 7.05E-65 | -0.6177078 | 0.039 | 0.339 | 1.91E-60 |
| Prdx5 | 8.96E-65 | -0.7991483 | 0.018 | 0.288 | 2.42E-60 |
| Rplp2 | 9.57E-65 | 0.8875285 | 0.925 | 0.917 | 2.59E-60 |
| Sowahc | 1.55E-64 | -1.1288761 | 0.05 | 0.356 | 4.18E-60 |
| Nr3c1 | 1.68E-64 | -0.5487315 | 0.059 | 0.385 | 4.55E-60 |
| Ddx3y | 2.86E-64 | -1.3332829 | 0.021 | 0.288 | 7.72E-60 |
| Atp1a2 | 2.86E-64 | -0.6557613 | 0.052 | 0.362 | 7.74E-60 |

| | | | | | |
|---|---|---|---|---|---|
| U2af1 | 3.53E-64 | -0.2583158 | 0.097 | 0.475 | 9.54E-60 |
| Tgfbr3 | 3.59E-64 | -0.8869567 | 0.163 | 0.589 | 9.71E-60 |
| Stx11 | 4.44E-64 | -0.820456 | 0.022 | 0.298 | 1.20E-59 |
| Lpp | 4.91E-64 | -0.5901278 | 0.097 | 0.461 | 1.33E-59 |
| Ifi27 | 5.08E-64 | -0.6111578 | 0.041 | 0.342 | 1.37E-59 |
| Hs6st2 | 5.44E-64 | -1.1056181 | 0.05 | 0.358 | 1.47E-59 |
| Ptprz1 | 6.01E-64 | -1.1177736 | 0.01 | 0.263 | 1.63E-59 |
| Pxdc1 | 7.28E-64 | -0.6083247 | 0.129 | 0.512 | 1.97E-59 |
| Vim | 9.27E-64 | -0.4842043 | 0.255 | 0.739 | 2.51E-59 |
| Slc3a2 | 9.74E-64 | -0.464784 | 0.089 | 0.438 | 2.63E-59 |
| Csrnp1 | 1.17E-63 | -0.52867 | 0.212 | 0.676 | 3.16E-59 |
| Slc10a6 | 1.65E-63 | -0.7746599 | 0.05 | 0.359 | 4.45E-59 |
| Arf5 | 1.86E-63 | -0.3808488 | 0.126 | 0.52 | 5.03E-59 |
| Pfn1 | 3.69E-63 | -0.4773258 | 0.087 | 0.44 | 9.98E-59 |
| Cryab | 8.08E-63 | -1.521112 | 0.097 | 0.444 | 2.18E-58 |
| Bcl7c | 8.56E-63 | -0.5578521 | 0.054 | 0.364 | 2.31E-58 |
| Arid5a | 1.40E-62 | -0.6388341 | 0.1 | 0.468 | 3.79E-58 |
| Tnxb | 1.58E-62 | -0.8022602 | 0.034 | 0.318 | 4.26E-58 |
| Bcl3 | 1.61E-62 | -0.6492988 | 0.054 | 0.368 | 4.35E-58 |
| Heyl | 4.04E-62 | -0.8020865 | 0.045 | 0.339 | 1.09E-57 |
| Gxylt2 | 4.38E-62 | -0.7290508 | 0.065 | 0.389 | 1.18E-57 |
| Col6a1 | 5.43E-62 | -0.7958618 | 0.021 | 0.283 | 1.47E-57 |
| Rhoa | 6.05E-62 | -0.3955856 | 0.106 | 0.478 | 1.64E-57 |
| H2-K1 | 8.24E-62 | -0.2934874 | 0.122 | 0.509 | 2.23E-57 |
| Vdac2 | 1.01E-61 | -0.5793721 | 0.037 | 0.321 | 2.74E-57 |
| Eif4a2 | 1.24E-61 | -0.424783 | 0.17 | 0.603 | 3.34E-57 |
| Arhgap24 | 1.39E-61 | -1.002039 | 0.102 | 0.455 | 3.76E-57 |
| Eif3m | 1.45E-61 | -0.3193714 | 0.094 | 0.453 | 3.91E-57 |
| Prdx2 | 1.85E-61 | -0.6449461 | 0.041 | 0.324 | 5.01E-57 |
| Use1 | 2.12E-61 | -0.3098324 | 0.171 | 0.598 | 5.74E-57 |
| Rps9 | 2.76E-61 | -0.5863808 | 0.812 | 0.934 | 7.46E-57 |
| Ncoa1 | 3.40E-61 | -0.7827138 | 0.083 | 0.421 | 9.18E-57 |
| Map2 | 4.36E-61 | -0.6901374 | 0.098 | 0.452 | 1.18E-56 |
| Ccnd3 | 5.96E-61 | -0.7722024 | 0.038 | 0.325 | 1.61E-56 |
| Gas6 | 2.50E-60 | -0.741364 | 0.055 | 0.358 | 6.75E-56 |
| Mrpl30 | 3.18E-60 | -0.655397 | 0.04 | 0.322 | 8.59E-56 |
| Lims1 | 5.61E-60 | -0.5948507 | 0.065 | 0.376 | 1.52E-55 |
| Acta1 | 6.40E-60 | -0.7891352 | 0.088 | 0.411 | 1.73E-55 |
| Fry | 8.44E-60 | -0.6611019 | 0.1 | 0.444 | 2.28E-55 |
| Calr | 8.49E-60 | -0.5178715 | 0.146 | 0.544 | 2.30E-55 |
| Eif4b | 8.78E-60 | -0.4582873 | 0.053 | 0.353 | 2.38E-55 |
| Emp3 | 1.26E-59 | -0.3789468 | 0.07 | 0.391 | 3.42E-55 |
| Irf1 | 1.66E-59 | -0.8113012 | 0.04 | 0.324 | 4.48E-55 |

| | | | | | |
|---|---|---|---|---|---|
| Dusp1 | 1.86E-59 | -0.7941434 | 0.173 | 0.569 | 5.03E-55 |
| Man2a2 | 2.10E-59 | -0.6740714 | 0.028 | 0.291 | 5.67E-55 |
| Sde2 | 2.12E-59 | -0.4534261 | 0.103 | 0.456 | 5.73E-55 |
| Zeb1 | 3.30E-59 | -0.432415 | 0.1 | 0.452 | 8.91E-55 |
| Serpinh1 | 4.31E-59 | -0.9066995 | 0.072 | 0.385 | 1.17E-54 |
| Chd7 | 5.36E-59 | -0.5781291 | 0.054 | 0.355 | 1.45E-54 |
| Fkbp2 | 5.62E-59 | -0.645141 | 0.035 | 0.308 | 1.52E-54 |
| Cav1 | 7.59E-59 | -0.6111638 | 0.046 | 0.335 | 2.05E-54 |
| Rplp0 | 8.48E-59 | -0.8382095 | 0.641 | 0.927 | 2.29E-54 |
| Wwtr1 | 9.46E-59 | -0.5399555 | 0.081 | 0.409 | 2.56E-54 |
| Itm2b | 1.02E-58 | -0.3958756 | 0.197 | 0.647 | 2.76E-54 |
| Tubb2b | 1.22E-58 | -0.6574511 | 0.033 | 0.304 | 3.31E-54 |
| Rpl29 | 1.76E-58 | -0.6954447 | 0.51 | 0.909 | 4.77E-54 |
| Birc2 | 2.09E-58 | -0.7059611 | 0.043 | 0.323 | 5.65E-54 |
| Rapsn | 3.61E-58 | -0.7244062 | 0.017 | 0.261 | 9.77E-54 |
| Ube2s | 3.97E-58 | -0.4398486 | 0.094 | 0.436 | 1.07E-53 |
| Top2b | 4.17E-58 | -0.5301858 | 0.058 | 0.357 | 1.13E-53 |
| Selenof | 7.32E-58 | -0.2897526 | 0.106 | 0.461 | 1.98E-53 |
| Eif5a | 1.09E-57 | -0.4171559 | 0.211 | 0.664 | 2.94E-53 |
| Chrdl2 | 1.63E-57 | -0.8760104 | 0.047 | 0.324 | 4.41E-53 |
| Serpinb6a | 2.39E-57 | -0.3731886 | 0.102 | 0.449 | 6.46E-53 |
| Olfml2b | 2.40E-57 | -0.6453722 | 0.027 | 0.284 | 6.48E-53 |
| Esr1 | 5.30E-57 | -0.8973381 | 0.052 | 0.337 | 1.43E-52 |
| Ubtf | 7.21E-57 | -0.6669542 | 0.011 | 0.246 | 1.95E-52 |
| Sntb2 | 1.03E-56 | -0.5452232 | 0.095 | 0.435 | 2.79E-52 |
| Pdzd2 | 1.19E-56 | -0.9192887 | 0.032 | 0.297 | 3.22E-52 |
| Emg1 | 1.59E-56 | -0.4633203 | 0.031 | 0.295 | 4.30E-52 |
| Bmp6 | 2.30E-56 | -1.6435993 | 0.022 | 0.264 | 6.22E-52 |
| Mylpf | 3.80E-56 | -0.5858836 | 0.079 | 0.39 | 1.03E-51 |
| Myh9 | 4.80E-56 | -0.3840926 | 0.071 | 0.384 | 1.30E-51 |
| Tln1 | 4.90E-56 | -0.4759846 | 0.055 | 0.344 | 1.32E-51 |
| Nktr | 5.32E-56 | -0.4703016 | 0.213 | 0.66 | 1.44E-51 |
| Tuba1a | 7.83E-56 | -0.5241854 | 0.08 | 0.399 | 2.12E-51 |
| Iqgap2 | 8.75E-56 | -0.7252978 | 0.039 | 0.307 | 2.37E-51 |
| Ncoa7 | 1.46E-55 | -0.9190174 | 0.03 | 0.285 | 3.96E-51 |
| Foxo1 | 1.76E-55 | -0.9766812 | 0.029 | 0.28 | 4.77E-51 |
| Hmgn1 | 1.96E-55 | -0.4580007 | 0.119 | 0.476 | 5.31E-51 |
| Akr1a1 | 2.13E-55 | -0.4953276 | 0.065 | 0.36 | 5.75E-51 |
| Slf2 | 2.78E-55 | -0.6504548 | 0.044 | 0.316 | 7.52E-51 |
| Bmyc | 3.73E-55 | -0.9221968 | 0.025 | 0.269 | 1.01E-50 |
| Elmsan1 | 4.14E-55 | -0.6296182 | 0.038 | 0.307 | 1.12E-50 |
| Snrpb | 5.29E-55 | -0.3317263 | 0.097 | 0.427 | 1.43E-50 |
| Aes | 6.85E-55 | -0.471949 | 0.046 | 0.318 | 1.85E-50 |

| | | | | | |
|---|---|---|---|---|---|
| Eif4h | 1.45E-54 | -0.3874096 | 0.062 | 0.356 | 3.92E-50 |
| Ddah2 | 1.49E-54 | -0.5805675 | 0.016 | 0.25 | 4.04E-50 |
| Ptov1 | 1.52E-54 | -0.5122411 | 0.045 | 0.318 | 4.10E-50 |
| Mt1 | 3.17E-54 | 1.38081599 | 0.888 | 0.797 | 8.56E-50 |
| Arhgef2 | 4.77E-54 | -0.3716736 | 0.111 | 0.452 | 1.29E-49 |
| Sirt2 | 6.87E-54 | -0.5696897 | 0.041 | 0.306 | 1.86E-49 |
| Ssh2 | 7.99E-54 | -0.7436057 | 0.093 | 0.414 | 2.16E-49 |
| Pde3a | 8.51E-54 | -1.2124062 | 0.019 | 0.25 | 2.30E-49 |
| Rras | 8.63E-54 | -0.481183 | 0.037 | 0.295 | 2.33E-49 |
| Nppc | 1.36E-53 | -1.9934653 | 0.049 | 0.31 | 3.69E-49 |
| Sf3b1 | 1.37E-53 | -0.4678395 | 0.298 | 0.792 | 3.70E-49 |
| Nfkb1 | 1.38E-53 | -0.76745 | 0.195 | 0.583 | 3.72E-49 |
| H2-D1 | 1.48E-53 | -0.272839 | 0.201 | 0.638 | 4.01E-49 |
| Pbrm1 | 1.61E-53 | -0.4006205 | 0.066 | 0.36 | 4.36E-49 |
| Rp9 | 1.68E-53 | -0.3126261 | 0.04 | 0.306 | 4.54E-49 |
| Uqcrh | 1.90E-53 | -0.4160628 | 0.258 | 0.744 | 5.14E-49 |
| Rps12-ps3 | 2.05E-53 | -0.3625368 | 0.07 | 0.362 | 5.54E-49 |
| Rab20 | 2.08E-53 | -0.7872126 | 0.018 | 0.25 | 5.62E-49 |
| Anapc16 | 3.04E-53 | -0.5408982 | 0.038 | 0.295 | 8.23E-49 |
| Gsk3b | 4.07E-53 | -0.403983 | 0.062 | 0.351 | 1.10E-48 |
| Hnrnpr | 6.60E-53 | -0.3813942 | 0.073 | 0.373 | 1.78E-48 |
| Mt2 | 7.12E-53 | 1.4990122 | 0.816 | 0.805 | 1.92E-48 |
| Ndufa13 | 8.16E-53 | -0.3413957 | 0.109 | 0.443 | 2.21E-48 |
| Ccl19 | 8.43E-53 | -1.0513868 | 0.009 | 0.221 | 2.28E-48 |
| Htatsf1 | 9.08E-53 | -0.40608 | 0.05 | 0.32 | 2.46E-48 |
| Akr1b3 | 1.03E-52 | -0.615877 | 0.05 | 0.321 | 2.80E-48 |
| Nfia | 1.12E-52 | -0.662615 | 0.238 | 0.7 | 3.03E-48 |
| Rassf1 | 1.42E-52 | -0.6215698 | 0.039 | 0.296 | 3.85E-48 |
| Dnajc3 | 2.05E-52 | -0.3916035 | 0.095 | 0.42 | 5.55E-48 |
| Dmd | 2.40E-52 | -1.3670502 | 0.192 | 0.593 | 6.49E-48 |
| Idh2 | 2.60E-52 | -0.607196 | 0.018 | 0.245 | 7.02E-48 |
| Rel | 2.76E-52 | -0.4043988 | 0.041 | 0.298 | 7.47E-48 |
| Psap | 3.44E-52 | -0.3508757 | 0.076 | 0.376 | 9.31E-48 |
| Tnnc2 | 4.06E-52 | -0.4534373 | 0.089 | 0.391 | 1.10E-47 |
| Zhx3 | 4.31E-52 | -0.6641593 | 0.061 | 0.341 | 1.16E-47 |
| Antxr2 | 1.03E-51 | -0.7199138 | 0.019 | 0.245 | 2.78E-47 |
| Scand1 | 1.10E-51 | -0.5802733 | 0.04 | 0.292 | 2.98E-47 |
| Tmem109 | 1.12E-51 | -0.628844 | 0.008 | 0.216 | 3.04E-47 |
| Ncald | 1.59E-51 | -0.6386952 | 0.018 | 0.244 | 4.30E-47 |
| Golim4 | 1.81E-51 | -0.3541108 | 0.095 | 0.422 | 4.90E-47 |
| Rpl18 | 2.00E-51 | -0.5159045 | 0.81 | 0.923 | 5.42E-47 |
| Ccnd1 | 2.15E-51 | -0.9339878 | 0.013 | 0.23 | 5.80E-47 |
| Psmb4 | 2.52E-51 | -0.3122106 | 0.067 | 0.348 | 6.81E-47 |

| | | | | | |
|---|---|---|---|---|---|
| Atp5f1 | 2.84E-51 | -0.2597775 | 0.113 | 0.452 | 7.69E-47 |
| Adarb1 | 3.55E-51 | -0.4813955 | 0.07 | 0.354 | 9.60E-47 |
| Tnik | 3.76E-51 | -0.7148741 | 0.092 | 0.398 | 1.02E-46 |
| Creb1 | 4.36E-51 | -0.6859662 | 0.018 | 0.243 | 1.18E-46 |
| Ssr4 | 5.37E-51 | -0.2543228 | 0.085 | 0.388 | 1.45E-46 |
| Rhoh | 6.09E-51 | -0.8709497 | 0.068 | 0.349 | 1.65E-46 |
| Filip1l | 1.50E-50 | -1.0979969 | 0.151 | 0.509 | 4.06E-46 |
| Bicc1 | 1.77E-50 | -0.620149 | 0.079 | 0.377 | 4.79E-46 |
| Irs2 | 1.90E-50 | -0.3613989 | 0.156 | 0.532 | 5.13E-46 |
| Cp | 1.90E-50 | -0.2671133 | 0.117 | 0.452 | 5.14E-46 |
| Pnp | 2.97E-50 | -0.502226 | 0.059 | 0.333 | 8.04E-46 |
| Ank1 | 3.07E-50 | -0.5928817 | 0.019 | 0.242 | 8.29E-46 |
| S1pr3 | 3.49E-50 | -0.5770411 | 0.044 | 0.296 | 9.44E-46 |
| Pde10a | 3.97E-50 | -0.9426846 | 0.067 | 0.344 | 1.07E-45 |
| Ncam1 | 5.84E-50 | -0.3493051 | 0.162 | 0.542 | 1.58E-45 |
| Eif3i | 1.12E-49 | -0.3026001 | 0.076 | 0.368 | 3.03E-45 |
| Atp5g1 | 1.12E-49 | -0.3191846 | 0.091 | 0.395 | 3.03E-45 |
| Cyr61 | 1.26E-49 | -0.9603423 | 0.111 | 0.419 | 3.41E-45 |
| Egfr | 1.28E-49 | -0.7899602 | 0.059 | 0.324 | 3.45E-45 |
| Vtn | 1.43E-49 | -0.6548875 | 0.012 | 0.221 | 3.86E-45 |
| Pik3r1 | 2.41E-49 | -0.5360259 | 0.061 | 0.332 | 6.51E-45 |
| Rtraf | 2.79E-49 | -0.4331757 | 0.057 | 0.328 | 7.56E-45 |
| Mrfap1 | 3.02E-49 | -0.277546 | 0.18 | 0.58 | 8.17E-45 |
| Tecr | 3.38E-49 | -0.4975943 | 0.041 | 0.285 | 9.13E-45 |
| Cct5 | 6.84E-49 | -0.2997865 | 0.074 | 0.36 | 1.85E-44 |
| Prkcq | 7.41E-49 | -0.9215533 | 0.007 | 0.205 | 2.00E-44 |
| Kdm2a | 7.56E-49 | -0.4993567 | 0.038 | 0.282 | 2.04E-44 |
| Uqcrc2 | 8.12E-49 | -0.4328729 | 0.036 | 0.277 | 2.20E-44 |
| Tubb6 | 1.31E-48 | -0.4522848 | 0.084 | 0.379 | 3.55E-44 |
| Eif3d | 1.76E-48 | -0.4528054 | 0.038 | 0.278 | 4.76E-44 |
| Tmem176b | 1.80E-48 | -0.3208412 | 0.148 | 0.5 | 4.85E-44 |
| Ier3 | 3.18E-48 | -0.8546036 | 0.131 | 0.445 | 8.59E-44 |
| Glul | 3.27E-48 | -0.4194395 | 0.157 | 0.515 | 8.83E-44 |
| Anp32a | 4.83E-48 | -0.4429821 | 0.032 | 0.262 | 1.31E-43 |
| Rnf11 | 6.43E-48 | -0.3368654 | 0.066 | 0.337 | 1.74E-43 |
| Psmb6 | 1.01E-47 | -0.2624036 | 0.079 | 0.361 | 2.73E-43 |
| Itpkc | 1.47E-47 | -0.9196627 | 0.028 | 0.248 | 3.96E-43 |
| Ndufv1 | 1.55E-47 | -0.4480724 | 0.046 | 0.289 | 4.18E-43 |
| Impdh2 | 1.63E-47 | -0.4885175 | 0.043 | 0.282 | 4.42E-43 |
| Arhgap5 | 1.98E-47 | -0.3730413 | 0.069 | 0.344 | 5.35E-43 |
| Sri | 2.11E-47 | -0.4946094 | 0.013 | 0.217 | 5.71E-43 |
| Col5a1 | 2.33E-47 | -0.5933992 | 0.011 | 0.211 | 6.29E-43 |
| Rpl8 | 2.53E-47 | -0.4964352 | 0.81 | 0.934 | 6.83E-43 |

| | | | | | |
|---|---|---|---|---|---|
| Drap1 | 2.92E-47 | -0.2703262 | 0.089 | 0.383 | 7.89E-43 |
| Pdia3 | 4.41E-47 | -0.3042309 | 0.096 | 0.399 | 1.19E-42 |
| Atf3 | 5.75E-47 | -0.4882516 | 0.281 | 0.673 | 1.55E-42 |
| Gadd45g | 6.24E-47 | -1.1624329 | 0.066 | 0.321 | 1.69E-42 |
| Pmm1 | 6.51E-47 | -0.6225631 | 0.007 | 0.197 | 1.76E-42 |
| Srsf7 | 6.70E-47 | -0.2530548 | 0.104 | 0.414 | 1.81E-42 |
| Nr4a1 | 8.82E-47 | -0.6085041 | 0.187 | 0.543 | 2.38E-42 |
| Tenm4 | 9.14E-47 | -0.8164347 | 0.066 | 0.329 | 2.47E-42 |
| Rhoc | 1.26E-46 | -0.5578015 | 0.011 | 0.209 | 3.41E-42 |
| Ar | 1.31E-46 | -0.3461366 | 0.108 | 0.415 | 3.53E-42 |
| Crtc3 | 1.37E-46 | -0.3416195 | 0.102 | 0.405 | 3.69E-42 |
| Zfp131 | 1.90E-46 | -0.3647956 | 0.072 | 0.344 | 5.15E-42 |
| Rpl13 | 2.24E-46 | -0.5764568 | 0.908 | 0.924 | 6.05E-42 |
| Atp2b4 | 2.69E-46 | -0.4006711 | 0.051 | 0.3 | 7.27E-42 |
| Snrpd2 | 3.03E-46 | -0.2704704 | 0.078 | 0.353 | 8.20E-42 |
| Sdha | 3.08E-46 | -0.5216576 | 0.02 | 0.229 | 8.34E-42 |
| Actn3 | 3.53E-46 | -0.5133631 | 0.024 | 0.235 | 9.54E-42 |
| Gdi2 | 7.49E-46 | -0.4097697 | 0.051 | 0.3 | 2.03E-41 |
| Psma4 | 7.77E-46 | -0.3172569 | 0.06 | 0.318 | 2.10E-41 |
| 2410015M2C | 8.14E-46 | -0.3845305 | 0.038 | 0.27 | 2.20E-41 |
| Ndufab1 | 8.23E-46 | -0.2877345 | 0.063 | 0.322 | 2.22E-41 |
| Map4 | 9.59E-46 | -0.2539327 | 0.083 | 0.362 | 2.59E-41 |
| Ndufb7 | 1.31E-45 | -0.4436223 | 0.029 | 0.245 | 3.56E-41 |
| Ndufs2 | 2.48E-45 | -0.3823247 | 0.031 | 0.254 | 6.70E-41 |
| Rbms3 | 3.44E-45 | -0.760202 | 0.098 | 0.384 | 9.30E-41 |
| Rpl11 | 4.01E-45 | -0.5364587 | 0.672 | 0.922 | 1.08E-40 |
| Srf | 4.15E-45 | -0.6299519 | 0.009 | 0.196 | 1.12E-40 |
| Litaf | 4.15E-45 | -0.2970535 | 0.085 | 0.367 | 1.12E-40 |
| Hspa13 | 4.59E-45 | -0.521424 | 0.027 | 0.242 | 1.24E-40 |
| Lrig1 | 4.84E-45 | -0.3632844 | 0.065 | 0.326 | 1.31E-40 |
| Mtch1 | 6.87E-45 | -0.5378685 | 0.026 | 0.236 | 1.86E-40 |
| Nedd4 | 8.52E-45 | -0.2603032 | 0.264 | 0.722 | 2.30E-40 |
| Polr2e | 8.76E-45 | -0.6250124 | 0.01 | 0.195 | 2.37E-40 |
| Bcl2 | 9.16E-45 | -0.5946001 | 0.046 | 0.282 | 2.48E-40 |
| Map2k2 | 9.26E-45 | -0.4542088 | 0.029 | 0.244 | 2.50E-40 |
| Tubb5 | 1.09E-44 | -0.4521355 | 0.045 | 0.275 | 2.96E-40 |
| Thsd4 | 1.32E-44 | -0.9388718 | 0.015 | 0.208 | 3.57E-40 |
| Lama2 | 1.35E-44 | -0.7687319 | 0.097 | 0.381 | 3.66E-40 |
| Lsm6 | 1.49E-44 | -0.4342238 | 0.03 | 0.245 | 4.03E-40 |
| Mpdz | 1.74E-44 | -0.3086005 | 0.065 | 0.319 | 4.71E-40 |
| Mrps28 | 1.77E-44 | -0.4485956 | 0.029 | 0.243 | 4.79E-40 |
| Psmb5 | 1.96E-44 | -0.4177294 | 0.03 | 0.244 | 5.30E-40 |
| Ror1 | 2.43E-44 | -1.4487016 | 0.052 | 0.29 | 6.56E-40 |

| | | | | | |
|---|---|---|---|---|---|
| Ccdc50 | 2.44E-44 | -0.3129951 | 0.095 | 0.382 | 6.59E-40 |
| Eif4g3 | 2.48E-44 | -0.5786322 | 0.049 | 0.288 | 6.71E-40 |
| Lhfp | 2.88E-44 | -0.314582 | 0.084 | 0.361 | 7.78E-40 |
| Manf | 2.88E-44 | -0.4034609 | 0.067 | 0.323 | 7.80E-40 |
| Ckm | 3.48E-44 | -0.4917401 | 0.063 | 0.311 | 9.42E-40 |
| Vgll2 | 7.13E-44 | -0.7074675 | 0.028 | 0.237 | 1.93E-39 |
| Calu | 7.24E-44 | -0.3227776 | 0.063 | 0.317 | 1.96E-39 |
| Olfml2a | 8.08E-44 | -0.5405566 | 0.003 | 0.177 | 2.19E-39 |
| Zfhx3 | 8.71E-44 | -0.3209768 | 0.086 | 0.363 | 2.36E-39 |
| Col5a3 | 8.96E-44 | -0.5147927 | 0.034 | 0.253 | 2.42E-39 |
| St3gal5 | 9.70E-44 | -0.5971746 | 0.033 | 0.247 | 2.62E-39 |
| Mpc2 | 1.33E-43 | -0.3542785 | 0.047 | 0.28 | 3.59E-39 |
| Prkab2 | 1.55E-43 | -0.3002698 | 0.055 | 0.299 | 4.18E-39 |
| Nudc | 1.83E-43 | -0.3527908 | 0.05 | 0.282 | 4.94E-39 |
| Rpsa | 1.84E-43 | -0.5107765 | 0.8 | 0.931 | 4.99E-39 |
| Ergic3 | 1.88E-43 | -0.4774716 | 0.02 | 0.218 | 5.08E-39 |
| Cd34 | 2.02E-43 | -0.4656804 | 0.037 | 0.255 | 5.45E-39 |
| Arl3 | 2.11E-43 | -0.3905913 | 0.032 | 0.249 | 5.72E-39 |
| Maged1 | 2.44E-43 | -0.3698369 | 0.052 | 0.289 | 6.61E-39 |
| Col4a2 | 2.83E-43 | -0.4582139 | 0.021 | 0.22 | 7.65E-39 |
| Tmem47 | 4.25E-43 | -0.3457937 | 0.046 | 0.273 | 1.15E-38 |
| Polr1d | 4.73E-43 | -0.2814014 | 0.107 | 0.404 | 1.28E-38 |
| Adgra3 | 5.28E-43 | -0.5961914 | 0.016 | 0.207 | 1.43E-38 |
| Erf | 7.32E-43 | -0.6580004 | 0.016 | 0.205 | 1.98E-38 |
| Zyx | 7.37E-43 | -0.3821897 | 0.065 | 0.316 | 1.99E-38 |
| Nfil3 | 7.50E-43 | -0.5258343 | 0.045 | 0.272 | 2.03E-38 |
| Txn2 | 8.92E-43 | -0.4372746 | 0.016 | 0.207 | 2.41E-38 |
| Igfbp4 | 9.57E-43 | -0.4102626 | 0.071 | 0.328 | 2.59E-38 |
| Bhlhe40 | 1.00E-42 | -0.2936316 | 0.138 | 0.46 | 2.70E-38 |
| Hmgcs1 | 1.02E-42 | -0.6954076 | 0.015 | 0.202 | 2.77E-38 |
| Rpl35a | 1.09E-42 | 0.74649339 | 0.879 | 0.92 | 2.94E-38 |
| Abca8b | 1.19E-42 | -0.361669 | 0.11 | 0.405 | 3.23E-38 |
| Ldlr | 1.22E-42 | -0.7029983 | 0.021 | 0.213 | 3.31E-38 |
| Tnfaip3 | 1.30E-42 | -0.6894266 | 0.062 | 0.305 | 3.52E-38 |
| Galnt17 | 1.62E-42 | -0.945754 | 0.015 | 0.201 | 4.37E-38 |
| Pi4k2b | 3.24E-42 | -0.3841094 | 0.046 | 0.275 | 8.75E-38 |
| Rpl10a | 3.99E-42 | -0.5206195 | 0.639 | 0.922 | 1.08E-37 |
| Cyc1 | 4.73E-42 | -0.3805681 | 0.03 | 0.237 | 1.28E-37 |
| mt-Nd2 | 9.66E-42 | -0.4568374 | 0.64 | 0.953 | 2.61E-37 |
| Phb2 | 1.23E-41 | -0.459562 | 0.029 | 0.231 | 3.32E-37 |
| Adamts1 | 1.39E-41 | -0.3558689 | 0.134 | 0.44 | 3.76E-37 |
| Cuedc1 | 1.87E-41 | -0.4623315 | 0.032 | 0.241 | 5.05E-37 |
| Csnk1d | 2.39E-41 | -0.2524195 | 0.103 | 0.391 | 6.46E-37 |

| | | | | | |
|---|---|---|---|---|---|
| Fth1 | 2.68E-41 | 0.9568081 | 0.901 | 0.932 | 7.25E-37 |
| Lamc1 | 3.35E-41 | -0.2900213 | 0.079 | 0.339 | 9.05E-37 |
| Btg1 | 4.34E-41 | 1.29230918 | 0.75 | 0.859 | 1.17E-36 |
| Bgn | 9.63E-41 | -0.438546 | 0.039 | 0.252 | 2.61E-36 |
| Acaa1a | 1.17E-40 | -0.374613 | 0.034 | 0.239 | 3.18E-36 |
| Megf10 | 1.21E-40 | -0.57388 | 0.05 | 0.273 | 3.28E-36 |
| Atl2 | 1.26E-40 | -0.4651368 | 0.037 | 0.246 | 3.41E-36 |
| Fndc3b | 1.29E-40 | -0.2734997 | 0.09 | 0.359 | 3.49E-36 |
| Smim10l1 | 1.42E-40 | -0.4470786 | 0.021 | 0.209 | 3.85E-36 |
| Tnpo1 | 1.51E-40 | -0.3045936 | 0.082 | 0.338 | 4.10E-36 |
| Fis1 | 1.53E-40 | -0.2812301 | 0.067 | 0.301 | 4.14E-36 |
| Tgfb2 | 1.53E-40 | -0.3811471 | 0.039 | 0.252 | 4.15E-36 |
| Chd9 | 1.64E-40 | -0.5399413 | 0.041 | 0.253 | 4.43E-36 |
| Pdgfa | 1.82E-40 | -0.5671017 | 0.025 | 0.216 | 4.91E-36 |
| Ccnl1 | 2.33E-40 | -0.2939596 | 0.365 | 0.825 | 6.30E-36 |
| Mpst | 2.47E-40 | -0.4927125 | 0.015 | 0.196 | 6.67E-36 |
| Irf2bpl | 2.61E-40 | -0.3997487 | 0.063 | 0.304 | 7.07E-36 |
| Cox4i1 | 2.99E-40 | -0.4399091 | 0.404 | 0.888 | 8.09E-36 |
| Ubb | 3.50E-40 | -0.9481596 | 0.63 | 0.904 | 9.46E-36 |
| Rhob | 3.97E-40 | -0.3962449 | 0.098 | 0.367 | 1.07E-35 |
| Erfe | 4.43E-40 | -0.6282091 | 0.062 | 0.295 | 1.20E-35 |
| Ndufc2 | 4.57E-40 | -0.2564661 | 0.069 | 0.309 | 1.23E-35 |
| Dtnbp1 | 6.27E-40 | -0.5793448 | 0.021 | 0.206 | 1.70E-35 |
| Rps10 | 6.68E-40 | -0.472457 | 0.825 | 0.921 | 1.81E-35 |
| Serpine1 | 9.46E-40 | -0.372117 | 0.111 | 0.39 | 2.56E-35 |
| Pdgfrl | 1.05E-39 | -0.5110673 | 0.011 | 0.183 | 2.83E-35 |
| Psmb2 | 1.17E-39 | -0.432372 | 0.04 | 0.249 | 3.15E-35 |
| Fam168a | 1.54E-39 | -0.6557009 | 0.015 | 0.192 | 4.16E-35 |
| Hip1 | 1.75E-39 | -0.4457325 | 0.04 | 0.249 | 4.74E-35 |
| Psmb7 | 1.96E-39 | -0.2541918 | 0.04 | 0.248 | 5.29E-35 |
| Pcdh7 | 2.12E-39 | -0.6082928 | 0.031 | 0.229 | 5.73E-35 |
| Cnpy2 | 2.31E-39 | -0.3350087 | 0.021 | 0.205 | 6.25E-35 |
| Cyp51 | 2.67E-39 | -0.5500143 | 0.029 | 0.217 | 7.22E-35 |
| Msc | 2.81E-39 | -0.6567846 | 0.022 | 0.207 | 7.61E-35 |
| Ptprg | 3.70E-39 | -0.6347709 | 0.033 | 0.233 | 1.00E-34 |
| Lin7a | 3.72E-39 | -0.7395583 | 0.014 | 0.187 | 1.01E-34 |
| Nudt4 | 4.30E-39 | -0.4691834 | 0.063 | 0.289 | 1.16E-34 |
| Gadd45a | 4.35E-39 | -0.6810907 | 0.251 | 0.601 | 1.18E-34 |
| Pvalb | 4.49E-39 | -0.5300821 | 0.022 | 0.205 | 1.21E-34 |
| Msmo1 | 4.71E-39 | -0.6250595 | 0.028 | 0.214 | 1.27E-34 |
| Ddit3 | 4.76E-39 | -0.5358661 | 0.032 | 0.227 | 1.29E-34 |
| Itpr2 | 5.01E-39 | -0.4326022 | 0.086 | 0.339 | 1.35E-34 |
| Cxxc5 | 7.34E-39 | -0.4815473 | 0.014 | 0.187 | 1.99E-34 |

| | | | | | |
|---|---|---|---|---|---|
| Ptpra | 7.88E-39 | -0.3974235 | 0.029 | 0.22 | 2.13E-34 |
| Dcaf8 | 8.64E-39 | -0.2600133 | 0.075 | 0.319 | 2.34E-34 |
| Idi1 | 1.08E-38 | -0.6222136 | 0.022 | 0.201 | 2.92E-34 |
| Ufc1 | 1.10E-38 | -0.3594988 | 0.023 | 0.207 | 2.98E-34 |
| Selenom | 1.27E-38 | -0.2617578 | 0.067 | 0.298 | 3.42E-34 |
| Ccl2 | 1.37E-38 | -2.1274454 | 0.043 | 0.242 | 3.70E-34 |
| Rpl7a | 1.60E-38 | -0.4922872 | 0.538 | 0.897 | 4.33E-34 |
| Tmem176a | 1.74E-38 | -0.2505546 | 0.085 | 0.34 | 4.71E-34 |
| Ddx5 | 1.79E-38 | 0.88670002 | 0.822 | 0.931 | 4.85E-34 |
| Mvp | 1.81E-38 | -0.4200375 | 0.029 | 0.22 | 4.90E-34 |
| Cavin3 | 1.82E-38 | -0.3703729 | 0.038 | 0.237 | 4.92E-34 |
| Uqcr11 | 1.83E-38 | -0.2636723 | 0.069 | 0.303 | 4.96E-34 |
| Sertad1 | 1.95E-38 | -0.2651819 | 0.066 | 0.299 | 5.27E-34 |
| Sec11a | 2.06E-38 | -0.3380476 | 0.037 | 0.239 | 5.58E-34 |
| Trib1 | 2.39E-38 | -0.645502 | 0.359 | 0.745 | 6.45E-34 |
| Rtn2 | 2.95E-38 | -0.5356928 | 0.015 | 0.185 | 7.98E-34 |
| Traf3ip3 | 3.74E-38 | -0.5069275 | 0.02 | 0.198 | 1.01E-33 |
| Ankrd29 | 4.09E-38 | -0.5644836 | 0.01 | 0.174 | 1.11E-33 |
| Dbp | 4.35E-38 | -0.5793455 | 0.017 | 0.19 | 1.18E-33 |
| Abi3bp | 4.40E-38 | -0.459956 | 0.028 | 0.217 | 1.19E-33 |
| Rpl24 | 5.14E-38 | 0.76647379 | 0.878 | 0.914 | 1.39E-33 |
| Ndufv2 | 7.36E-38 | -0.3375152 | 0.034 | 0.228 | 1.99E-33 |
| Rps7 | 8.64E-38 | -0.4409219 | 0.806 | 0.919 | 2.34E-33 |
| Calcr | 9.85E-38 | -0.6958937 | 0.017 | 0.19 | 2.66E-33 |
| Chpt1 | 1.24E-37 | -0.476516 | 0.035 | 0.229 | 3.35E-33 |
| Csnk1e | 1.32E-37 | -0.268556 | 0.062 | 0.289 | 3.56E-33 |
| Zfp568 | 1.38E-37 | -0.7837802 | 0.023 | 0.197 | 3.74E-33 |
| Nr4a3 | 1.47E-37 | -0.6222589 | 0.034 | 0.227 | 3.96E-33 |
| Ing2 | 1.55E-37 | -0.5100515 | 0.034 | 0.225 | 4.18E-33 |
| Rpl6 | 1.57E-37 | 0.76529938 | 0.906 | 0.922 | 4.26E-33 |
| Eif6 | 1.66E-37 | -0.4625072 | 0.03 | 0.217 | 4.49E-33 |
| Trip10 | 1.97E-37 | -0.4078311 | 0.025 | 0.208 | 5.33E-33 |
| Pde4d | 2.85E-37 | -1.0670906 | 0.103 | 0.36 | 7.71E-33 |
| Igsf3 | 2.89E-37 | -0.4686792 | 0.024 | 0.204 | 7.81E-33 |
| Ehmt1 | 3.80E-37 | -0.5327054 | 0.012 | 0.176 | 1.03E-32 |
| Aldh1a1 | 3.93E-37 | -0.5162956 | 0.011 | 0.17 | 1.06E-32 |
| Ikzf4 | 3.93E-37 | -0.6622505 | 0.022 | 0.198 | 1.06E-32 |
| 5-Mar | 5.10E-37 | -0.3414006 | 0.04 | 0.24 | 1.38E-32 |
| Smg6 | 5.35E-37 | -0.3015289 | 0.054 | 0.269 | 1.45E-32 |
| Egr2 | 5.93E-37 | -0.5122328 | 0.049 | 0.254 | 1.60E-32 |
| Gbf1 | 5.95E-37 | -0.410143 | 0.021 | 0.197 | 1.61E-32 |
| Taok1 | 6.20E-37 | -0.3484273 | 0.072 | 0.309 | 1.68E-32 |
| Cdk4 | 6.75E-37 | -0.3673219 | 0.027 | 0.208 | 1.82E-32 |

| | | | | | |
|---|---|---|---|---|---|
| Zfp148 | 1.12E-36 | -0.3896681 | 0.035 | 0.227 | 3.03E-32 |
| Pcbp4 | 1.12E-36 | -0.4052567 | 0.014 | 0.179 | 3.04E-32 |
| Fkbp4 | 1.69E-36 | -0.2769235 | 0.071 | 0.3 | 4.57E-32 |
| Fstl1 | 2.22E-36 | -0.3221179 | 0.054 | 0.268 | 6.00E-32 |
| Eif3g | 2.92E-36 | -0.2625042 | 0.036 | 0.228 | 7.89E-32 |
| Unc93b1 | 3.16E-36 | -0.6032044 | 0.004 | 0.15 | 8.55E-32 |
| Ppia | 3.22E-36 | -0.3518629 | 0.445 | 0.883 | 8.72E-32 |
| Ahdc1 | 4.92E-36 | -0.6053824 | 0.02 | 0.189 | 1.33E-31 |
| Atraid | 5.35E-36 | -0.339655 | 0.027 | 0.205 | 1.45E-31 |
| Relb | 5.41E-36 | -0.2589801 | 0.057 | 0.273 | 1.46E-31 |
| Rpl19 | 6.29E-36 | -0.4269437 | 0.844 | 0.921 | 1.70E-31 |
| Dynlt3 | 6.64E-36 | -0.2833693 | 0.05 | 0.253 | 1.80E-31 |
| Ttc28 | 7.41E-36 | -0.5002924 | 0.08 | 0.316 | 2.00E-31 |
| Gm2564 | 9.48E-36 | -0.6791405 | 0.007 | 0.155 | 2.56E-31 |
| Psmg4 | 9.97E-36 | -0.4053892 | 0.018 | 0.185 | 2.70E-31 |
| Hsbp1 | 1.04E-35 | -0.2691393 | 0.043 | 0.237 | 2.82E-31 |
| Per1 | 1.11E-35 | -0.4735747 | 0.011 | 0.168 | 2.99E-31 |
| Gab1 | 1.46E-35 | -0.6068219 | 0.023 | 0.195 | 3.96E-31 |
| Trim44 | 1.48E-35 | -0.4056276 | 0.024 | 0.197 | 4.01E-31 |
| Eef2 | 1.70E-35 | -0.4380848 | 0.495 | 0.909 | 4.61E-31 |
| Zfp622 | 1.73E-35 | -0.4024159 | 0.031 | 0.214 | 4.68E-31 |
| Erp29 | 1.85E-35 | -0.4007664 | 0.016 | 0.177 | 5.01E-31 |
| C1qtnf3 | 2.22E-35 | -0.6715884 | 0.004 | 0.147 | 6.00E-31 |
| Ube2r2 | 2.63E-35 | -0.3209016 | 0.037 | 0.226 | 7.11E-31 |
| Wipf1 | 2.76E-35 | -0.5160268 | 0.022 | 0.193 | 7.45E-31 |
| Zmym4 | 2.76E-35 | -0.5659625 | 0.02 | 0.187 | 7.47E-31 |
| Rpl9 | 3.24E-35 | -0.3920004 | 0.884 | 0.927 | 8.77E-31 |
| Msi2 | 4.28E-35 | -0.4123379 | 0.083 | 0.318 | 1.16E-30 |
| Pdha1 | 4.49E-35 | -0.378678 | 0.026 | 0.199 | 1.21E-30 |
| Dhcr24 | 5.02E-35 | -0.300343 | 0.06 | 0.275 | 1.36E-30 |
| Nfkbie | 5.14E-35 | -0.4352146 | 0.016 | 0.178 | 1.39E-30 |
| Vwa1 | 5.50E-35 | -0.4900908 | 0.016 | 0.177 | 1.49E-30 |
| Grb10 | 6.98E-35 | -0.3909957 | 0.054 | 0.26 | 1.89E-30 |
| Dguok | 7.67E-35 | -0.4671044 | 0.003 | 0.143 | 2.07E-30 |
| Zfp36l1 | 8.03E-35 | -0.3913052 | 0.442 | 0.843 | 2.17E-30 |
| Nf1 | 9.28E-35 | -0.286539 | 0.041 | 0.234 | 2.51E-30 |
| N6amt1 | 9.36E-35 | -0.3156592 | 0.046 | 0.241 | 2.53E-30 |
| Garem1 | 1.12E-34 | -0.7344009 | 0.026 | 0.196 | 3.02E-30 |
| Fkbp8 | 1.17E-34 | -0.457323 | 0.014 | 0.17 | 3.18E-30 |
| Fhl1 | 1.22E-34 | -0.3334018 | 0.014 | 0.17 | 3.30E-30 |
| Thap3 | 1.22E-34 | -0.4871812 | 0.016 | 0.176 | 3.30E-30 |
| Snrpa | 1.29E-34 | -0.3880045 | 0.024 | 0.192 | 3.49E-30 |
| Ankrd10 | 1.50E-34 | -0.4056141 | 0.028 | 0.204 | 4.07E-30 |

| | | | | | |
|---|---|---|---|---|---|
| Malsu1 | 1.53E-34 | -0.2880394 | 0.037 | 0.222 | 4.14E-30 |
| Poldip3 | 1.60E-34 | -0.3093988 | 0.008 | 0.155 | 4.33E-30 |
| 2-Mar | 1.66E-34 | -0.3907493 | 0.016 | 0.173 | 4.48E-30 |
| Capzb | 2.36E-34 | -0.284509 | 0.032 | 0.212 | 6.39E-30 |
| Ank2 | 2.43E-34 | -0.5554086 | 0.068 | 0.286 | 6.57E-30 |
| Polr2m | 2.53E-34 | -0.2679005 | 0.023 | 0.192 | 6.83E-30 |
| Scn1b | 3.73E-34 | -0.4080362 | 0.004 | 0.143 | 1.01E-29 |
| Rnf130 | 4.16E-34 | -0.5520644 | 0.003 | 0.14 | 1.12E-29 |
| Iqsec3 | 4.19E-34 | -0.4998354 | 0 | 0.132 | 1.13E-29 |
| Acbd3 | 4.99E-34 | -0.3025902 | 0.035 | 0.217 | 1.35E-29 |
| Taldo1 | 5.22E-34 | -0.2528251 | 0.022 | 0.187 | 1.41E-29 |
| Atp6v1a | 5.60E-34 | -0.3073771 | 0.023 | 0.19 | 1.51E-29 |
| Bin1 | 6.26E-34 | -0.3295214 | 0.028 | 0.2 | 1.69E-29 |
| Exoc4 | 6.58E-34 | -0.7348664 | 0.019 | 0.179 | 1.78E-29 |
| Rasa1 | 6.94E-34 | -0.4676867 | 0.032 | 0.209 | 1.88E-29 |
| Spock3 | 7.74E-34 | -0.8576019 | 0.009 | 0.153 | 2.09E-29 |
| Tmtc2 | 8.84E-34 | -0.8751851 | 0.016 | 0.172 | 2.39E-29 |
| Dync1li2 | 8.86E-34 | -0.2919223 | 0.025 | 0.192 | 2.40E-29 |
| Rfx7 | 1.02E-33 | -0.3111932 | 0.028 | 0.198 | 2.76E-29 |
| Dnaja4 | 1.11E-33 | -0.9256496 | 0.05 | 0.241 | 3.00E-29 |
| Cdkn2d | 1.13E-33 | -0.4876667 | 0.022 | 0.186 | 3.06E-29 |
| mt-Nd4 | 1.19E-33 | -0.3829203 | 0.706 | 0.952 | 3.21E-29 |
| Adamts5 | 1.21E-33 | -0.3454005 | 0.083 | 0.306 | 3.27E-29 |
| Brwd1 | 1.22E-33 | -0.3219099 | 0.05 | 0.247 | 3.31E-29 |
| Bax | 1.23E-33 | -0.4064661 | 0.014 | 0.167 | 3.33E-29 |
| Rnh1 | 1.29E-33 | -0.4398386 | 0.038 | 0.218 | 3.49E-29 |
| Sra1 | 1.35E-33 | -0.4130449 | 0.01 | 0.157 | 3.66E-29 |
| Palld | 1.80E-33 | -0.436267 | 0.082 | 0.307 | 4.88E-29 |
| Rev3l | 1.89E-33 | -0.283864 | 0.06 | 0.267 | 5.11E-29 |
| Atp5o.1 | 1.98E-33 | -0.349028 | 0.036 | 0.214 | 5.36E-29 |
| Chd8 | 2.36E-33 | -0.3830603 | 0.023 | 0.186 | 6.39E-29 |
| Plcb4 | 2.69E-33 | -0.4460317 | 0.023 | 0.187 | 7.28E-29 |
| Ube2e2 | 3.13E-33 | -0.6641333 | 0.011 | 0.158 | 8.46E-29 |
| Psmd8 | 3.29E-33 | -0.283464 | 0.044 | 0.228 | 8.90E-29 |
| Zbtb7a | 3.35E-33 | -0.2870726 | 0.046 | 0.233 | 9.07E-29 |
| Socs2 | 3.62E-33 | -0.394866 | 0.039 | 0.221 | 9.79E-29 |
| Mcrip1 | 3.80E-33 | -0.2682775 | 0.021 | 0.182 | 1.03E-28 |
| Rnaset2a | 4.15E-33 | -0.3629825 | 0.012 | 0.161 | 1.12E-28 |
| Gm8797 | 4.21E-33 | -0.4258598 | 0.019 | 0.177 | 1.14E-28 |
| Rpl18a | 4.64E-33 | -0.402713 | 0.805 | 0.919 | 1.25E-28 |
| Ttc19 | 4.84E-33 | -0.373499 | 0.028 | 0.195 | 1.31E-28 |
| Jag1 | 5.31E-33 | -0.4723075 | 0.016 | 0.169 | 1.44E-28 |
| Cyth2 | 5.62E-33 | -0.3421785 | 0.023 | 0.184 | 1.52E-28 |

| | | | | | |
|---|---|---|---|---|---|
| Lars2 | 6.32E-33 | -0.4445298 | 0.129 | 0.388 | 1.71E-28 |
| Colgalt1 | 7.46E-33 | -0.305468 | 0.015 | 0.167 | 2.02E-28 |
| Hint2 | 8.05E-33 | -0.3784944 | 0.013 | 0.16 | 2.18E-28 |
| Ndufs8 | 9.35E-33 | -0.2796218 | 0.022 | 0.182 | 2.53E-28 |
| Zbtb16 | 9.82E-33 | -0.2609595 | 0.061 | 0.267 | 2.65E-28 |
| Tppp3 | 1.04E-32 | -0.489081 | 0.017 | 0.17 | 2.82E-28 |
| 9530068E07I | 1.14E-32 | -0.2818348 | 0.02 | 0.177 | 3.08E-28 |
| Trmt112 | 1.26E-32 | -0.2733217 | 0.03 | 0.201 | 3.41E-28 |
| Bnc2 | 1.27E-32 | -0.8276903 | 0.045 | 0.229 | 3.44E-28 |
| Ptprd | 1.39E-32 | -0.3968967 | 0.097 | 0.337 | 3.75E-28 |
| Usp2 | 1.40E-32 | -0.3382259 | 0.05 | 0.243 | 3.78E-28 |
| Foxp2 | 1.40E-32 | -0.6632467 | 0.071 | 0.28 | 3.80E-28 |
| Pard3b | 1.41E-32 | -0.7475002 | 0.026 | 0.189 | 3.82E-28 |
| Kdelr1 | 2.44E-32 | -0.2857757 | 0.03 | 0.197 | 6.59E-28 |
| Ltbp3 | 2.54E-32 | -0.256291 | 0.02 | 0.177 | 6.88E-28 |
| Amotl2 | 2.64E-32 | -0.4725973 | 0.024 | 0.185 | 7.15E-28 |
| Rps8 | 2.90E-32 | 0.4815376 | 0.974 | 0.927 | 7.85E-28 |
| 2210016L21I | 2.95E-32 | -0.3278902 | 0.012 | 0.158 | 7.98E-28 |
| Txndc17 | 3.15E-32 | -0.2624164 | 0.043 | 0.225 | 8.53E-28 |
| Rex1bd | 3.86E-32 | -0.3488035 | 0.024 | 0.183 | 1.04E-27 |
| Lmo2 | 4.16E-32 | -0.4173161 | 0.01 | 0.153 | 1.12E-27 |
| Stx4a | 5.11E-32 | -0.2573467 | 0.021 | 0.178 | 1.38E-27 |
| Eef1b2 | 5.70E-32 | -0.3739898 | 0.47 | 0.882 | 1.54E-27 |
| Jarid2 | 5.80E-32 | -0.3248919 | 0.052 | 0.245 | 1.57E-27 |
| Larp4b | 6.05E-32 | -0.4313562 | 0.023 | 0.182 | 1.64E-27 |
| Prdx4 | 7.08E-32 | -0.332758 | 0.021 | 0.177 | 1.92E-27 |
| Bod1l | 7.12E-32 | -0.2503463 | 0.082 | 0.303 | 1.93E-27 |
| Loxl2 | 1.09E-31 | -0.5587632 | 0.01 | 0.15 | 2.95E-27 |
| Ola1 | 1.11E-31 | -0.3070987 | 0.035 | 0.204 | 3.00E-27 |
| Snx1 | 1.29E-31 | -0.4071428 | 0.02 | 0.174 | 3.48E-27 |
| Hk1 | 1.67E-31 | -0.5515617 | 0.007 | 0.139 | 4.50E-27 |
| Fcgrt | 1.71E-31 | -0.4471117 | 0.016 | 0.16 | 4.61E-27 |
| 2810474O19 | 1.95E-31 | -0.3576039 | 0.066 | 0.268 | 5.28E-27 |
| Cacybp | 2.30E-31 | -0.3179533 | 0.154 | 0.429 | 6.22E-27 |
| Hmgcr | 2.37E-31 | -0.4295212 | 0.017 | 0.166 | 6.42E-27 |
| Fam129a | 2.88E-31 | -0.3785956 | 0.034 | 0.203 | 7.78E-27 |
| Mapk8 | 3.13E-31 | -0.3635037 | 0.029 | 0.19 | 8.47E-27 |
| Deptor | 3.43E-31 | -0.3202526 | 0.103 | 0.344 | 9.27E-27 |
| Dtd1 | 3.60E-31 | -0.348756 | 0.017 | 0.164 | 9.75E-27 |
| Mxra8 | 3.81E-31 | -0.2565547 | 0.023 | 0.178 | 1.03E-26 |
| Igdcc4 | 5.26E-31 | -0.4468176 | 0.008 | 0.14 | 1.42E-26 |
| Mtmr3 | 7.27E-31 | -0.5010492 | 0.012 | 0.152 | 1.97E-26 |
| Ostc | 7.59E-31 | -0.2945421 | 0.012 | 0.153 | 2.05E-26 |

| | | | | | |
|---|---|---|---|---|---|
| Fbn1 | 8.34E-31 | -0.2648682 | 0.034 | 0.2 | 2.25E-26 |
| Ppp1r14a | 8.34E-31 | -0.424486 | 0.009 | 0.143 | 2.26E-26 |
| Pja2 | 8.49E-31 | -0.3518539 | 0.031 | 0.196 | 2.30E-26 |
| Mllt3 | 8.54E-31 | -0.5448201 | 0.017 | 0.161 | 2.31E-26 |
| Echs1 | 9.56E-31 | -0.3403496 | 0.015 | 0.158 | 2.59E-26 |
| Mrpl42 | 9.65E-31 | -0.363466 | 0.02 | 0.169 | 2.61E-26 |
| Zbtb38 | 1.10E-30 | -0.5094634 | 0.02 | 0.167 | 2.99E-26 |
| Npdc1 | 1.24E-30 | -0.3578861 | 0.009 | 0.142 | 3.34E-26 |
| Fam96b | 1.31E-30 | -0.3512111 | 0.025 | 0.177 | 3.55E-26 |
| Abhd14a | 1.46E-30 | -0.375515 | 0.01 | 0.146 | 3.94E-26 |
| Pgm5 | 1.73E-30 | -0.5111344 | 0.025 | 0.179 | 4.67E-26 |
| Ube2e1 | 1.83E-30 | -0.430786 | 0.025 | 0.177 | 4.94E-26 |
| Cdc42ep5 | 2.40E-30 | -0.4258593 | 0.011 | 0.146 | 6.49E-26 |
| Ubr3 | 2.79E-30 | -0.4501648 | 0.025 | 0.178 | 7.55E-26 |
| Szrd1 | 3.17E-30 | -0.379536 | 0.015 | 0.157 | 8.57E-26 |
| Ehd2 | 3.41E-30 | -0.3736116 | 0.01 | 0.145 | 9.23E-26 |
| Ntn5 | 3.75E-30 | -0.4424118 | 0.022 | 0.168 | 1.01E-25 |
| Id1 | 3.97E-30 | -0.5054184 | 0.079 | 0.283 | 1.07E-25 |
| Rpl10 | 4.25E-30 | -0.3489356 | 0.776 | 0.918 | 1.15E-25 |
| Atf5 | 4.34E-30 | -0.3982304 | 0.009 | 0.14 | 1.17E-25 |
| Tpi1 | 4.99E-30 | -0.2641648 | 0.027 | 0.182 | 1.35E-25 |
| Trio | 5.77E-30 | -0.3946556 | 0.025 | 0.177 | 1.56E-25 |
| Hoxc10 | 5.80E-30 | -0.2837014 | 0.036 | 0.202 | 1.57E-25 |
| Islr | 6.93E-30 | -0.4819928 | 0.006 | 0.13 | 1.87E-25 |
| Eci2 | 7.08E-30 | -0.3167765 | 0.019 | 0.164 | 1.92E-25 |
| Zc3h14 | 1.00E-29 | -0.3104337 | 0.014 | 0.152 | 2.71E-25 |
| Itpripl2 | 1.04E-29 | -0.2683904 | 0.029 | 0.186 | 2.81E-25 |
| Tnip1 | 1.10E-29 | -0.469352 | 0.01 | 0.14 | 2.98E-25 |
| Idh3g | 1.43E-29 | -0.2850872 | 0.027 | 0.178 | 3.86E-25 |
| Gm17056 | 1.48E-29 | -0.4111889 | 0.037 | 0.199 | 4.01E-25 |
| Sptssa | 1.56E-29 | -0.2764762 | 0.021 | 0.167 | 4.22E-25 |
| Ephx1 | 1.97E-29 | -0.4635748 | 0.015 | 0.152 | 5.32E-25 |
| Herpud2 | 2.44E-29 | -0.3378677 | 0.012 | 0.146 | 6.59E-25 |
| Pcmtd1 | 2.71E-29 | -0.3092967 | 0.043 | 0.213 | 7.34E-25 |
| Klhl13 | 3.28E-29 | -0.3644862 | 0.01 | 0.14 | 8.86E-25 |
| Ppp2r2a | 3.44E-29 | 0.31792111 | 0.126 | 0.379 | 9.32E-25 |
| St5 | 3.80E-29 | -0.3330185 | 0.034 | 0.194 | 1.03E-24 |
| Phf14 | 4.38E-29 | -0.4794402 | 0.028 | 0.178 | 1.19E-24 |
| Fscn1 | 4.42E-29 | -0.3461264 | 0.018 | 0.157 | 1.20E-24 |
| Kcnma1 | 4.69E-29 | -0.7752846 | 0.013 | 0.146 | 1.27E-24 |
| Polr2j | 5.02E-29 | -0.3026111 | 0.011 | 0.143 | 1.36E-24 |
| Kat6b | 5.50E-29 | -0.3253332 | 0.05 | 0.227 | 1.49E-24 |
| Hcfc1r1 | 5.72E-29 | -0.2642146 | 0.029 | 0.18 | 1.55E-24 |

| | | | | | |
|---|---|---|---|---|---|
| Pias1 | 5.79E-29 | -0.3541529 | 0.026 | 0.176 | 1.57E-24 |
| Pfdn1 | 6.46E-29 | -0.2570448 | 0.03 | 0.182 | 1.75E-24 |
| Mettl26 | 6.77E-29 | -0.4191222 | 0.012 | 0.143 | 1.83E-24 |
| Rad50 | 6.89E-29 | -0.3733958 | 0.015 | 0.15 | 1.86E-24 |
| Dock7 | 8.09E-29 | -0.4299034 | 0.015 | 0.15 | 2.19E-24 |
| Dock4 | 8.42E-29 | -0.6860022 | 0.009 | 0.135 | 2.28E-24 |
| Emc10 | 9.67E-29 | -0.4357438 | 0.017 | 0.155 | 2.61E-24 |
| Csrp1 | 9.84E-29 | -0.3448162 | 0.03 | 0.181 | 2.66E-24 |
| Rsrc1 | 9.90E-29 | -0.3145409 | 0.042 | 0.208 | 2.68E-24 |
| Irgq | 1.14E-28 | -0.4650051 | 0.009 | 0.133 | 3.07E-24 |
| Trps1 | 1.20E-28 | -0.4548713 | 0.076 | 0.277 | 3.24E-24 |
| Mmp2 | 1.20E-28 | -0.2697622 | 0.026 | 0.175 | 3.24E-24 |
| Col6a2 | 1.21E-28 | -0.3883386 | 0.003 | 0.118 | 3.28E-24 |
| Col19a1 | 1.28E-28 | -0.6017945 | 0.017 | 0.155 | 3.45E-24 |
| Rem1 | 1.34E-28 | -0.4168626 | 0.002 | 0.116 | 3.63E-24 |
| Rps16 | 1.38E-28 | -0.3619701 | 0.789 | 0.922 | 3.74E-24 |
| G3bp2 | 1.46E-28 | -0.302314 | 0.037 | 0.197 | 3.95E-24 |
| Malat1 | 1.52E-28 | 0.32209325 | 0.977 | 1 | 4.12E-24 |
| Lrch3 | 1.61E-28 | -0.3543576 | 0.022 | 0.164 | 4.34E-24 |
| Ptprf | 1.62E-28 | -0.4291137 | 0.005 | 0.124 | 4.38E-24 |
| Usp3 | 1.96E-28 | -0.3605266 | 0.01 | 0.138 | 5.30E-24 |
| Bok | 1.99E-28 | -0.424739 | 0.004 | 0.12 | 5.38E-24 |
| Pea15a | 2.02E-28 | -0.3850967 | 0.021 | 0.16 | 5.46E-24 |
| Ppic | 2.04E-28 | -0.3130291 | 0.01 | 0.135 | 5.51E-24 |
| Adamts4 | 2.11E-28 | -0.5798237 | 0.036 | 0.195 | 5.71E-24 |
| Nrep | 3.96E-28 | -0.3539957 | 0.011 | 0.137 | 1.07E-23 |
| Gnaq | 3.99E-28 | -0.3272234 | 0.046 | 0.211 | 1.08E-23 |
| Commd1 | 4.23E-28 | -0.281566 | 0.022 | 0.163 | 1.14E-23 |
| Rad23a | 4.35E-28 | -0.355605 | 0.026 | 0.172 | 1.18E-23 |
| Tmem131 | 4.78E-28 | -0.3642612 | 0.027 | 0.175 | 1.29E-23 |
| Antxr1 | 5.11E-28 | -0.3495938 | 0.018 | 0.155 | 1.38E-23 |
| Col5a2 | 5.78E-28 | -0.295614 | 0.057 | 0.237 | 1.56E-23 |
| Carhsp1 | 5.83E-28 | -0.2738615 | 0.024 | 0.166 | 1.58E-23 |
| Chchd3 | 6.03E-28 | -0.323786 | 0.025 | 0.168 | 1.63E-23 |
| Suclg1 | 6.16E-28 | -0.4146528 | 0.011 | 0.137 | 1.67E-23 |
| Smim3 | 7.18E-28 | -0.4588422 | 0.016 | 0.149 | 1.94E-23 |
| Lamtor2 | 7.21E-28 | -0.2549303 | 0.02 | 0.157 | 1.95E-23 |
| Lamtor4 | 7.28E-28 | -0.322793 | 0.013 | 0.143 | 1.97E-23 |
| Mettl9 | 7.67E-28 | -0.2951978 | 0.01 | 0.134 | 2.07E-23 |
| Rpl36 | 1.37E-27 | 0.79035425 | 0.824 | 0.908 | 3.69E-23 |
| Rpn2 | 1.37E-27 | -0.3035458 | 0.019 | 0.155 | 3.72E-23 |
| Tiprl | 1.56E-27 | -0.3251424 | 0.011 | 0.136 | 4.22E-23 |
| Eya4 | 1.57E-27 | -0.5272062 | 0.043 | 0.205 | 4.26E-23 |

| | | | | | |
|---|---|---|---|---|---|
| Pura | 1.95E-27 | 0.26233299 | 0.145 | 0.414 | 5.28E-23 |
| Rpl41 | 2.02E-27 | 0.78107573 | 0.849 | 0.901 | 5.47E-23 |
| Exoc3 | 2.03E-27 | -0.3255063 | 0.024 | 0.166 | 5.49E-23 |
| Cask | 2.65E-27 | -0.4085366 | 0.016 | 0.147 | 7.16E-23 |
| Btbd9 | 2.69E-27 | -0.3354631 | 0.01 | 0.134 | 7.28E-23 |
| Dusp3 | 3.12E-27 | -0.3421467 | 0.029 | 0.172 | 8.43E-23 |
| Plxnd1 | 3.24E-27 | -0.3334871 | 0.017 | 0.148 | 8.77E-23 |
| Igsf8 | 3.85E-27 | -0.3996001 | 0.004 | 0.116 | 1.04E-22 |
| Cope | 3.86E-27 | -0.2562251 | 0.03 | 0.177 | 1.04E-22 |
| Clasp2 | 3.88E-27 | -0.3951247 | 0.025 | 0.167 | 1.05E-22 |
| H3f3b | 4.40E-27 | -0.5227853 | 0.588 | 0.908 | 1.19E-22 |
| Asah1 | 4.64E-27 | -0.2675075 | 0.028 | 0.17 | 1.26E-22 |
| Dgcr6 | 5.66E-27 | -0.4117341 | 0.003 | 0.112 | 1.53E-22 |
| Ckb | 6.22E-27 | -0.3846078 | 0.009 | 0.126 | 1.68E-22 |
| Ube4b | 6.23E-27 | -0.4868302 | 0.017 | 0.147 | 1.68E-22 |
| Hivep3 | 6.88E-27 | -0.3776689 | 0.014 | 0.141 | 1.86E-22 |
| Mbd5 | 7.05E-27 | -0.6061714 | 0.01 | 0.13 | 1.91E-22 |
| Lama3 | 7.38E-27 | -0.6093246 | 0.004 | 0.115 | 2.00E-22 |
| Ppp1r18 | 8.13E-27 | -0.3504829 | 0.02 | 0.152 | 2.20E-22 |
| Cpq | 8.61E-27 | -0.6102969 | 0.01 | 0.129 | 2.33E-22 |
| Klf2 | 9.17E-27 | -0.5650376 | 0.037 | 0.189 | 2.48E-22 |
| Tma7 | 9.51E-27 | -0.3003354 | 0.026 | 0.166 | 2.57E-22 |
| Guk1 | 1.13E-26 | -0.3049101 | 0.022 | 0.155 | 3.06E-22 |
| Aatf | 1.28E-26 | -0.3593725 | 0.015 | 0.142 | 3.46E-22 |
| Rpl4 | 1.44E-26 | -0.3190634 | 0.555 | 0.919 | 3.91E-22 |
| Ppp4r3b | 1.60E-26 | -0.2935596 | 0.029 | 0.172 | 4.34E-22 |
| Snx17 | 1.72E-26 | -0.2560506 | 0.007 | 0.121 | 4.66E-22 |
| Plekha6 | 1.98E-26 | -0.3611893 | 0.005 | 0.116 | 5.37E-22 |
| Abhd16a | 2.30E-26 | -0.3892782 | 0.004 | 0.113 | 6.21E-22 |
| Loxl1 | 2.66E-26 | -0.3529113 | 0.01 | 0.127 | 7.20E-22 |
| Runx1 | 2.99E-26 | 1.47626482 | 0.646 | 0.768 | 8.08E-22 |
| Rnf217 | 3.11E-26 | -0.3140777 | 0.033 | 0.18 | 8.41E-22 |
| Nfatc2 | 3.19E-26 | -0.459662 | 0.015 | 0.139 | 8.64E-22 |
| Sacs | 3.47E-26 | -0.3612233 | 0.025 | 0.162 | 9.37E-22 |
| Ift27 | 3.80E-26 | -0.2555262 | 0.012 | 0.134 | 1.03E-21 |
| Diaph2 | 3.82E-26 | -0.2831126 | 0.049 | 0.211 | 1.03E-21 |
| Crybg1 | 4.40E-26 | -0.4126655 | 0.029 | 0.167 | 1.19E-21 |
| Gk5 | 5.18E-26 | -0.4417927 | 0.015 | 0.139 | 1.40E-21 |
| 1110065P20I | 5.59E-26 | -0.3195675 | 0.008 | 0.12 | 1.51E-21 |
| Ugp2 | 5.72E-26 | -0.2937099 | 0.032 | 0.177 | 1.55E-21 |
| Robo1 | 6.32E-26 | -0.5157056 | 0.007 | 0.118 | 1.71E-21 |
| Mgll | 6.80E-26 | -0.3436236 | 0.025 | 0.158 | 1.84E-21 |
| Tpr | 6.97E-26 | 0.25544639 | 0.169 | 0.442 | 1.89E-21 |

| | | | | | |
|---|---|---|---|---|---|
| Zfyve9 | 7.05E-26 | -0.2655595 | 0.054 | 0.22 | 1.91E-21 |
| Zfp771 | 7.25E-26 | -0.2515774 | 0.012 | 0.133 | 1.96E-21 |
| Morf4l2 | 7.60E-26 | 0.28051082 | 0.134 | 0.383 | 2.06E-21 |
| Peak1 | 7.78E-26 | -0.4637584 | 0.037 | 0.187 | 2.10E-21 |
| Dld | 8.99E-26 | -0.3081679 | 0.014 | 0.136 | 2.43E-21 |
| Vapa | 9.11E-26 | 0.3107507 | 0.176 | 0.454 | 2.46E-21 |
| Tmem14c | 1.20E-25 | -0.3037655 | 0.016 | 0.139 | 3.24E-21 |
| Mcc | 1.40E-25 | -0.6120157 | 0.019 | 0.146 | 3.80E-21 |
| Tm2d2 | 1.48E-25 | -0.3420658 | 0.012 | 0.13 | 4.01E-21 |
| Orai1 | 1.76E-25 | -0.3919514 | 0.013 | 0.131 | 4.75E-21 |
| Hspb11 | 1.84E-25 | -0.3495451 | 0.014 | 0.135 | 4.98E-21 |
| Tdp2 | 1.85E-25 | -0.3716571 | 0.016 | 0.139 | 5.00E-21 |
| Dtymk | 2.03E-25 | -0.2685871 | 0.016 | 0.139 | 5.48E-21 |
| Phf20 | 2.05E-25 | -0.4500994 | 0.032 | 0.173 | 5.54E-21 |
| Gtf2i | 2.79E-25 | -0.2888299 | 0.027 | 0.162 | 7.54E-21 |
| Mrps16 | 3.00E-25 | -0.3061231 | 0.009 | 0.121 | 8.11E-21 |
| Arid1b | 3.22E-25 | -0.3662909 | 0.07 | 0.246 | 8.70E-21 |
| Brd3 | 3.27E-25 | -0.3107682 | 0.032 | 0.171 | 8.83E-21 |
| Bub3 | 3.48E-25 | -0.3342586 | 0.009 | 0.119 | 9.40E-21 |
| Scd2 | 3.64E-25 | -0.3212461 | 0.013 | 0.13 | 9.84E-21 |
| Bex3 | 4.37E-25 | -0.2618887 | 0.021 | 0.148 | 1.18E-20 |
| Nucks1 | 5.84E-25 | 0.36797599 | 0.152 | 0.405 | 1.58E-20 |
| Micu2 | 6.01E-25 | -0.3535331 | 0.003 | 0.105 | 1.63E-20 |
| Clcn4 | 6.35E-25 | -0.3067627 | 0.024 | 0.155 | 1.72E-20 |
| Insig1 | 6.48E-25 | -0.2567688 | 0.021 | 0.149 | 1.75E-20 |
| Cstf3 | 6.85E-25 | -0.2955097 | 0.021 | 0.149 | 1.85E-20 |
| Chmp2a | 7.03E-25 | 0.26013096 | 0.15 | 0.404 | 1.90E-20 |
| Kcnn3 | 7.49E-25 | -0.3950734 | 0.012 | 0.128 | 2.02E-20 |
| Sdc2 | 7.53E-25 | -0.3810094 | 0.023 | 0.151 | 2.04E-20 |
| Evi5 | 7.71E-25 | -0.3422515 | 0.023 | 0.151 | 2.08E-20 |
| Gphn | 8.33E-25 | -1.4881106 | 0.032 | 0.168 | 2.25E-20 |
| Rabl6 | 8.35E-25 | -0.3709359 | 0.016 | 0.136 | 2.26E-20 |
| Smad5 | 8.38E-25 | -0.2742025 | 0.022 | 0.149 | 2.27E-20 |
| Kmt2b | 8.46E-25 | -0.3186046 | 0.009 | 0.119 | 2.29E-20 |
| Ehmt2 | 9.42E-25 | -0.3331224 | 0.009 | 0.118 | 2.55E-20 |
| Vav3 | 1.01E-24 | -0.534775 | 0.041 | 0.188 | 2.73E-20 |
| Ptgfrn | 1.04E-24 | -0.2658811 | 0.028 | 0.161 | 2.80E-20 |
| Myod1 | 1.17E-24 | 1.87281242 | 0.336 | 0.188 | 3.16E-20 |
| Pgp | 1.22E-24 | -0.3608889 | 0.008 | 0.115 | 3.30E-20 |
| Musk | 1.57E-24 | -0.2655759 | 0.041 | 0.189 | 4.24E-20 |
| Tfg | 1.75E-24 | -0.2515471 | 0.018 | 0.14 | 4.74E-20 |
| Uchl3 | 1.87E-24 | -0.3113591 | 0.006 | 0.11 | 5.04E-20 |
| Jph1 | 1.99E-24 | -0.3755076 | 0.038 | 0.183 | 5.38E-20 |

| | | | | | |
|---|---|---|---|---|---|
| Rcan2 | 2.06E-24 | -0.2604821 | 0.08 | 0.262 | 5.56E-20 |
| Pard3 | 2.10E-24 | -0.2603529 | 0.042 | 0.189 | 5.67E-20 |
| Armcx2 | 2.49E-24 | -0.4339374 | 0.008 | 0.114 | 6.73E-20 |
| Slc43a2 | 2.56E-24 | -0.3860823 | 0.006 | 0.109 | 6.93E-20 |
| Prex1 | 2.77E-24 | -0.3254325 | 0.01 | 0.122 | 7.50E-20 |
| Naca | 2.85E-24 | -0.3252954 | 0.518 | 0.91 | 7.71E-20 |
| Fzd4 | 3.11E-24 | -0.255718 | 0.013 | 0.128 | 8.41E-20 |
| Palmd | 3.21E-24 | -0.3265314 | 0.031 | 0.167 | 8.67E-20 |
| Pafah1b1 | 3.25E-24 | 0.25759426 | 0.231 | 0.555 | 8.79E-20 |
| Tsen34 | 3.28E-24 | -0.3460609 | 0.01 | 0.118 | 8.88E-20 |
| Arpc5l | 3.41E-24 | -0.2604493 | 0.016 | 0.133 | 9.21E-20 |
| Vps28 | 3.65E-24 | -0.2805002 | 0.021 | 0.144 | 9.88E-20 |
| Hras | 3.88E-24 | -0.2719346 | 0.009 | 0.116 | 1.05E-19 |
| Fam124a | 4.06E-24 | -0.3145458 | 0.03 | 0.163 | 1.10E-19 |
| Ssbp2 | 4.27E-24 | -0.2817186 | 0.055 | 0.213 | 1.16E-19 |
| Fer | 4.28E-24 | -0.4012828 | 0.013 | 0.126 | 1.16E-19 |
| Kank2 | 4.38E-24 | -0.3161524 | 0.013 | 0.127 | 1.18E-19 |
| Snd1 | 4.45E-24 | -0.2613663 | 0.018 | 0.139 | 1.20E-19 |
| Nfu1 | 4.64E-24 | -0.3347123 | 0.004 | 0.104 | 1.26E-19 |
| Sox6 | 6.69E-24 | -0.7446649 | 0.012 | 0.124 | 1.81E-19 |
| Rpl28 | 6.79E-24 | -0.3260283 | 0.807 | 0.92 | 1.84E-19 |
| Commd10 | 7.44E-24 | -0.6553172 | 0.003 | 0.1 | 2.01E-19 |
| Ythdc2 | 7.45E-24 | -0.2607941 | 0.017 | 0.136 | 2.01E-19 |
| Map3k4 | 8.35E-24 | -0.2970654 | 0.007 | 0.11 | 2.26E-19 |
| Snap23 | 8.70E-24 | -0.3736914 | 0.018 | 0.136 | 2.35E-19 |
| 4833420G17 | 8.85E-24 | -0.3388822 | 0.006 | 0.106 | 2.39E-19 |
| St3gal2 | 8.89E-24 | -0.3438628 | 0.023 | 0.147 | 2.40E-19 |
| Nbea | 9.22E-24 | -0.3707226 | 0.028 | 0.157 | 2.49E-19 |
| Xist | 9.33E-24 | 1.35056086 | 0.448 | 0.289 | 2.52E-19 |
| Dctn3 | 9.60E-24 | -0.3233653 | 0.022 | 0.145 | 2.60E-19 |
| Lamp2 | 9.77E-24 | 0.37547806 | 0.235 | 0.562 | 2.64E-19 |
| Mtdh | 9.94E-24 | 0.33287664 | 0.206 | 0.503 | 2.69E-19 |
| Fem1b | 1.03E-23 | -0.3135125 | 0.013 | 0.126 | 2.79E-19 |
| Fam172a | 1.20E-23 | -0.5185916 | 0.008 | 0.112 | 3.24E-19 |
| Hmgn2 | 1.22E-23 | -0.316765 | 0.005 | 0.104 | 3.29E-19 |
| Plat | 1.29E-23 | -0.4928085 | 0.014 | 0.127 | 3.48E-19 |
| Tifa | 1.38E-23 | -0.4103438 | 0.01 | 0.118 | 3.72E-19 |
| Fam114a1 | 1.61E-23 | -0.2761193 | 0.008 | 0.111 | 4.35E-19 |
| Tspan12 | 1.70E-23 | -0.3971239 | 0.007 | 0.108 | 4.61E-19 |
| Itm2c | 1.78E-23 | -0.3486359 | 0.008 | 0.11 | 4.80E-19 |
| Tmem106b | 1.82E-23 | -0.3346779 | 0.009 | 0.114 | 4.93E-19 |
| Dph3 | 1.85E-23 | 0.32250012 | 0.136 | 0.371 | 5.01E-19 |
| Lamtor5 | 1.97E-23 | -0.310224 | 0.02 | 0.138 | 5.32E-19 |

| | | | | | |
|---|---|---|---|---|---|
| Smarcd3 | 1.99E-23 | -0.2726162 | 0.012 | 0.121 | 5.37E-19 |
| Mrc2 | 2.61E-23 | -0.4762245 | 0.004 | 0.1 | 7.05E-19 |
| Vps29 | 2.84E-23 | -0.2502181 | 0.026 | 0.151 | 7.68E-19 |
| Xpo7 | 3.14E-23 | -0.3702611 | 0.012 | 0.121 | 8.49E-19 |
| Ndufa2 | 3.71E-23 | 0.37605424 | 0.134 | 0.364 | 1.00E-18 |
| C1qbp | 4.28E-23 | 0.29721806 | 0.123 | 0.339 | 1.16E-18 |
| C330007P06I | 4.46E-23 | -0.2693112 | 0.011 | 0.119 | 1.21E-18 |
| Nectin1 | 4.54E-23 | -0.2987584 | 0.014 | 0.125 | 1.23E-18 |
| Gopc | 5.11E-23 | -0.3674462 | 0.006 | 0.105 | 1.38E-18 |
| Syne1 | 5.26E-23 | -0.3559969 | 0.035 | 0.17 | 1.42E-18 |
| Fau | 5.27E-23 | 0.57274447 | 0.902 | 0.923 | 1.42E-18 |
| Smad2 | 5.44E-23 | -0.4145464 | 0.01 | 0.114 | 1.47E-18 |
| Mast4 | 5.70E-23 | 0.3628226 | 0.139 | 0.369 | 1.54E-18 |
| Eci1 | 6.33E-23 | -0.3213814 | 0.013 | 0.122 | 1.71E-18 |
| Tmem51 | 6.68E-23 | -0.4224351 | 0.007 | 0.106 | 1.81E-18 |
| Rps18 | 7.15E-23 | -0.3452362 | 0.827 | 0.925 | 1.93E-18 |
| Ppdpf | 8.10E-23 | -0.2828366 | 0.007 | 0.106 | 2.19E-18 |
| G0s2 | 8.29E-23 | -0.8085942 | 0.019 | 0.135 | 2.24E-18 |
| Serinc5 | 8.72E-23 | -0.3280543 | 0.013 | 0.121 | 2.36E-18 |
| Anapc13 | 8.90E-23 | -0.2506292 | 0.024 | 0.145 | 2.41E-18 |
| Uqcrq | 9.64E-23 | 0.41760335 | 0.136 | 0.365 | 2.61E-18 |
| Vkorc1 | 1.04E-22 | -0.2677206 | 0.019 | 0.133 | 2.82E-18 |
| Eif3j1 | 1.13E-22 | 2.13774821 | 0.402 | 0.31 | 3.05E-18 |
| Atxn1 | 1.19E-22 | -0.4549875 | 0.01 | 0.115 | 3.21E-18 |
| Mdk | 1.23E-22 | -0.4275957 | 0.01 | 0.112 | 3.34E-18 |
| Tmem57 | 1.43E-22 | -0.2618117 | 0.022 | 0.14 | 3.87E-18 |
| Crispld2 | 1.44E-22 | -0.4140289 | 0.021 | 0.137 | 3.90E-18 |
| Prdx1 | 1.49E-22 | 0.44995562 | 0.225 | 0.543 | 4.02E-18 |
| Ogdh | 1.58E-22 | -0.3284787 | 0.032 | 0.161 | 4.27E-18 |
| Stoml2 | 1.69E-22 | -0.2576517 | 0.005 | 0.1 | 4.56E-18 |
| Gabpb1 | 2.00E-22 | -0.3163346 | 0.022 | 0.139 | 5.40E-18 |
| Trpc1 | 2.12E-22 | -0.3366405 | 0.012 | 0.117 | 5.74E-18 |
| Myo9a | 2.26E-22 | -0.2544196 | 0.046 | 0.188 | 6.11E-18 |
| Srsf10 | 2.39E-22 | 0.3001811 | 0.125 | 0.343 | 6.45E-18 |
| Cd9 | 2.58E-22 | 0.28008858 | 0.175 | 0.433 | 6.97E-18 |
| Rcn1 | 2.68E-22 | -0.2567313 | 0.014 | 0.122 | 7.25E-18 |
| Ddost | 2.78E-22 | -0.2755731 | 0.012 | 0.116 | 7.52E-18 |
| Prrc1 | 2.89E-22 | -0.2939681 | 0.007 | 0.104 | 7.81E-18 |
| Tmed4 | 2.94E-22 | -0.325298 | 0.014 | 0.122 | 7.96E-18 |
| Thoc2 | 3.93E-22 | 0.26205943 | 0.113 | 0.32 | 1.06E-17 |
| Cox6b1 | 4.01E-22 | 0.25899065 | 0.147 | 0.38 | 1.08E-17 |
| Tipin | 4.13E-22 | -0.3022632 | 0.009 | 0.107 | 1.12E-17 |
| Phlda1 | 4.42E-22 | -0.4768993 | 0.118 | 0.309 | 1.20E-17 |

| | | | | | |
|---|---|---|---|---|---|
| Tmed2 | 4.85E-22 | 0.40683064 | 0.13 | 0.348 | 1.31E-17 |
| Nipbl | 5.70E-22 | 0.25157898 | 0.237 | 0.559 | 1.54E-17 |
| Clk1 | 6.19E-22 | 0.27071925 | 0.273 | 0.627 | 1.67E-17 |
| Stat5b | 7.93E-22 | -0.2559874 | 0.029 | 0.149 | 2.14E-17 |
| Ago2 | 8.25E-22 | 0.28766991 | 0.125 | 0.336 | 2.23E-17 |
| Irf8 | 1.04E-21 | -0.2657933 | 0.023 | 0.137 | 2.82E-17 |
| Nol3 | 1.09E-21 | -0.2666516 | 0.009 | 0.106 | 2.95E-17 |
| Nxn | 1.12E-21 | -0.4128581 | 0.012 | 0.115 | 3.03E-17 |
| Rps14 | 1.14E-21 | -0.2861857 | 0.845 | 0.919 | 3.09E-17 |
| Hmox1 | 1.19E-21 | 2.50225731 | 0.196 | 0.069 | 3.22E-17 |
| Cwc27 | 1.24E-21 | -0.2622651 | 0.014 | 0.12 | 3.36E-17 |
| Timp3 | 1.24E-21 | -0.455772 | 0.13 | 0.33 | 3.36E-17 |
| Spr | 1.26E-21 | -0.2790817 | 0.011 | 0.113 | 3.42E-17 |
| Lbx1 | 1.64E-21 | -0.3070777 | 0.008 | 0.103 | 4.44E-17 |
| Nedd8 | 1.68E-21 | 0.2770564 | 0.097 | 0.284 | 4.53E-17 |
| Eea1 | 1.68E-21 | 0.2880411 | 0.109 | 0.304 | 4.55E-17 |
| Scd1 | 1.80E-21 | -0.296409 | 0.021 | 0.133 | 4.86E-17 |
| Tmem147 | 1.82E-21 | -0.3078162 | 0.01 | 0.108 | 4.92E-17 |
| Inpp5f | 1.91E-21 | -0.2720461 | 0.015 | 0.12 | 5.16E-17 |
| Fto | 2.32E-21 | -0.254626 | 0.057 | 0.206 | 6.27E-17 |
| Stbd1 | 2.35E-21 | -0.3562265 | 0.008 | 0.101 | 6.36E-17 |
| Ndufs3 | 2.51E-21 | -0.2699571 | 0.031 | 0.154 | 6.80E-17 |
| Dock1 | 2.62E-21 | -0.2917526 | 0.022 | 0.135 | 7.08E-17 |
| Hist1h2bc | 2.70E-21 | -0.4138183 | 0.074 | 0.232 | 7.31E-17 |
| Naa35 | 2.87E-21 | -0.2666809 | 0.007 | 0.1 | 7.75E-17 |
| Ccny | 3.26E-21 | -0.2744056 | 0.01 | 0.109 | 8.81E-17 |
| Adamts10 | 3.63E-21 | -0.323137 | 0.021 | 0.132 | 9.82E-17 |
| Exoc6b | 4.52E-21 | -0.4901257 | 0.023 | 0.136 | 1.22E-16 |
| Ccnl2 | 4.67E-21 | 0.25238587 | 0.157 | 0.395 | 1.26E-16 |
| Cmip | 4.82E-21 | -0.2843718 | 0.008 | 0.101 | 1.30E-16 |
| Cdkn2c | 5.31E-21 | -0.2663965 | 0.008 | 0.101 | 1.44E-16 |
| B2m | 5.85E-21 | 0.30839524 | 0.201 | 0.481 | 1.58E-16 |
| Cul1 | 7.60E-21 | 0.31647418 | 0.122 | 0.328 | 2.06E-16 |
| Zfp24 | 8.10E-21 | -0.2741815 | 0.008 | 0.1 | 2.19E-16 |
| Hk2 | 8.60E-21 | 2.19339443 | 0.271 | 0.141 | 2.32E-16 |
| Zfp652 | 9.73E-21 | -0.2933335 | 0.013 | 0.114 | 2.63E-16 |
| Hspb1 | 1.02E-20 | -1.1745281 | 0.508 | 0.708 | 2.75E-16 |
| Sae1 | 1.04E-20 | -0.2779055 | 0.009 | 0.102 | 2.82E-16 |
| Adcy2 | 1.23E-20 | -0.8896542 | 0.026 | 0.137 | 3.32E-16 |
| Scaf11 | 1.33E-20 | 0.37516188 | 0.165 | 0.411 | 3.59E-16 |
| Pde7b | 1.45E-20 | -0.9485555 | 0.032 | 0.15 | 3.92E-16 |
| Arhgap28 | 1.77E-20 | -0.2915677 | 0.035 | 0.16 | 4.78E-16 |
| Psmd13 | 1.99E-20 | -0.2848873 | 0.012 | 0.109 | 5.39E-16 |

| | | | | | |
|---|---|---|---|---|---|
| Gpx3 | 2.22E-20 | -0.3124681 | 0.63 | 0.94 | 5.99E-16 |
| B3galt1 | 2.29E-20 | -0.6981797 | 0.014 | 0.114 | 6.18E-16 |
| Cxcl2 | 2.37E-20 | -0.7271938 | 0.019 | 0.123 | 6.42E-16 |
| Ywhab | 2.38E-20 | 0.40642524 | 0.103 | 0.289 | 6.43E-16 |
| Pfkfb3 | 2.43E-20 | -0.2937549 | 0.009 | 0.101 | 6.57E-16 |
| Rab28 | 2.45E-20 | -0.2661093 | 0.011 | 0.107 | 6.64E-16 |
| Zfand3 | 2.59E-20 | -0.2632504 | 0.029 | 0.145 | 6.99E-16 |
| Gde1 | 3.24E-20 | -0.2614636 | 0.009 | 0.1 | 8.75E-16 |
| Anxa5 | 3.75E-20 | 0.30601331 | 0.194 | 0.455 | 1.01E-15 |
| Dip2c | 3.87E-20 | -0.3269838 | 0.077 | 0.233 | 1.05E-15 |
| Rap1gds1 | 4.05E-20 | -0.296969 | 0.024 | 0.134 | 1.09E-15 |
| Dbi | 4.25E-20 | 0.32355645 | 0.089 | 0.258 | 1.15E-15 |
| Psd3 | 4.30E-20 | -0.4121641 | 0.011 | 0.106 | 1.16E-15 |
| Grasp | 5.14E-20 | -0.253261 | 0.022 | 0.129 | 1.39E-15 |
| Myl6 | 5.91E-20 | 0.30605886 | 0.33 | 0.713 | 1.60E-15 |
| Arhgef7 | 6.04E-20 | -0.356938 | 0.013 | 0.109 | 1.63E-15 |
| Tubgcp4 | 6.93E-20 | -0.3906538 | 0.014 | 0.112 | 1.87E-15 |
| Rab10 | 7.17E-20 | 0.26506858 | 0.153 | 0.379 | 1.94E-15 |
| Trim35 | 7.78E-20 | -0.3077389 | 0.024 | 0.131 | 2.10E-15 |
| Phpt1 | 9.96E-20 | -0.2734191 | 0.01 | 0.1 | 2.69E-15 |
| Trap1 | 1.17E-19 | -0.2529018 | 0.013 | 0.107 | 3.16E-15 |
| Dcn | 1.22E-19 | 2.01760478 | 0.162 | 0.049 | 3.30E-15 |
| Hmg20a | 1.23E-19 | -0.3919628 | 0.01 | 0.1 | 3.31E-15 |
| Arcn1 | 1.38E-19 | 0.29783616 | 0.113 | 0.304 | 3.74E-15 |
| Stag2 | 1.60E-19 | 0.25234433 | 0.097 | 0.273 | 4.32E-15 |
| Canx | 1.65E-19 | 0.3530861 | 0.126 | 0.329 | 4.45E-15 |
| Rbbp4 | 1.69E-19 | 0.33772586 | 0.092 | 0.265 | 4.58E-15 |
| Eif1 | 1.72E-19 | 0.57333252 | 0.832 | 0.921 | 4.64E-15 |
| Csgalnact1 | 1.77E-19 | -0.5193133 | 0.015 | 0.112 | 4.78E-15 |
| Cttn | 1.85E-19 | -0.2833019 | 0.014 | 0.109 | 5.00E-15 |
| Pold4 | 1.97E-19 | 0.35142721 | 0.09 | 0.259 | 5.34E-15 |
| Igf1 | 2.04E-19 | -0.3316514 | 0.012 | 0.106 | 5.52E-15 |
| Sesn1 | 2.11E-19 | -0.287674 | 0.013 | 0.108 | 5.70E-15 |
| Fbxw11 | 2.20E-19 | -0.4376782 | 0.013 | 0.107 | 5.95E-15 |
| Prpf39 | 2.24E-19 | 0.29589298 | 0.138 | 0.345 | 6.07E-15 |
| Selenok | 2.37E-19 | 0.31698226 | 0.239 | 0.539 | 6.42E-15 |
| Ghr | 2.39E-19 | -0.5472431 | 0.025 | 0.131 | 6.46E-15 |
| Cdkal1 | 2.59E-19 | -0.3138123 | 0.026 | 0.135 | 7.00E-15 |
| Cst3 | 3.07E-19 | 0.61553442 | 0.203 | 0.478 | 8.29E-15 |
| Prox1 | 3.10E-19 | 0.31587648 | 0.206 | 0.477 | 8.39E-15 |
| Srp9 | 3.25E-19 | 0.25300202 | 0.118 | 0.313 | 8.78E-15 |
| Ndufs1 | 3.52E-19 | -0.3369386 | 0.019 | 0.119 | 9.51E-15 |
| Tacc1 | 3.73E-19 | 0.43545039 | 0.132 | 0.334 | 1.01E-14 |

| | | | | | |
|---|---|---|---|---|---|
| Erp44 | 4.11E-19 | -0.2645034 | 0.011 | 0.103 | 1.11E-14 |
| Epb41l3 | 5.36E-19 | -0.3072907 | 0.013 | 0.106 | 1.45E-14 |
| Nr3c2 | 5.76E-19 | -0.6615022 | 0.02 | 0.12 | 1.56E-14 |
| Phldb2 | 6.20E-19 | -0.343895 | 0.014 | 0.107 | 1.68E-14 |
| Ist1 | 6.30E-19 | 0.25261058 | 0.101 | 0.276 | 1.70E-14 |
| Brd2 | 6.45E-19 | 0.35683684 | 0.202 | 0.467 | 1.74E-14 |
| Nop53 | 6.50E-19 | 0.34467831 | 0.227 | 0.524 | 1.76E-14 |
| Atrnl1 | 7.36E-19 | -0.6910915 | 0.016 | 0.111 | 1.99E-14 |
| Hnrnph1 | 7.53E-19 | 0.31627183 | 0.288 | 0.631 | 2.04E-14 |
| Fbxl17 | 8.81E-19 | -0.7841692 | 0.011 | 0.1 | 2.38E-14 |
| Rps25 | 9.53E-19 | 0.67775666 | 0.808 | 0.912 | 2.58E-14 |
| Mef2a | 1.03E-18 | 0.25519964 | 0.116 | 0.305 | 2.79E-14 |
| Rps27l | 1.13E-18 | 0.40453937 | 0.166 | 0.391 | 3.06E-14 |
| Smchd1 | 1.14E-18 | 0.3135759 | 0.125 | 0.319 | 3.09E-14 |
| Arhgap12 | 1.27E-18 | -0.2656798 | 0.012 | 0.103 | 3.43E-14 |
| Ik | 1.33E-18 | 0.28628284 | 0.101 | 0.273 | 3.59E-14 |
| Egr3 | 1.35E-18 | -0.4196556 | 0.027 | 0.131 | 3.64E-14 |
| Coro1c | 1.35E-18 | -0.2567703 | 0.021 | 0.121 | 3.66E-14 |
| Nap1l5 | 1.45E-18 | -0.2766435 | 0.024 | 0.126 | 3.92E-14 |
| Cdc27 | 1.80E-18 | -0.267548 | 0.018 | 0.115 | 4.87E-14 |
| Stip1 | 2.01E-18 | 0.50734199 | 0.164 | 0.391 | 5.43E-14 |
| Psmd4 | 2.38E-18 | 0.41895053 | 0.117 | 0.304 | 6.45E-14 |
| Atp9b | 2.59E-18 | -0.2722571 | 0.036 | 0.15 | 6.99E-14 |
| Sbds | 2.60E-18 | 0.26393817 | 0.121 | 0.311 | 7.02E-14 |
| Cdk8 | 3.48E-18 | -0.7550946 | 0.02 | 0.116 | 9.42E-14 |
| Cmss1 | 3.55E-18 | -2.6031344 | 0.028 | 0.129 | 9.59E-14 |
| Tshr | 4.40E-18 | -0.4455928 | 0.015 | 0.106 | 1.19E-13 |
| Ntn1 | 4.65E-18 | -0.264048 | 0.014 | 0.105 | 1.26E-13 |
| Frem1 | 4.98E-18 | -0.3871902 | 0.015 | 0.106 | 1.35E-13 |
| Arhgap6 | 5.15E-18 | -0.614828 | 0.038 | 0.151 | 1.39E-13 |
| Clint1 | 5.21E-18 | 0.31534644 | 0.09 | 0.253 | 1.41E-13 |
| Fosl2 | 5.34E-18 | 0.31330058 | 0.223 | 0.496 | 1.44E-13 |
| Rps24 | 6.66E-18 | 0.39591899 | 0.938 | 0.929 | 1.80E-13 |
| 4932438A13 | 7.17E-18 | 0.27613296 | 0.185 | 0.421 | 1.94E-13 |
| Slk | 7.53E-18 | 0.2722525 | 0.105 | 0.276 | 2.04E-13 |
| Arid5b | 8.32E-18 | 1.39296283 | 0.552 | 0.574 | 2.25E-13 |
| Mbnl1 | 9.30E-18 | 0.36835634 | 0.186 | 0.424 | 2.51E-13 |
| F2r | 9.50E-18 | 0.38260839 | 0.151 | 0.367 | 2.57E-13 |
| Cisd2 | 1.07E-17 | 0.35235095 | 0.1 | 0.267 | 2.90E-13 |
| Ccl7 | 1.09E-17 | -0.7294024 | 0.015 | 0.105 | 2.94E-13 |
| Sdc4 | 1.19E-17 | -0.4836259 | 0.621 | 0.893 | 3.23E-13 |
| Abca8a | 1.22E-17 | 0.53329472 | 0.137 | 0.337 | 3.30E-13 |
| Tomm7 | 1.30E-17 | 0.57276691 | 0.13 | 0.322 | 3.51E-13 |

| | | | | | |
|---|---|---|---|---|---|
| Aplp2 | 1.33E-17 | 0.32204369 | 0.162 | 0.373 | 3.60E-13 |
| Cd302 | 1.44E-17 | 0.46903616 | 0.104 | 0.272 | 3.89E-13 |
| Tmcc1 | 1.56E-17 | -0.3782262 | 0.038 | 0.15 | 4.21E-13 |
| Cavin1 | 1.74E-17 | 0.28375228 | 0.09 | 0.248 | 4.72E-13 |
| Hspa8 | 1.90E-17 | -0.391072 | 0.875 | 0.926 | 5.15E-13 |
| Ntn4 | 1.98E-17 | 1.66330005 | 0.206 | 0.091 | 5.36E-13 |
| Rora | 2.08E-17 | -0.8653824 | 0.256 | 0.522 | 5.64E-13 |
| Dennd1a | 2.19E-17 | -0.5040207 | 0.02 | 0.114 | 5.93E-13 |
| Tgfb3 | 2.23E-17 | -0.2849652 | 0.014 | 0.101 | 6.03E-13 |
| Ncor2 | 2.34E-17 | 0.33758249 | 0.088 | 0.241 | 6.32E-13 |
| Ilf2 | 2.91E-17 | 0.37254805 | 0.143 | 0.343 | 7.87E-13 |
| Ndufa7 | 3.23E-17 | 0.41976675 | 0.15 | 0.359 | 8.73E-13 |
| Cep112 | 3.41E-17 | -0.2871484 | 0.017 | 0.107 | 9.22E-13 |
| Psmd7 | 3.85E-17 | 0.26746956 | 0.073 | 0.217 | 1.04E-12 |
| Lnpep | 3.87E-17 | 0.3150119 | 0.139 | 0.336 | 1.05E-12 |
| Csnk1g1 | 3.91E-17 | -0.2696091 | 0.018 | 0.109 | 1.06E-12 |
| Esd | 3.93E-17 | 0.38801806 | 0.092 | 0.249 | 1.06E-12 |
| Sp3 | 4.19E-17 | 0.29911272 | 0.096 | 0.259 | 1.13E-12 |
| Setbp1 | 4.33E-17 | -0.5517888 | 0.031 | 0.134 | 1.17E-12 |
| Ddx3x | 4.60E-17 | 0.34790625 | 0.304 | 0.661 | 1.24E-12 |
| Map7d1 | 4.64E-17 | 0.31714536 | 0.115 | 0.295 | 1.26E-12 |
| Ankrd11 | 4.66E-17 | 0.31458769 | 0.255 | 0.556 | 1.26E-12 |
| Ranbp1 | 4.72E-17 | 0.3187333 | 0.133 | 0.324 | 1.28E-12 |
| Rpl23 | 4.74E-17 | 0.39726867 | 0.956 | 0.934 | 1.28E-12 |
| Ndufb2 | 4.79E-17 | 0.37794656 | 0.091 | 0.246 | 1.29E-12 |
| Bdp1 | 5.32E-17 | 0.3658684 | 0.08 | 0.229 | 1.44E-12 |
| Slc7a2 | 6.79E-17 | -0.2548102 | 0.026 | 0.123 | 1.84E-12 |
| Csrp2 | 7.59E-17 | -0.2774096 | 0.027 | 0.124 | 2.05E-12 |
| Gabarapl2 | 1.00E-16 | 0.28700877 | 0.125 | 0.302 | 2.71E-12 |
| Actr3 | 1.11E-16 | 0.30717666 | 0.077 | 0.217 | 3.00E-12 |
| Marcks | 1.17E-16 | 0.31301481 | 0.134 | 0.324 | 3.17E-12 |
| Cd47 | 1.26E-16 | 0.35221551 | 0.173 | 0.398 | 3.42E-12 |
| Eif4g1 | 1.34E-16 | 0.39881802 | 0.183 | 0.413 | 3.63E-12 |
| Atxn7l3b | 1.52E-16 | 0.37761019 | 0.139 | 0.333 | 4.10E-12 |
| Aff4 | 1.84E-16 | 0.32589651 | 0.23 | 0.5 | 4.98E-12 |
| Bzw1 | 1.87E-16 | 0.34066148 | 0.135 | 0.327 | 5.06E-12 |
| Atrx | 1.99E-16 | 0.38186376 | 0.29 | 0.621 | 5.37E-12 |
| Zfp36l2 | 2.22E-16 | 0.37581434 | 0.299 | 0.614 | 6.02E-12 |
| Clic4 | 2.28E-16 | 0.26793353 | 0.3 | 0.619 | 6.18E-12 |
| Safb2 | 2.34E-16 | 0.36424914 | 0.138 | 0.33 | 6.32E-12 |
| Eif4e | 3.02E-16 | 0.36339223 | 0.116 | 0.288 | 8.17E-12 |
| Kdm1a | 3.62E-16 | 0.37527552 | 0.074 | 0.214 | 9.78E-12 |
| Rapgef6 | 3.75E-16 | 0.2539202 | 0.118 | 0.291 | 1.01E-11 |

| | | | | | |
|---|---|---|---|---|---|
| Spop | 4.28E-16 | 0.42461465 | 0.185 | 0.418 | 1.16E-11 |
| Spag9 | 4.32E-16 | 0.34789178 | 0.227 | 0.491 | 1.17E-11 |
| Abl2 | 4.42E-16 | -0.2500041 | 0.019 | 0.106 | 1.20E-11 |
| Snrpe | 4.48E-16 | 0.44079975 | 0.122 | 0.299 | 1.21E-11 |
| Pkn2 | 4.97E-16 | 0.37749332 | 0.148 | 0.343 | 1.34E-11 |
| G3bp1 | 5.16E-16 | 0.32623841 | 0.098 | 0.256 | 1.40E-11 |
| Plpp3 | 5.42E-16 | 0.4750146 | 0.13 | 0.306 | 1.46E-11 |
| Rad23b | 5.53E-16 | 0.27727411 | 0.113 | 0.282 | 1.50E-11 |
| Pcf11 | 5.79E-16 | 0.48856175 | 0.158 | 0.368 | 1.56E-11 |
| Wdr26 | 9.04E-16 | 0.38057423 | 0.11 | 0.275 | 2.45E-11 |
| Mrpl33 | 1.15E-15 | 0.48738209 | 0.1 | 0.253 | 3.11E-11 |
| Cox7c | 1.19E-15 | 0.48306131 | 0.236 | 0.514 | 3.21E-11 |
| Taf1d | 1.23E-15 | 0.34266003 | 0.17 | 0.383 | 3.32E-11 |
| Slc6a6 | 1.53E-15 | 0.29764103 | 0.123 | 0.295 | 4.15E-11 |
| Hspa9 | 1.73E-15 | 0.41833745 | 0.16 | 0.367 | 4.68E-11 |
| Kdm5a | 1.94E-15 | 0.37897914 | 0.123 | 0.293 | 5.24E-11 |
| Sfpq | 1.99E-15 | 0.27954038 | 0.297 | 0.625 | 5.39E-11 |
| Tnrc6b | 2.12E-15 | 0.36906155 | 0.191 | 0.419 | 5.74E-11 |
| Ctnnb1 | 2.44E-15 | 0.31378868 | 0.09 | 0.235 | 6.59E-11 |
| Ankhd1 | 3.81E-15 | 0.28616635 | 0.116 | 0.279 | 1.03E-10 |
| Ost4 | 3.91E-15 | 0.33729587 | 0.088 | 0.228 | 1.06E-10 |
| Zfp280d | 4.05E-15 | 0.27466574 | 0.089 | 0.232 | 1.10E-10 |
| Ppp1r10 | 4.19E-15 | 0.26470948 | 0.097 | 0.248 | 1.13E-10 |
| Xpr1 | 4.30E-15 | 0.38529837 | 0.137 | 0.32 | 1.16E-10 |
| Rbm8a | 4.66E-15 | 0.44845948 | 0.18 | 0.391 | 1.26E-10 |
| Cbx5 | 4.67E-15 | 0.31924583 | 0.069 | 0.197 | 1.26E-10 |
| Srsf6 | 4.97E-15 | 0.26762848 | 0.095 | 0.243 | 1.34E-10 |
| Pde4dip | 6.00E-15 | -0.5637978 | 0.024 | 0.109 | 1.62E-10 |
| Hsp90ab1 | 6.42E-15 | -0.4074174 | 0.903 | 0.943 | 1.74E-10 |
| Kcnt2 | 6.97E-15 | -0.5634084 | 0.023 | 0.108 | 1.89E-10 |
| Hipk1 | 8.92E-15 | 0.30852196 | 0.074 | 0.204 | 2.41E-10 |
| Zfpm2 | 9.10E-15 | -0.4339098 | 0.022 | 0.106 | 2.46E-10 |
| Ndrg1 | 1.12E-14 | 0.51918418 | 0.224 | 0.457 | 3.03E-10 |
| Tcf25 | 1.22E-14 | 0.57305052 | 0.205 | 0.449 | 3.29E-10 |
| Ddr2 | 1.42E-14 | 0.40035321 | 0.076 | 0.208 | 3.83E-10 |
| Hnrnpd | 1.52E-14 | 0.45400234 | 0.177 | 0.385 | 4.11E-10 |
| Ets2 | 1.73E-14 | 0.26918509 | 0.133 | 0.309 | 4.67E-10 |
| Irak1 | 1.73E-14 | 0.33132486 | 0.092 | 0.236 | 4.68E-10 |
| Igf1r | 1.77E-14 | 0.2714953 | 0.181 | 0.392 | 4.80E-10 |
| Pabpc1 | 1.85E-14 | 0.25605559 | 0.339 | 0.696 | 5.01E-10 |
| Ddx27 | 2.08E-14 | 0.26109054 | 0.055 | 0.169 | 5.62E-10 |
| Smad7 | 2.11E-14 | 0.35204193 | 0.11 | 0.259 | 5.70E-10 |
| Tcea1 | 2.38E-14 | 0.27458273 | 0.094 | 0.237 | 6.42E-10 |

| | | | | | |
|---|---|---|---|---|---|
| Crebbp | 2.42E-14 | 0.36998655 | 0.208 | 0.444 | 6.56E-10 |
| Cox7a2 | 2.54E-14 | 0.44950658 | 0.21 | 0.451 | 6.86E-10 |
| Cggbp1 | 2.62E-14 | 0.37847019 | 0.105 | 0.252 | 7.07E-10 |
| Rpl34 | 2.65E-14 | 0.47463031 | 0.845 | 0.913 | 7.15E-10 |
| Usp10 | 3.22E-14 | 0.27237088 | 0.058 | 0.172 | 8.70E-10 |
| Sf3b3 | 3.22E-14 | 0.38542654 | 0.104 | 0.256 | 8.72E-10 |
| AC160336.1 | 3.23E-14 | 1.77408258 | 0.181 | 0.084 | 8.75E-10 |
| Gpatch8 | 3.40E-14 | 0.26424476 | 0.095 | 0.237 | 9.20E-10 |
| Hdac4 | 3.60E-14 | 0.25533567 | 0.072 | 0.197 | 9.74E-10 |
| Eif1b | 4.09E-14 | 0.27949641 | 0.057 | 0.17 | 1.10E-09 |
| Flna | 5.97E-14 | 0.31947916 | 0.109 | 0.26 | 1.61E-09 |
| Samd4b | 6.72E-14 | 0.34640776 | 0.129 | 0.292 | 1.82E-09 |
| Hnrnph2 | 7.30E-14 | 0.48107155 | 0.097 | 0.24 | 1.97E-09 |
| Etf1 | 7.90E-14 | 0.47808077 | 0.197 | 0.417 | 2.14E-09 |
| Serinc1 | 9.29E-14 | 0.38526588 | 0.109 | 0.259 | 2.51E-09 |
| Taf7 | 1.05E-13 | 0.34123837 | 0.125 | 0.287 | 2.84E-09 |
| Slc38a2 | 1.24E-13 | 1.40466815 | 0.556 | 0.681 | 3.35E-09 |
| Stk40 | 1.32E-13 | 0.38347781 | 0.154 | 0.341 | 3.56E-09 |
| Igf2bp2 | 1.33E-13 | 1.60468225 | 0.145 | 0.056 | 3.60E-09 |
| Gng5 | 1.40E-13 | 0.37838508 | 0.272 | 0.561 | 3.79E-09 |
| Scn7a | 1.41E-13 | 1.53871627 | 0.174 | 0.081 | 3.82E-09 |
| Syncrip | 1.45E-13 | 0.46020231 | 0.22 | 0.459 | 3.92E-09 |
| Mrpl52 | 1.45E-13 | 0.60500978 | 0.169 | 0.369 | 3.93E-09 |
| Sub1 | 1.54E-13 | 0.3672263 | 0.16 | 0.345 | 4.16E-09 |
| Ppm1a | 1.68E-13 | 0.33305709 | 0.11 | 0.262 | 4.56E-09 |
| Atxn2 | 1.87E-13 | 0.26202946 | 0.052 | 0.158 | 5.05E-09 |
| Prkx | 2.00E-13 | 0.44477634 | 0.104 | 0.248 | 5.40E-09 |
| Twsg1 | 2.07E-13 | 0.31220586 | 0.08 | 0.206 | 5.60E-09 |
| Bcas2 | 2.09E-13 | 0.26721418 | 0.066 | 0.18 | 5.65E-09 |
| Elk4 | 2.16E-13 | 0.27330093 | 0.062 | 0.175 | 5.85E-09 |
| Aamp | 2.56E-13 | 0.3697698 | 0.091 | 0.224 | 6.93E-09 |
| Wapl | 2.65E-13 | 0.42897756 | 0.163 | 0.353 | 7.16E-09 |
| Rybp | 2.78E-13 | 0.33207405 | 0.083 | 0.213 | 7.51E-09 |
| Pcbp1 | 2.93E-13 | 0.48422931 | 0.165 | 0.357 | 7.92E-09 |
| Txnip | 2.95E-13 | 0.25413466 | 0.119 | 0.266 | 7.97E-09 |
| Hnrnpa0 | 3.17E-13 | 0.36093434 | 0.082 | 0.21 | 8.58E-09 |
| Ubap2l | 3.23E-13 | 0.42140314 | 0.162 | 0.355 | 8.75E-09 |
| Romo1 | 3.26E-13 | 0.31024493 | 0.076 | 0.199 | 8.81E-09 |
| Ddx21 | 3.50E-13 | 1.91859275 | 0.413 | 0.409 | 9.47E-09 |
| Elf2 | 3.74E-13 | 0.38571596 | 0.147 | 0.325 | 1.01E-08 |
| Rab5a | 4.24E-13 | 0.3137453 | 0.108 | 0.254 | 1.15E-08 |
| Mkln1 | 4.38E-13 | 0.41364261 | 0.157 | 0.338 | 1.18E-08 |
| Drp2 | 4.50E-13 | 0.38955368 | 0.129 | 0.289 | 1.22E-08 |

| | | | | | |
|---|---|---|---|---|---|
| Baz1a | 4.78E-13 | 0.32211189 | 0.113 | 0.263 | 1.29E-08 |
| Thoc7 | 5.07E-13 | 0.5360278 | 0.075 | 0.197 | 1.37E-08 |
| Supt5 | 5.31E-13 | 0.26582276 | 0.071 | 0.188 | 1.44E-08 |
| Rock2 | 5.53E-13 | 0.26374437 | 0.355 | 0.702 | 1.50E-08 |
| Acadl | 5.66E-13 | 0.3416645 | 0.07 | 0.184 | 1.53E-08 |
| Hnrnpc | 5.68E-13 | 0.60195201 | 0.248 | 0.517 | 1.54E-08 |
| Mrpl20 | 6.06E-13 | 0.25277109 | 0.062 | 0.171 | 1.64E-08 |
| Ssr3 | 6.31E-13 | 0.38768319 | 0.141 | 0.313 | 1.71E-08 |
| S100a11 | 6.41E-13 | 0.5172598 | 0.125 | 0.281 | 1.73E-08 |
| Medag | 6.51E-13 | 0.43441387 | 0.265 | 0.511 | 1.76E-08 |
| Trim56 | 6.82E-13 | 0.26954087 | 0.062 | 0.172 | 1.84E-08 |
| Atad2b | 7.69E-13 | 0.25870304 | 0.064 | 0.176 | 2.08E-08 |
| Xrn1 | 7.72E-13 | 0.53939132 | 0.083 | 0.21 | 2.09E-08 |
| Cwc15 | 7.86E-13 | 0.33385457 | 0.047 | 0.145 | 2.12E-08 |
| Il6st | 8.42E-13 | 0.41451735 | 0.156 | 0.339 | 2.28E-08 |
| Ncl | 8.66E-13 | 1.23824104 | 0.588 | 0.753 | 2.34E-08 |
| Zfp207 | 8.74E-13 | 0.41451844 | 0.128 | 0.289 | 2.36E-08 |
| Tardbp | 1.01E-12 | 0.37765019 | 0.107 | 0.252 | 2.73E-08 |
| Ubl5 | 1.03E-12 | 0.30252814 | 0.064 | 0.174 | 2.79E-08 |
| Fmc1 | 1.15E-12 | 0.25398493 | 0.066 | 0.177 | 3.11E-08 |
| Kmt2a | 1.36E-12 | 0.34725046 | 0.211 | 0.437 | 3.68E-08 |
| Pum1 | 1.39E-12 | 0.46962659 | 0.188 | 0.399 | 3.75E-08 |
| Pnisr | 1.39E-12 | 0.4519006 | 0.278 | 0.565 | 3.77E-08 |
| Dek | 1.42E-12 | 0.50931302 | 0.233 | 0.48 | 3.83E-08 |
| Dazap2 | 1.54E-12 | 0.31208056 | 0.105 | 0.244 | 4.15E-08 |
| Ipo7 | 1.69E-12 | 0.4409114 | 0.111 | 0.257 | 4.56E-08 |
| Chmp4b | 1.83E-12 | 0.51075699 | 0.147 | 0.315 | 4.94E-08 |
| Syap1 | 1.89E-12 | 0.33687133 | 0.068 | 0.179 | 5.10E-08 |
| Ube2n | 1.92E-12 | 0.34096975 | 0.069 | 0.18 | 5.20E-08 |
| Rps6kb1 | 2.21E-12 | 0.33440282 | 0.078 | 0.196 | 5.98E-08 |
| Mybbp1a | 2.26E-12 | 0.53848648 | 0.104 | 0.24 | 6.11E-08 |
| Sec63 | 2.61E-12 | 0.41911633 | 0.121 | 0.271 | 7.05E-08 |
| Rpl27a | 2.64E-12 | 0.38273022 | 0.915 | 0.929 | 7.13E-08 |
| Itgb1 | 2.92E-12 | 0.55415585 | 0.252 | 0.515 | 7.88E-08 |
| Magt1 | 3.48E-12 | 0.33274422 | 0.041 | 0.131 | 9.40E-08 |
| Cox6c | 3.67E-12 | 0.64827992 | 0.249 | 0.51 | 9.92E-08 |
| Txnrd1 | 3.93E-12 | 1.55056664 | 0.43 | 0.432 | 1.06E-07 |
| Gnai3 | 3.94E-12 | 0.43369039 | 0.105 | 0.24 | 1.07E-07 |
| Wac | 3.97E-12 | 0.42948665 | 0.182 | 0.38 | 1.07E-07 |
| Srrm1 | 4.01E-12 | 0.4849709 | 0.303 | 0.618 | 1.08E-07 |
| Sik1 | 4.50E-12 | 0.44685997 | 0.155 | 0.331 | 1.22E-07 |
| 11-Sep | 4.58E-12 | 0.30267893 | 0.077 | 0.194 | 1.24E-07 |
| Pds5a | 5.01E-12 | 0.28384399 | 0.066 | 0.173 | 1.36E-07 |

| | | | | | |
|---|---|---|---|---|---|
| Mat2a | 5.91E-12 | 0.33972468 | 0.3 | 0.574 | 1.60E-07 |
| Zfp91 | 6.55E-12 | 0.34174401 | 0.115 | 0.258 | 1.77E-07 |
| Zfc3h1 | 6.65E-12 | 0.26220588 | 0.073 | 0.186 | 1.80E-07 |
| R3hdm2 | 6.68E-12 | 0.25309058 | 0.085 | 0.206 | 1.81E-07 |
| Rps17 | 7.03E-12 | 0.61535023 | 0.734 | 0.88 | 1.90E-07 |
| H2afz | 8.83E-12 | 0.53450923 | 0.232 | 0.464 | 2.39E-07 |
| Ube2l3 | 9.73E-12 | 0.43201916 | 0.11 | 0.248 | 2.63E-07 |
| Ptbp1 | 1.06E-11 | 0.48653605 | 0.095 | 0.223 | 2.87E-07 |
| Pak2 | 1.07E-11 | 0.49684813 | 0.141 | 0.301 | 2.90E-07 |
| Hes1 | 1.18E-11 | 0.31065216 | 0.093 | 0.216 | 3.18E-07 |
| Ipo5 | 1.18E-11 | 0.29571415 | 0.081 | 0.197 | 3.20E-07 |
| Hsph1 | 1.19E-11 | -0.4197422 | 0.549 | 0.739 | 3.21E-07 |
| Snw1 | 1.29E-11 | 0.32930478 | 0.103 | 0.235 | 3.48E-07 |
| Txlna | 1.34E-11 | 0.26150541 | 0.073 | 0.185 | 3.62E-07 |
| Ube2k | 1.36E-11 | 0.4319197 | 0.122 | 0.267 | 3.67E-07 |
| Asb5 | 1.38E-11 | 2.0309977 | 0.434 | 0.444 | 3.74E-07 |
| Pdcd6ip | 1.48E-11 | 0.39425803 | 0.062 | 0.165 | 3.99E-07 |
| Kras | 1.54E-11 | 0.41222611 | 0.088 | 0.208 | 4.17E-07 |
| Cyth3 | 1.68E-11 | 0.32282992 | 0.114 | 0.252 | 4.56E-07 |
| Dnajc1 | 1.71E-11 | 0.38867282 | 0.12 | 0.265 | 4.61E-07 |
| Tcp1 | 1.76E-11 | 0.67816539 | 0.161 | 0.343 | 4.75E-07 |
| Utp3 | 1.76E-11 | 0.4991103 | 0.062 | 0.164 | 4.76E-07 |
| Kpna4 | 2.02E-11 | 0.29801246 | 0.086 | 0.204 | 5.45E-07 |
| Purb | 2.12E-11 | 0.58856786 | 0.155 | 0.325 | 5.72E-07 |
| Hdgf | 2.12E-11 | 0.53198757 | 0.106 | 0.239 | 5.73E-07 |
| Elob | 2.16E-11 | 0.58390436 | 0.215 | 0.436 | 5.84E-07 |
| Fxr1 | 2.39E-11 | 0.65225056 | 0.222 | 0.451 | 6.46E-07 |
| Med13l | 2.41E-11 | 0.44887399 | 0.167 | 0.343 | 6.52E-07 |
| Rab18 | 2.57E-11 | 0.52460411 | 0.204 | 0.409 | 6.95E-07 |
| Metap2 | 2.58E-11 | 0.77764749 | 0.173 | 0.356 | 6.98E-07 |
| Rell1 | 2.67E-11 | 0.26523986 | 0.053 | 0.147 | 7.21E-07 |
| Neat1 | 2.68E-11 | 0.34629529 | 0.354 | 0.656 | 7.24E-07 |
| Atp5l | 2.90E-11 | 0.60728295 | 0.29 | 0.571 | 7.85E-07 |
| Gtf2h5 | 3.17E-11 | 0.36484537 | 0.093 | 0.215 | 8.56E-07 |
| Prelp | 3.24E-11 | 0.34654129 | 0.133 | 0.281 | 8.77E-07 |
| Rif1 | 3.57E-11 | 0.39820743 | 0.095 | 0.217 | 9.66E-07 |
| Fbxl3 | 3.84E-11 | 0.29726569 | 0.059 | 0.157 | 1.04E-06 |
| Ywhag | 3.87E-11 | 0.34574412 | 0.07 | 0.176 | 1.05E-06 |
| Ankrd12 | 4.59E-11 | 0.6210995 | 0.123 | 0.263 | 1.24E-06 |
| Brd8 | 4.98E-11 | 0.26861887 | 0.059 | 0.157 | 1.35E-06 |
| Actr2 | 5.09E-11 | 0.28689758 | 0.062 | 0.161 | 1.38E-06 |
| Ensa | 5.59E-11 | 0.37392814 | 0.12 | 0.258 | 1.51E-06 |
| Ahctf1 | 5.63E-11 | 0.33478383 | 0.099 | 0.225 | 1.52E-06 |

| | | | | | |
|---|---|---|---|---|---|
| Cd44 | 6.48E-11 | 0.25020414 | 0.082 | 0.194 | 1.75E-06 |
| Sf1 | 7.28E-11 | 0.46324786 | 0.155 | 0.321 | 1.97E-06 |
| Ddx24 | 8.08E-11 | 0.6160282 | 0.152 | 0.314 | 2.18E-06 |
| Snrpc | 8.17E-11 | 0.28977813 | 0.047 | 0.132 | 2.21E-06 |
| Tgif1 | 8.47E-11 | 0.34648361 | 0.21 | 0.411 | 2.29E-06 |
| Serping1 | 8.49E-11 | 0.54574668 | 0.279 | 0.551 | 2.30E-06 |
| Ier3ip1 | 8.68E-11 | 0.43708046 | 0.079 | 0.19 | 2.35E-06 |
| Ubr2 | 9.28E-11 | 0.42418693 | 0.086 | 0.201 | 2.51E-06 |
| B230219D22 | 1.00E-10 | 0.53233546 | 0.104 | 0.229 | 2.71E-06 |
| Cpeb4 | 1.04E-10 | 0.32981606 | 0.07 | 0.173 | 2.81E-06 |
| Sf3b6 | 1.04E-10 | 0.42099924 | 0.078 | 0.186 | 2.81E-06 |
| Dnajc19 | 1.13E-10 | 0.37334156 | 0.046 | 0.13 | 3.07E-06 |
| Ptp4a2 | 1.14E-10 | 0.68320668 | 0.169 | 0.342 | 3.08E-06 |
| Hipk3 | 1.25E-10 | 0.58796087 | 0.115 | 0.251 | 3.38E-06 |
| Rtf1 | 1.36E-10 | 0.33812346 | 0.091 | 0.207 | 3.68E-06 |
| Ppp3ca | 1.37E-10 | 0.46540966 | 0.156 | 0.314 | 3.72E-06 |
| Utp11 | 1.45E-10 | 0.26158867 | 0.06 | 0.155 | 3.93E-06 |
| Tfdp2 | 1.51E-10 | 0.2779301 | 0.069 | 0.167 | 4.08E-06 |
| Snrpd1 | 2.23E-10 | 0.41933314 | 0.101 | 0.223 | 6.02E-06 |
| Atp5k | 2.42E-10 | 0.32410266 | 0.055 | 0.145 | 6.54E-06 |
| Pknox1 | 2.43E-10 | 0.37350918 | 0.097 | 0.216 | 6.57E-06 |
| Nupr1 | 2.48E-10 | 0.34185273 | 0.068 | 0.166 | 6.71E-06 |
| Chchd1 | 2.54E-10 | 0.32154761 | 0.068 | 0.165 | 6.86E-06 |
| Frmd6 | 2.70E-10 | 0.5141279 | 0.226 | 0.441 | 7.30E-06 |
| Zmat2 | 2.76E-10 | 0.44438528 | 0.084 | 0.193 | 7.45E-06 |
| Ypel5 | 2.77E-10 | 0.29492621 | 0.054 | 0.142 | 7.50E-06 |
| Mdm2 | 2.79E-10 | 0.25025617 | 0.06 | 0.151 | 7.54E-06 |
| Smarca5 | 2.89E-10 | 0.37181407 | 0.299 | 0.579 | 7.82E-06 |
| Lpar1 | 3.04E-10 | 0.51581129 | 0.112 | 0.24 | 8.22E-06 |
| Chic2 | 3.13E-10 | 0.66000694 | 0.184 | 0.367 | 8.46E-06 |
| Setd7 | 3.17E-10 | 0.33353666 | 0.062 | 0.154 | 8.58E-06 |
| Rpl7l1 | 3.24E-10 | 0.32433866 | 0.063 | 0.157 | 8.77E-06 |
| Ylpm1 | 3.71E-10 | 0.2567712 | 0.067 | 0.163 | 1.00E-05 |
| U2surp | 3.93E-10 | 0.63933165 | 0.138 | 0.287 | 1.06E-05 |
| P4ha1 | 3.95E-10 | 0.2918067 | 0.139 | 0.284 | 1.07E-05 |
| Rps26 | 3.98E-10 | 0.64644015 | 0.742 | 0.842 | 1.08E-05 |
| Hnrnpab | 4.18E-10 | 0.52997124 | 0.228 | 0.444 | 1.13E-05 |
| Mapre1 | 4.46E-10 | 0.68627761 | 0.174 | 0.346 | 1.21E-05 |
| Zfp655 | 4.49E-10 | 0.35298862 | 0.057 | 0.147 | 1.21E-05 |
| Spred1 | 4.77E-10 | 0.39036949 | 0.17 | 0.336 | 1.29E-05 |
| Fubp1 | 5.18E-10 | 0.68987679 | 0.246 | 0.48 | 1.40E-05 |
| Tia1 | 5.48E-10 | 0.32317835 | 0.066 | 0.161 | 1.48E-05 |
| Cdkn1a | 5.61E-10 | 0.44299148 | 0.159 | 0.314 | 1.52E-05 |

| | | | | | |
|---|---|---|---|---|---|
| Man2a1 | 5.66E-10 | 0.26143387 | 0.042 | 0.119 | 1.53E-05 |
| Prnp | 6.07E-10 | 0.37003453 | 0.056 | 0.145 | 1.64E-05 |
| Cebpb | 6.47E-10 | 0.52857326 | 0.237 | 0.456 | 1.75E-05 |
| Slc16a2 | 6.81E-10 | 0.2501241 | 0.048 | 0.129 | 1.84E-05 |
| Set | 7.38E-10 | 0.48365248 | 0.238 | 0.463 | 1.99E-05 |
| Map2k3 | 7.77E-10 | 0.32508906 | 0.09 | 0.198 | 2.10E-05 |
| Eif5 | 7.92E-10 | 0.31680011 | 0.378 | 0.709 | 2.14E-05 |
| Arl6ip1 | 8.14E-10 | 0.28849563 | 0.075 | 0.176 | 2.20E-05 |
| Cep170 | 8.51E-10 | 0.29398414 | 0.05 | 0.134 | 2.30E-05 |
| Twistnb | 9.59E-10 | 0.45221387 | 0.085 | 0.193 | 2.59E-05 |
| 1810037I17R | 1.14E-09 | 0.64532521 | 0.139 | 0.277 | 3.09E-05 |
| Ube2v2 | 1.21E-09 | 0.38291086 | 0.071 | 0.168 | 3.26E-05 |
| Rbm6 | 1.28E-09 | 0.42889231 | 0.094 | 0.207 | 3.45E-05 |
| Snhg6 | 1.32E-09 | 0.41247179 | 0.079 | 0.179 | 3.57E-05 |
| Rpl31 | 1.33E-09 | 0.78808887 | 0.636 | 0.842 | 3.61E-05 |
| Nfyb | 1.38E-09 | 0.3478355 | 0.11 | 0.23 | 3.74E-05 |
| Gstm1 | 1.38E-09 | 0.51629978 | 0.254 | 0.487 | 3.74E-05 |
| Sdcbp | 1.61E-09 | 0.35170407 | 0.089 | 0.192 | 4.35E-05 |
| Rab1a | 1.69E-09 | 0.50290914 | 0.103 | 0.219 | 4.58E-05 |
| Nfkbib | 1.76E-09 | 0.35393977 | 0.074 | 0.173 | 4.76E-05 |
| Atp2b1 | 1.87E-09 | 0.57802858 | 0.183 | 0.35 | 5.06E-05 |
| Tm9sf4 | 1.94E-09 | 0.38141038 | 0.065 | 0.156 | 5.25E-05 |
| Mob1a | 2.11E-09 | 0.31475958 | 0.061 | 0.149 | 5.71E-05 |
| Nmt2 | 2.19E-09 | 0.3736442 | 0.063 | 0.152 | 5.92E-05 |
| Xrn2 | 2.39E-09 | 0.48041631 | 0.14 | 0.282 | 6.47E-05 |
| Dicer1 | 2.40E-09 | 0.28833124 | 0.069 | 0.161 | 6.48E-05 |
| Gpx1 | 2.62E-09 | 0.43735439 | 0.073 | 0.167 | 7.09E-05 |
| Fgfr1op2 | 2.67E-09 | 0.505098 | 0.108 | 0.225 | 7.21E-05 |
| Naa15 | 2.70E-09 | 0.67129816 | 0.15 | 0.297 | 7.30E-05 |
| Iqgap1 | 2.85E-09 | 0.59322011 | 0.19 | 0.366 | 7.70E-05 |
| Rsbn1 | 2.86E-09 | 0.29477259 | 0.071 | 0.164 | 7.73E-05 |
| Myd88 | 3.09E-09 | 0.30615433 | 0.085 | 0.186 | 8.36E-05 |
| Mysm1 | 3.43E-09 | 0.25413281 | 0.059 | 0.145 | 9.28E-05 |
| Psma1 | 3.59E-09 | 0.30047106 | 0.054 | 0.136 | 9.71E-05 |
| Ergic2 | 3.82E-09 | 0.36896166 | 0.091 | 0.199 | 0.00010342 |
| Yme1l1 | 3.93E-09 | 0.3335059 | 0.079 | 0.177 | 0.0001063 |
| Esyt2 | 3.93E-09 | 0.45595595 | 0.15 | 0.295 | 0.00010633 |
| Usp36 | 4.07E-09 | 0.35368224 | 0.065 | 0.154 | 0.00010994 |
| Nasp | 4.16E-09 | 0.37771296 | 0.099 | 0.211 | 0.00011251 |
| Gls | 4.19E-09 | 0.62033517 | 0.168 | 0.326 | 0.00011319 |
| Klf9 | 4.27E-09 | 0.31460797 | 0.392 | 0.712 | 0.00011555 |
| Tcp11l2 | 4.34E-09 | 0.29006113 | 0.063 | 0.149 | 0.00011725 |
| Scaf8 | 4.45E-09 | 0.29842631 | 0.061 | 0.147 | 0.00012031 |

| | | | | | |
|---|---|---|---|---|---|
| Abi1 | 4.97E-09 | 0.47049882 | 0.091 | 0.198 | 0.00013432 |
| Tnpo3 | 5.05E-09 | 0.32072605 | 0.062 | 0.148 | 0.00013654 |
| Sod1 | 5.41E-09 | 0.55858987 | 0.202 | 0.384 | 0.00014642 |
| Fas | 5.45E-09 | 0.55664619 | 0.149 | 0.291 | 0.00014736 |
| Chmp3 | 5.92E-09 | 0.40882674 | 0.056 | 0.138 | 0.00015998 |
| Hnrnpk | 5.96E-09 | 0.59898967 | 0.317 | 0.618 | 0.00016114 |
| Srsf1 | 6.21E-09 | 0.37510091 | 0.098 | 0.207 | 0.00016806 |
| Utp14a | 6.65E-09 | 0.32236347 | 0.088 | 0.188 | 0.00017994 |
| Nfe2l2 | 7.01E-09 | 0.48923984 | 0.126 | 0.25 | 0.00018946 |
| Arpp19 | 7.26E-09 | 0.26709405 | 0.052 | 0.131 | 0.0001964 |
| Arfrp1 | 7.44E-09 | 0.30520013 | 0.038 | 0.107 | 0.00020123 |
| Zbtb44 | 7.70E-09 | 0.4243664 | 0.069 | 0.157 | 0.00020818 |
| Wdr45b | 8.57E-09 | 0.36338985 | 0.062 | 0.146 | 0.00023175 |
| Serbp1 | 8.95E-09 | 1.22196793 | 0.502 | 0.656 | 0.00024201 |
| Kmt2d | 9.17E-09 | 0.26676181 | 0.049 | 0.124 | 0.00024809 |
| Nufip2 | 1.02E-08 | 0.61148346 | 0.168 | 0.322 | 0.00027485 |
| Mafg | 1.04E-08 | 0.43682614 | 0.118 | 0.239 | 0.0002807 |
| Clip1 | 1.06E-08 | 0.63850037 | 0.148 | 0.289 | 0.00028699 |
| Snrpd3 | 1.10E-08 | 0.4716735 | 0.083 | 0.178 | 0.00029635 |
| Rpl30 | 1.10E-08 | 0.34709321 | 0.867 | 0.921 | 0.00029846 |
| Ivns1abp | 1.22E-08 | 0.63370199 | 0.145 | 0.283 | 0.00032928 |
| Tax1bp1 | 1.29E-08 | 0.64040601 | 0.277 | 0.528 | 0.00034785 |
| mt-Nd5 | 1.30E-08 | 0.26127234 | 0.412 | 0.775 | 0.0003525 |
| Naf1 | 1.31E-08 | 0.49278137 | 0.069 | 0.157 | 0.00035482 |
| Plpbp | 1.35E-08 | 0.43649475 | 0.107 | 0.219 | 0.00036546 |
| Dusp11 | 1.39E-08 | 0.57848257 | 0.124 | 0.248 | 0.00037527 |
| Rab8b | 1.61E-08 | 0.26586425 | 0.041 | 0.109 | 0.00043622 |
| Slc25a5 | 1.70E-08 | 0.63760425 | 0.25 | 0.463 | 0.00045914 |
| Pofut2 | 1.77E-08 | 0.44109399 | 0.068 | 0.155 | 0.00047813 |
| Hnrnpu | 1.78E-08 | 0.46398441 | 0.374 | 0.712 | 0.00048029 |
| Tpp2 | 1.83E-08 | 0.34113218 | 0.09 | 0.189 | 0.00049427 |
| Fam133b | 1.85E-08 | 0.56203115 | 0.127 | 0.25 | 0.00050125 |
| Rbmxl1 | 1.89E-08 | 0.40022429 | 0.078 | 0.17 | 0.00051216 |
| Aff1 | 2.11E-08 | 0.30251057 | 0.109 | 0.217 | 0.00057182 |
| Rb1cc1 | 2.27E-08 | 0.62270069 | 0.179 | 0.338 | 0.00061502 |
| Dpm3 | 2.32E-08 | 0.33819113 | 0.055 | 0.132 | 0.0006285 |
| Abhd2 | 2.35E-08 | 0.57451866 | 0.07 | 0.158 | 0.00063569 |
| Pan3 | 2.63E-08 | 0.29880504 | 0.066 | 0.15 | 0.00071011 |
| Gnl2 | 2.72E-08 | 0.3860765 | 0.071 | 0.159 | 0.00073565 |
| Smad1 | 2.92E-08 | 0.38507431 | 0.09 | 0.187 | 0.00079089 |
| Mia3 | 3.39E-08 | 0.28534535 | 0.073 | 0.16 | 0.00091678 |
| Fnbp4 | 3.45E-08 | 0.55321205 | 0.109 | 0.219 | 0.00093237 |
| Gabarap | 3.50E-08 | 0.4991913 | 0.357 | 0.678 | 0.00094519 |

| | | | | | |
|---|---|---|---|---|---|
| Tnnt3 | 3.56E-08 | 0.4065584 | 0.172 | 0.31 | 0.00096189 |
| Cdk11b | 3.63E-08 | 0.55770458 | 0.17 | 0.321 | 0.00098167 |
| Ep300 | 3.69E-08 | 0.4282555 | 0.102 | 0.208 | 0.00099696 |
| Ccdc59 | 3.71E-08 | 0.64109126 | 0.143 | 0.278 | 0.00100381 |
| Rcn2 | 4.42E-08 | 0.48765385 | 0.072 | 0.158 | 0.00119504 |
| Kmt5b | 4.46E-08 | 0.28771642 | 0.07 | 0.156 | 0.00120705 |
| Slco2a1 | 4.52E-08 | 1.32268602 | 0.127 | 0.065 | 0.00122222 |
| Got2 | 4.68E-08 | 0.26412889 | 0.058 | 0.133 | 0.00126652 |
| Denr | 4.95E-08 | 0.29618969 | 0.067 | 0.147 | 0.00133764 |
| Cstb | 5.20E-08 | 0.58497433 | 0.07 | 0.154 | 0.00140681 |
| Fcho2 | 5.84E-08 | 0.51308484 | 0.139 | 0.268 | 0.0015801 |
| Klf4 | 6.07E-08 | 0.50791625 | 0.269 | 0.463 | 0.00164166 |
| Rbm27 | 6.15E-08 | 0.48359489 | 0.084 | 0.177 | 0.00166427 |
| Smndc1 | 6.56E-08 | 0.45258606 | 0.088 | 0.183 | 0.00177284 |
| Rdx | 6.65E-08 | 0.61195959 | 0.181 | 0.338 | 0.00179944 |
| Casp4 | 7.09E-08 | 0.287747 | 0.063 | 0.142 | 0.00191671 |
| Arid4b | 7.30E-08 | 0.56610802 | 0.25 | 0.459 | 0.00197525 |
| Epb41l4aos | 7.65E-08 | 0.46860314 | 0.101 | 0.203 | 0.00206914 |
| Nap1l4 | 8.48E-08 | 0.26390151 | 0.069 | 0.149 | 0.0022923 |
| Rbbp6 | 8.88E-08 | 0.67543966 | 0.215 | 0.399 | 0.00240102 |
| Nup98 | 9.03E-08 | 0.56485403 | 0.205 | 0.377 | 0.00244073 |
| Mapk8ip3 | 9.20E-08 | 0.34120109 | 0.056 | 0.131 | 0.00248891 |
| Nme1 | 9.25E-08 | 0.46603214 | 0.095 | 0.194 | 0.00250179 |
| Lsm3 | 9.35E-08 | 0.33505139 | 0.055 | 0.129 | 0.00252959 |
| Galnt1 | 1.06E-07 | 0.30748278 | 0.052 | 0.123 | 0.00286868 |
| Vps13a | 1.08E-07 | 0.36257776 | 0.061 | 0.137 | 0.00292029 |
| Brix1 | 1.10E-07 | 0.33140662 | 0.057 | 0.131 | 0.00296143 |
| Pole4 | 1.10E-07 | 0.48966481 | 0.071 | 0.156 | 0.00296581 |
| Pcgf5 | 1.10E-07 | 0.58069483 | 0.131 | 0.252 | 0.00296836 |
| Afdn | 1.12E-07 | 0.4258703 | 0.061 | 0.137 | 0.00302042 |
| Gatad2a | 1.16E-07 | 0.31448055 | 0.047 | 0.115 | 0.0031493 |
| Rps23 | 1.19E-07 | 0.33362071 | 0.919 | 0.924 | 0.00321044 |
| Cpsf6 | 1.19E-07 | 0.32728498 | 0.065 | 0.144 | 0.00321212 |
| Eif4e2 | 1.28E-07 | 0.32714158 | 0.068 | 0.146 | 0.00346965 |
| Fmr1 | 1.31E-07 | 0.61951645 | 0.125 | 0.24 | 0.00355178 |
| Msn | 1.36E-07 | 0.79357347 | 0.212 | 0.391 | 0.00366547 |
| Polr3d | 1.44E-07 | 0.44649916 | 0.064 | 0.142 | 0.00389983 |
| Cse1l | 1.50E-07 | 0.25733116 | 0.04 | 0.103 | 0.00405052 |
| Psmc3 | 1.52E-07 | 0.41696133 | 0.097 | 0.193 | 0.00411112 |
| Zfp281 | 1.62E-07 | 0.37434307 | 0.05 | 0.118 | 0.00439383 |
| Suco | 1.66E-07 | 0.4582738 | 0.087 | 0.178 | 0.00448121 |
| Nop14 | 1.71E-07 | 0.52842906 | 0.185 | 0.343 | 0.00462959 |
| Son | 1.73E-07 | 0.99104297 | 0.559 | 0.77 | 0.00466767 |

| | | | | | |
|---|---|---|---|---|---|
| Phf10 | 1.74E-07 | 0.30692046 | 0.042 | 0.105 | 0.00471368 |
| Tmem256 | 1.76E-07 | 0.46048219 | 0.092 | 0.185 | 0.00475047 |
| Col4a3bp | 1.77E-07 | 0.36118456 | 0.048 | 0.115 | 0.00479661 |
| Cep83 | 1.80E-07 | 0.33342012 | 0.045 | 0.11 | 0.00486231 |
| Gtpbp4 | 1.81E-07 | 0.66615899 | 0.162 | 0.301 | 0.00489904 |
| Sypl | 1.82E-07 | 0.36109808 | 0.07 | 0.151 | 0.00491721 |
| Hprt | 1.87E-07 | 0.34992046 | 0.062 | 0.136 | 0.0050457 |
| Srp19 | 1.88E-07 | 0.37730251 | 0.041 | 0.104 | 0.00507409 |
| Krit1 | 1.89E-07 | 0.34452446 | 0.072 | 0.154 | 0.00511179 |
| Timm9 | 1.91E-07 | 0.33515219 | 0.054 | 0.126 | 0.00517068 |
| Brd7 | 1.95E-07 | 0.34031858 | 0.05 | 0.119 | 0.00526937 |
| Etnk1 | 2.20E-07 | 0.31767184 | 0.083 | 0.169 | 0.00594676 |
| Hnrnpl | 2.21E-07 | 0.59321323 | 0.113 | 0.221 | 0.0059631 |
| Ctnna1 | 2.24E-07 | 0.66993679 | 0.157 | 0.289 | 0.00606441 |
| Nop58 | 2.25E-07 | 0.84479682 | 0.569 | 0.703 | 0.00609743 |
| Clcf1 | 2.26E-07 | 0.32088924 | 0.057 | 0.129 | 0.0061028 |
| Riok3 | 2.30E-07 | 0.49407003 | 0.089 | 0.177 | 0.00622507 |
| Nav2 | 2.32E-07 | 0.520097 | 0.096 | 0.191 | 0.00627908 |
| Gpbp1l1 | 2.33E-07 | 0.30241546 | 0.063 | 0.137 | 0.00629118 |
| Api5 | 2.33E-07 | 0.30433494 | 0.043 | 0.106 | 0.00630465 |
| Syf2 | 2.44E-07 | 0.60199076 | 0.114 | 0.222 | 0.00660731 |
| Ftl1 | 2.46E-07 | 0.31599912 | 0.399 | 0.706 | 0.00665929 |
| Rnpc3 | 2.47E-07 | 0.32195422 | 0.06 | 0.132 | 0.00667006 |
| Eif4g2 | 2.52E-07 | 0.49726519 | 0.376 | 0.71 | 0.0068208 |
| Tmpo | 2.60E-07 | 0.45617671 | 0.094 | 0.187 | 0.00702321 |
| Gm2000 | 2.65E-07 | 0.36355886 | 0.068 | 0.145 | 0.00716806 |
| Pdk4 | 2.69E-07 | 1.41798463 | 0.191 | 0.132 | 0.00726626 |
| Lsm12 | 2.77E-07 | 0.46064477 | 0.074 | 0.157 | 0.00749595 |
| Xbp1 | 2.99E-07 | 0.52245655 | 0.164 | 0.297 | 0.00808465 |
| Sec61a1 | 3.00E-07 | 0.32666233 | 0.067 | 0.143 | 0.00810203 |
| Srsf2 | 3.26E-07 | 0.60009071 | 0.228 | 0.411 | 0.00881633 |
| Tank | 3.28E-07 | 0.45181019 | 0.076 | 0.158 | 0.00888128 |
| Abcf1 | 3.30E-07 | 0.7174286 | 0.112 | 0.218 | 0.00891369 |
| Nsrp1 | 3.45E-07 | 0.58174816 | 0.081 | 0.166 | 0.0093202 |
| Mfsd14a | 3.45E-07 | 0.3986798 | 0.062 | 0.136 | 0.00932713 |
| Rbm28 | 3.49E-07 | 0.55893139 | 0.074 | 0.157 | 0.00942541 |
| Sqstm1 | 3.54E-07 | 0.34653963 | 0.403 | 0.697 | 0.00958146 |
| Atp1b3 | 3.58E-07 | 0.5644645 | 0.097 | 0.192 | 0.00967796 |
| Sec62 | 3.78E-07 | 0.82799679 | 0.246 | 0.447 | 0.01021359 |
| Ilf3 | 3.82E-07 | 0.30314251 | 0.07 | 0.148 | 0.01031755 |
| Cnot1 | 3.85E-07 | 0.5417154 | 0.104 | 0.204 | 0.01040909 |
| Rbm18 | 4.01E-07 | 0.31595051 | 0.061 | 0.133 | 0.01084049 |
| Tra2a | 4.11E-07 | 0.54058203 | 0.328 | 0.607 | 0.01111158 |

| | | | | | |
|---|---|---|---|---|---|
| Gar1 | 4.40E-07 | 0.31608041 | 0.059 | 0.129 | 0.01190887 |
| Cfh | 4.52E-07 | 0.56979854 | 0.258 | 0.456 | 0.01221989 |
| 2410089E03I | 4.54E-07 | 0.26111797 | 0.043 | 0.104 | 0.01227415 |
| Pafah1b2 | 4.83E-07 | 0.27683591 | 0.049 | 0.114 | 0.01305164 |
| Nab1 | 4.99E-07 | 0.2934267 | 0.053 | 0.12 | 0.01350256 |
| Zfp593 | 5.01E-07 | 0.29355339 | 0.05 | 0.116 | 0.01355454 |
| Dkc1 | 5.31E-07 | 0.34290284 | 0.053 | 0.121 | 0.01434992 |
| Snrnp70 | 5.89E-07 | 0.79278785 | 0.277 | 0.51 | 0.01593905 |
| Ppig | 5.97E-07 | 0.85592047 | 0.188 | 0.342 | 0.01614913 |
| Tsix | 5.98E-07 | 1.50630111 | 0.152 | 0.096 | 0.01615953 |
| Oser1 | 6.91E-07 | 0.27844144 | 0.045 | 0.106 | 0.01869049 |
| Tns1 | 7.18E-07 | 0.39065875 | 0.07 | 0.144 | 0.01940387 |
| Osbpl8 | 7.34E-07 | 0.25245077 | 0.063 | 0.134 | 0.019838 |
| Tmsb10 | 8.53E-07 | 0.70590184 | 0.27 | 0.489 | 0.02305409 |
| Arglu1 | 8.94E-07 | 0.73494009 | 0.203 | 0.369 | 0.02416482 |
| Erh | 8.98E-07 | 0.28676998 | 0.05 | 0.113 | 0.02427131 |
| Slc4a7 | 9.25E-07 | 0.57663116 | 0.116 | 0.22 | 0.02501275 |
| Ttc14 | 9.48E-07 | 0.38670293 | 0.096 | 0.185 | 0.0256245 |
| Uap1 | 9.80E-07 | 0.44847465 | 0.087 | 0.17 | 0.02650453 |
| Rpl21 | 1.08E-06 | 0.29270738 | 0.902 | 0.923 | 0.02929044 |
| Nrd1 | 1.15E-06 | 0.56571948 | 0.13 | 0.24 | 0.03120619 |
| mt-Nd3 | 1.22E-06 | 0.54489151 | 0.273 | 0.481 | 0.03300526 |
| Ubxn4 | 1.26E-06 | 0.58342505 | 0.288 | 0.518 | 0.03404576 |
| Sarnp | 1.26E-06 | 0.73929182 | 0.175 | 0.32 | 0.03419243 |
| Dnaja1 | 1.27E-06 | 0.61479164 | 0.688 | 0.804 | 0.03423222 |
| Tug1 | 1.55E-06 | 0.38264125 | 0.051 | 0.115 | 0.041998 |
| Wls | 1.65E-06 | 0.39264579 | 0.095 | 0.18 | 0.04473059 |

**Table S2: Differentially expressed transcripts between caFGFR1+ and caFGFR1- myonuclear clusters**

| | p_val | avg_log2FC | pct.1 | pct.2 | p_val_adj |
|---|---|---|---|---|---|
| Atp6v1h | 0 | 0.98516622 | 0.214 | 0.053 | 0 |
| Myl1 | 0 | -2.8877256 | 0.485 | 0.865 | 0 |
| Tns1 | 0 | 0.83828503 | 0.396 | 0.158 | 0 |
| Hdac4 | 0 | 0.65167488 | 0.476 | 0.206 | 0 |
| Ptpn4 | 0 | 0.57111502 | 0.359 | 0.147 | 0 |
| Glul | 0 | 1.11739961 | 0.152 | 0.02 | 0 |
| Ptpn14 | 0 | 0.65620334 | 0.269 | 0.094 | 0 |
| Pnpla7 | 0 | 0.99969128 | 0.345 | 0.12 | 0 |
| Ubr3 | 0 | 0.59297056 | 0.474 | 0.22 | 0 |
| Map3k20 | 0 | 0.53409837 | 0.478 | 0.223 | 0 |
| Ttn | 0 | -0.3678802 | 0.982 | 0.988 | 0 |
| Cry2 | 0 | 0.94420089 | 0.154 | 0.029 | 0 |
| Pla2g4e | 0 | 0.63286236 | 0.412 | 0.18 | 0 |
| Zc3h6 | 0 | 1.07104985 | 0.204 | 0.041 | 0 |
| Napb | 0 | 1.00173282 | 0.262 | 0.068 | 0 |
| Tmem189 | 0 | 1.04861637 | 0.174 | 0.035 | 0 |
| Xist | 0 | 0.67127546 | 0.463 | 0.207 | 0 |
| Amot | 0 | 0.92844768 | 0.196 | 0.044 | 0 |
| Slc7a11 | 0 | 1.01112109 | 0.18 | 0.039 | 0 |
| Foxo1 | 0 | 1.10893653 | 0.405 | 0.144 | 0 |
| Rorc | 0 | 1.00443954 | 0.16 | 0.028 | 0 |
| Txnip | 0 | 0.75068643 | 0.247 | 0.079 | 0 |
| Zranb2 | 0 | 0.98635204 | 0.204 | 0.047 | 0 |
| Wwp1 | 0 | 0.97567536 | 0.333 | 0.104 | 0 |
| Abca1 | 0 | 1.08761616 | 0.192 | 0.039 | 0 |
| Ecpas | 0 | 0.60131806 | 0.377 | 0.156 | 0 |
| Trim63 | 0 | 2.91625097 | 0.709 | 0.221 | 0 |
| Ddi2 | 0 | 0.8057431 | 0.294 | 0.103 | 0 |
| Vps13d | 0 | 0.58011292 | 0.292 | 0.109 | 0 |
| Cd36 | 0 | 1.11505534 | 0.319 | 0.112 | 0 |
| Kmt2c | 0 | 0.56367074 | 0.41 | 0.175 | 0 |
| Dnajb6 | 0 | 0.70486975 | 0.393 | 0.158 | 0 |
| Art3 | 0 | 0.78784393 | 0.337 | 0.134 | 0 |
| Myo18b | 0 | 1.0882946 | 0.696 | 0.342 | 0 |
| Acacb | 0 | 1.02003968 | 0.534 | 0.208 | 0 |
| Clip1 | 0 | 1.13586414 | 0.7 | 0.336 | 0 |
| Phkg1 | 0 | -1.4126374 | 0.507 | 0.68 | 0 |
| Cdk8 | 0 | -0.9238051 | 0.743 | 0.758 | 0 |
| Pdk4 | 0 | 2.00705187 | 0.242 | 0.01 | 0 |
| Lncpint | 0 | 1.03806184 | 0.811 | 0.461 | 0 |
| Tmem140 | 0 | 0.99295424 | 0.156 | 0.027 | 0 |

| | | | | | |
|---|---|---|---|---|---|
| Aldh1l1 | 0 | 1.19995003 | 0.137 | 0.011 | 0 |
| 9530026P05I | 0 | 1.72590741 | 0.377 | 0.088 | 0 |
| Ybx3 | 0 | 1.46939478 | 0.473 | 0.11 | 0 |
| Mybpc2 | 0 | -2.1643879 | 0.156 | 0.394 | 0 |
| Fgfr2 | 0 | 0.98082011 | 0.232 | 0.062 | 0 |
| Tacc2 | 0 | 1.19018599 | 0.882 | 0.61 | 0 |
| Zranb1 | 0 | 0.84386637 | 0.304 | 0.095 | 0 |
| Syne1 | 0 | 1.30093963 | 0.459 | 0.146 | 0 |
| Sesn1 | 0 | 0.985055 | 0.35 | 0.111 | 0 |
| Cep85l | 0 | 1.19102898 | 0.463 | 0.146 | 0 |
| Tmtc2 | 0 | 0.7271326 | 0.275 | 0.099 | 0 |
| Slc7a2 | 0 | 0.84614268 | 0.192 | 0.046 | 0 |
| Pdlim3 | 0 | 0.52056547 | 0.443 | 0.214 | 0 |
| Ankrd11 | 0 | 0.69110828 | 0.64 | 0.32 | 0 |
| Ptprg | 0 | 0.62638284 | 0.63 | 0.342 | 0 |
| Gm48099 | 0 | -1.8588746 | 0.464 | 0.549 | 0 |
| Zmiz1 | 0 | 0.65106195 | 0.338 | 0.129 | 0 |
| Zcchc24 | 0 | 1.08320293 | 0.188 | 0.035 | 0 |
| Asb14 | 0 | -1.3201228 | 0.467 | 0.592 | 0 |
| Ube4a | 0 | 1.15898613 | 0.275 | 0.069 | 0 |
| Peak1 | 0 | -1.2979358 | 0.786 | 0.858 | 0 |
| Tpm1 | 0 | -1.0332739 | 0.499 | 0.617 | 0 |
| Fam214a | 0 | 0.59550791 | 0.374 | 0.163 | 0 |
| Filip1 | 0 | 0.89939564 | 0.725 | 0.388 | 0 |
| Tfdp2 | 0 | 1.67614157 | 0.449 | 0.105 | 0 |
| Atp2c1 | 0 | 0.58998767 | 0.309 | 0.114 | 0 |
| Acsl6 | 0 | 1.10005338 | 0.148 | 0.022 | 0 |
| Myh4 | 0 | -1.4515026 | 0.35 | 0.563 | 0 |
| Ube2g1 | 0 | 0.52279875 | 0.524 | 0.268 | 0 |
| Ssh2 | 0 | 1.13572277 | 0.479 | 0.164 | 0 |
| Myo18a | 0 | 0.72198278 | 0.386 | 0.16 | 0 |
| Bcas3 | 0 | 0.50654643 | 0.57 | 0.307 | 0 |
| Stat5b | 0 | 0.69087444 | 0.372 | 0.154 | 0 |
| Mylk4 | 0 | -1.3011075 | 0.362 | 0.637 | 0 |
| Ranbp9 | 0 | 0.77452528 | 0.258 | 0.083 | 0 |
| Adcy2 | 0 | 0.55854177 | 0.539 | 0.263 | 0 |
| Arrdc3 | 0 | 1.16767177 | 0.131 | 0.013 | 0 |
| Pik3r1 | 0 | 1.29893432 | 0.332 | 0.092 | 0 |
| Erbin | 0 | 1.08499337 | 0.247 | 0.058 | 0 |
| Pde4d | 0 | -0.6584264 | 0.834 | 0.87 | 0 |
| Lpin1 | 0 | 0.67033824 | 0.512 | 0.252 | 0 |
| Immp2l | 0 | 0.63033887 | 0.615 | 0.329 | 0 |
| Hectd1 | 0 | 0.71672586 | 0.277 | 0.098 | 0 |

| | | | | | |
|---|---|---|---|---|---|
| Retreg1 | 0 | 2.01376015 | 0.502 | 0.087 | 0 |
| Fbxo32 | 0 | 2.85620432 | 0.711 | 0.216 | 0 |
| Klhl38 | 0 | 1.06594205 | 0.209 | 0.041 | 0 |
| Rbfox1 | 0 | 0.4670636 | 0.902 | 0.772 | 0 |
| Tango2 | 0 | 1.78042351 | 0.327 | 0.05 | 0 |
| Trp63 | 0 | 0.58409072 | 0.567 | 0.314 | 0 |
| Zbtb20 | 0 | 0.51301281 | 0.934 | 0.749 | 0 |
| Zdhhc14 | 0 | 0.81101069 | 0.266 | 0.081 | 0 |
| Prkn | 0 | 0.54118776 | 0.852 | 0.6 | 0 |
| Srrm2 | 0 | 0.55844319 | 0.586 | 0.302 | 0 |
| Fkbp5 | 0 | 1.28971949 | 0.206 | 0.036 | 0 |
| Gm26917 | 0 | -1.2236169 | 0.923 | 0.961 | 0 |
| CT010467.1 | 0 | 1.57095201 | 0.7 | 0.143 | 0 |
| Gm42418 | 0 | -0.3239786 | 0.995 | 0.917 | 0 |
| Mocs1 | 0 | 1.01605558 | 0.16 | 0.028 | 0 |
| Plin4 | 0 | 1.04923423 | 0.21 | 0.046 | 0 |
| Rab12 | 0 | 0.89644791 | 0.208 | 0.054 | 0 |
| Gm36201 | 0 | 0.96266175 | 0.157 | 0.027 | 0 |
| Slc8a1 | 0 | 0.57591144 | 0.436 | 0.211 | 0 |
| Svil | 0 | 0.74268279 | 0.682 | 0.349 | 0 |
| Zeb1 | 0 | 0.75900152 | 0.379 | 0.141 | 0 |
| Npc1 | 0 | 1.20604454 | 0.21 | 0.039 | 0 |
| Arhgap26.1 | 0 | 1.97836268 | 0.296 | 0.036 | 0 |
| Sh3rf2 | 0 | 0.74572627 | 0.265 | 0.089 | 0 |
| Myot | 0 | 0.79138274 | 0.497 | 0.206 | 0 |
| Alpk2 | 0 | 0.6338387 | 0.393 | 0.164 | 0 |
| Dym | 0 | 0.63595545 | 0.376 | 0.149 | 0 |
| Ppp6r3 | 0 | 1.05146349 | 0.417 | 0.16 | 0 |
| Malat1 | 0 | -0.4718344 | 1 | 0.999 | 0 |
| Ahnak | 0 | 1.06104042 | 0.311 | 0.096 | 0 |
| Aldh1a1 | 0 | 0.9652897 | 0.303 | 0.096 | 0 |
| Zfand5 | 0 | 0.80843934 | 0.267 | 0.081 | 0 |
| Klf9 | 0 | 0.97440226 | 0.259 | 0.064 | 0 |
| Sorbs1 | 0 | 0.5520495 | 0.474 | 0.25 | 0 |
| Nrap | 0 | 0.58833539 | 0.714 | 0.416 | 0 |
| mt-Nd1 | 0 | -1.5490432 | 0.227 | 0.432 | 0 |
| mt-Nd2 | 0 | -1.4668382 | 0.313 | 0.48 | 0 |
| mt-Nd3 | 0 | -1.3384779 | 0.06 | 0.183 | 0 |
| mt-Nd4 | 0 | -1.3751344 | 0.212 | 0.387 | 0 |
| mt-Cytb | 0 | -1.487485 | 0.266 | 0.447 | 0 |
| 5730419F03I | 0 | 0.99234293 | 0.252 | 0.065 | 0 |
| Kbtbd12 | 3.86E-306 | 0.47341669 | 0.48 | 0.238 | 1.04E-301 |
| Garem1 | 1.24E-305 | 0.88631955 | 0.181 | 0.044 | 3.35E-301 |

| | | | | | |
|---|---|---|---|---|---|
| Ubn2 | 7.37E-305 | 0.4816656 | 0.435 | 0.209 | 1.99E-300 |
| Coq8a | 1.20E-304 | 0.73099806 | 0.236 | 0.077 | 3.23E-300 |
| Wnk2 | 8.25E-303 | 0.61790085 | 0.279 | 0.104 | 2.23E-298 |
| Asb2 | 1.02E-300 | 0.51718213 | 0.504 | 0.265 | 2.75E-296 |
| Pde4b | 1.59E-299 | 0.59890434 | 0.304 | 0.121 | 4.30E-295 |
| Rab2a | 3.16E-299 | 0.54099723 | 0.297 | 0.116 | 8.54E-295 |
| Slc12a2 | 9.44E-298 | 0.77817673 | 0.185 | 0.048 | 2.55E-293 |
| Pvt1 | 3.24E-290 | 0.68041999 | 0.267 | 0.101 | 8.77E-286 |
| Ncor1 | 9.84E-289 | 0.48424546 | 0.352 | 0.154 | 2.66E-284 |
| Gm26740 | 2.40E-287 | 0.4001923 | 0.859 | 0.672 | 6.50E-283 |
| Rb1 | 2.64E-285 | 0.76476056 | 0.191 | 0.054 | 7.14E-281 |
| Eda2r | 1.21E-284 | 0.8403606 | 0.181 | 0.049 | 3.28E-280 |
| Otud7b | 7.26E-282 | 0.5017194 | 0.274 | 0.104 | 1.96E-277 |
| Ip6k3 | 6.65E-280 | 0.81216328 | 0.168 | 0.041 | 1.80E-275 |
| Lin52 | 1.15E-279 | 0.52026116 | 0.3 | 0.122 | 3.11E-275 |
| Phka1 | 2.29E-278 | -1.0246824 | 0.372 | 0.486 | 6.20E-274 |
| Rps6ka3 | 3.21E-278 | 0.58672825 | 0.273 | 0.107 | 8.69E-274 |
| Son | 4.70E-278 | 0.44486584 | 0.34 | 0.146 | 1.27E-273 |
| Hivep2 | 3.49E-277 | 0.50929915 | 0.674 | 0.429 | 9.44E-273 |
| Vps13b | 7.24E-277 | 0.42604672 | 0.373 | 0.17 | 1.96E-272 |
| Mef2a | 1.74E-276 | 0.45304427 | 0.311 | 0.13 | 4.70E-272 |
| Igf1r | 1.90E-274 | 0.58360663 | 0.284 | 0.115 | 5.13E-270 |
| Arnt | 7.54E-274 | 0.66961392 | 0.188 | 0.054 | 2.04E-269 |
| Foxo3 | 7.01E-271 | 0.60591928 | 0.221 | 0.075 | 1.90E-266 |
| Lrmda | 8.28E-271 | 0.50747719 | 0.567 | 0.341 | 2.24E-266 |
| Amy1 | 1.68E-269 | 0.625414 | 0.171 | 0.044 | 4.53E-265 |
| Akap13 | 1.10E-266 | 0.37255116 | 0.505 | 0.266 | 2.96E-262 |
| Kif1b | 6.51E-266 | 0.40163933 | 0.65 | 0.392 | 1.76E-261 |
| Ubr2 | 4.94E-263 | 0.64051614 | 0.216 | 0.073 | 1.34E-258 |
| Smg1 | 5.04E-263 | 0.46838939 | 0.355 | 0.164 | 1.36E-258 |
| Dcaf6 | 3.53E-262 | 0.38528386 | 0.413 | 0.201 | 9.55E-258 |
| Tob2 | 1.18E-261 | 0.72886919 | 0.149 | 0.033 | 3.20E-257 |
| Ppp1r12a | 7.90E-261 | 0.43539721 | 0.328 | 0.146 | 2.14E-256 |
| Uri1 | 9.20E-261 | 0.57325298 | 0.227 | 0.08 | 2.49E-256 |
| Pcgf5 | 2.44E-260 | 0.44780173 | 0.269 | 0.105 | 6.59E-256 |
| Ptp4a3 | 2.72E-260 | 0.7788568 | 0.164 | 0.043 | 7.36E-256 |
| Tmcc1 | 6.24E-260 | 0.3941083 | 0.431 | 0.217 | 1.69E-255 |
| Enox2 | 1.27E-259 | 0.58874336 | 0.407 | 0.212 | 3.42E-255 |
| Birc6 | 1.30E-259 | 0.46519929 | 0.279 | 0.113 | 3.51E-255 |
| Bckdha | 2.03E-259 | 0.78914897 | 0.169 | 0.046 | 5.48E-255 |
| Ccdc192 | 8.41E-258 | 1.10114841 | 0.107 | 0.013 | 2.28E-253 |
| Cblb | 1.43E-256 | 0.44934359 | 0.302 | 0.129 | 3.87E-252 |
| Crebrf | 5.68E-256 | 0.6432069 | 0.204 | 0.067 | 1.54E-251 |

| | | | | | |
|---|---|---|---|---|---|
| Kcnq1ot1 | 1.55E-255 | 0.51810643 | 0.379 | 0.186 | 4.20E-251 |
| Neat1 | 1.19E-254 | 0.4645366 | 0.836 | 0.65 | 3.22E-250 |
| Klhl24 | 2.11E-254 | 0.58300494 | 0.198 | 0.063 | 5.72E-250 |
| Pnisr | 4.56E-254 | 0.33415977 | 0.303 | 0.126 | 1.23E-249 |
| Cdkal1 | 9.41E-254 | 0.4788527 | 0.516 | 0.286 | 2.54E-249 |
| Zfp638 | 6.01E-253 | 0.37172733 | 0.422 | 0.21 | 1.63E-248 |
| Phip | 1.82E-252 | 0.504077 | 0.232 | 0.085 | 4.93E-248 |
| 2610035D17I | 2.50E-252 | 0.85439903 | 0.139 | 0.03 | 6.76E-248 |
| Strn3 | 3.25E-252 | 0.38360976 | 0.407 | 0.201 | 8.79E-248 |
| Huwe1 | 5.38E-252 | 0.48293364 | 0.262 | 0.105 | 1.45E-247 |
| Nedd4l | 1.56E-251 | 0.52448031 | 0.232 | 0.085 | 4.21E-247 |
| B3galnt2 | 1.91E-249 | 0.52524275 | 0.249 | 0.097 | 5.16E-245 |
| Bin1 | 2.32E-249 | 0.39040081 | 0.472 | 0.254 | 6.27E-245 |
| Cdh13 | 3.39E-249 | -0.6398759 | 0.548 | 0.649 | 9.18E-245 |
| Cyp27a1 | 6.45E-249 | 0.78344851 | 0.184 | 0.058 | 1.74E-244 |
| Mga | 3.22E-247 | 0.64901989 | 0.197 | 0.065 | 8.72E-243 |
| Atf7ip | 5.10E-247 | 0.49156267 | 0.244 | 0.094 | 1.38E-242 |
| Itgb5 | 7.78E-247 | 0.75645963 | 0.156 | 0.041 | 2.10E-242 |
| Agpat3 | 1.00E-246 | 0.79217289 | 0.145 | 0.034 | 2.71E-242 |
| Gsk3b | 8.24E-246 | 0.47075596 | 0.328 | 0.151 | 2.23E-241 |
| Baz2b | 1.41E-245 | 0.40196586 | 0.345 | 0.161 | 3.82E-241 |
| Insr | 1.27E-244 | 0.40127462 | 0.372 | 0.179 | 3.45E-240 |
| Mllt10 | 1.61E-244 | 0.3975241 | 0.447 | 0.235 | 4.34E-240 |
| 3425401B19I | 4.12E-244 | 0.49826098 | 0.225 | 0.081 | 1.11E-239 |
| Gm19951 | 5.01E-244 | -1.1770422 | 0.561 | 0.627 | 1.35E-239 |
| Ralgapa2 | 7.30E-244 | 0.4791719 | 0.253 | 0.101 | 1.97E-239 |
| Ttll7 | 1.73E-243 | 0.45721962 | 0.293 | 0.126 | 4.68E-239 |
| Nmt1 | 7.91E-243 | 0.62920647 | 0.17 | 0.049 | 2.14E-238 |
| Atf6 | 5.87E-242 | 0.47391233 | 0.25 | 0.098 | 1.59E-237 |
| Zcchc7 | 5.43E-241 | 0.36070868 | 0.397 | 0.197 | 1.47E-236 |
| Ampd3 | 9.94E-241 | 0.79540595 | 0.14 | 0.032 | 2.69E-236 |
| 4632427E13I | 1.70E-240 | 0.69710894 | 0.142 | 0.034 | 4.60E-236 |
| B3galt1 | 4.23E-240 | 0.58930635 | 0.22 | 0.081 | 1.14E-235 |
| Mia2 | 2.09E-239 | 0.47596463 | 0.25 | 0.101 | 5.64E-235 |
| Ppp3ca | 3.30E-239 | 0.35462584 | 0.657 | 0.409 | 8.92E-235 |
| Rps6ka2 | 5.02E-239 | 0.3827476 | 0.345 | 0.161 | 1.36E-234 |
| Wdr33 | 7.07E-239 | 0.42009622 | 0.33 | 0.153 | 1.91E-234 |
| Lrrc58 | 1.17E-237 | 0.86053696 | 0.12 | 0.023 | 3.17E-233 |
| Gm20388 | 1.32E-237 | 0.50469965 | 0.288 | 0.128 | 3.56E-233 |
| Smyd2 | 8.26E-237 | 0.63137227 | 0.177 | 0.055 | 2.23E-232 |
| Tbc1d5 | 1.75E-236 | 0.38408569 | 0.309 | 0.138 | 4.72E-232 |
| Akt2 | 2.83E-236 | 0.76745858 | 0.123 | 0.024 | 7.65E-232 |
| Srek1 | 1.59E-235 | 0.37251582 | 0.288 | 0.124 | 4.31E-231 |

| | | | | | |
|---|---|---|---|---|---|
| Vegfd | 3.54E-235 | 0.89591527 | 0.104 | 0.014 | 9.56E-231 |
| Ash1l | 1.57E-234 | 0.32769331 | 0.431 | 0.221 | 4.26E-230 |
| Usp2 | 8.54E-234 | 0.46773652 | 0.232 | 0.089 | 2.31E-229 |
| Tenm3 | 8.68E-234 | 0.50907902 | 0.261 | 0.109 | 2.35E-229 |
| Rapgef2 | 1.15E-233 | 0.50043006 | 0.235 | 0.092 | 3.11E-229 |
| Rb1cc1 | 9.44E-233 | 0.37517445 | 0.271 | 0.114 | 2.55E-228 |
| Tcf12 | 1.61E-232 | 0.36322285 | 0.338 | 0.16 | 4.35E-228 |
| Klhl7 | 3.68E-231 | 0.58943252 | 0.163 | 0.047 | 9.95E-227 |
| Camta1 | 3.57E-230 | 0.47048257 | 0.239 | 0.095 | 9.64E-226 |
| Xirp2 | 2.85E-228 | 0.33556824 | 0.547 | 0.323 | 7.71E-224 |
| Ank | 5.17E-227 | 0.36617492 | 0.345 | 0.166 | 1.40E-222 |
| Scaf11 | 1.03E-226 | 0.48535728 | 0.192 | 0.065 | 2.77E-222 |
| Jph1 | 2.43E-226 | -0.726612 | 0.578 | 0.629 | 6.56E-222 |
| Lrrc2 | 4.83E-226 | 0.47954611 | 0.281 | 0.124 | 1.30E-221 |
| Dysf | 5.37E-226 | 0.34918241 | 0.323 | 0.149 | 1.45E-221 |
| Brd4 | 6.81E-226 | 0.45496694 | 0.264 | 0.113 | 1.84E-221 |
| Ipo13 | 1.75E-225 | 0.69788801 | 0.141 | 0.036 | 4.74E-221 |
| Zfp280d | 1.55E-224 | 0.42258365 | 0.262 | 0.111 | 4.18E-220 |
| Lmod2 | 1.20E-223 | 0.62283335 | 0.204 | 0.074 | 3.25E-219 |
| Nsmce2 | 1.38E-223 | 0.41129449 | 0.304 | 0.139 | 3.72E-219 |
| Kat2b | 2.93E-223 | 0.65315049 | 0.161 | 0.049 | 7.92E-219 |
| Ppp2r3a | 9.09E-223 | 0.3261725 | 0.399 | 0.204 | 2.46E-218 |
| Abcc9 | 1.18E-222 | 0.28583108 | 0.339 | 0.16 | 3.19E-218 |
| Anxa11 | 2.94E-222 | 0.5742221 | 0.161 | 0.048 | 7.94E-218 |
| Rbm6 | 1.97E-220 | 0.38778836 | 0.312 | 0.147 | 5.32E-216 |
| Hnrnpu | 2.20E-219 | 0.34891906 | 0.273 | 0.118 | 5.96E-215 |
| Scaf8 | 6.69E-219 | 0.41704986 | 0.222 | 0.085 | 1.81E-214 |
| Ccnl2 | 8.05E-219 | 0.52815155 | 0.203 | 0.075 | 2.18E-214 |
| Chd2 | 9.21E-219 | 0.36623501 | 0.334 | 0.16 | 2.49E-214 |
| Bnip3 | 3.51E-218 | 0.66575494 | 0.128 | 0.029 | 9.50E-214 |
| Sfpq | 1.94E-217 | 0.48601002 | 0.188 | 0.065 | 5.25E-213 |
| Arhgap35 | 2.41E-217 | 0.48600921 | 0.214 | 0.082 | 6.51E-213 |
| Dyrk1a | 3.66E-217 | 0.28767808 | 0.368 | 0.183 | 9.91E-213 |
| Cacng1 | 1.75E-216 | 0.48906993 | 0.208 | 0.079 | 4.73E-212 |
| Tgfbr1 | 3.46E-216 | 0.5056288 | 0.195 | 0.07 | 9.35E-212 |
| Gigyf2 | 5.48E-216 | 0.37365379 | 0.257 | 0.109 | 1.48E-211 |
| Eif4g3 | 5.55E-216 | 0.35649613 | 0.687 | 0.448 | 1.50E-211 |
| Psme4 | 6.39E-216 | 0.29159358 | 0.37 | 0.182 | 1.73E-211 |
| Rsrp1 | 6.54E-215 | 0.33760005 | 0.278 | 0.122 | 1.77E-210 |
| Mkln1 | 1.60E-214 | 0.4128219 | 0.272 | 0.121 | 4.31E-210 |
| Csnk1a1 | 2.96E-214 | 0.45519059 | 0.191 | 0.068 | 8.01E-210 |
| B4galt1 | 6.54E-214 | 0.69279916 | 0.136 | 0.035 | 1.77E-209 |
| Ucp3 | 7.60E-212 | 0.72567345 | 0.117 | 0.025 | 2.05E-207 |

| | | | | | |
|---|---|---|---|---|---|
| Pde7a | 1.28E-211 | 0.4091593 | 0.522 | 0.309 | 3.46E-207 |
| Fggy | 1.96E-211 | 0.46307307 | 0.21 | 0.081 | 5.30E-207 |
| Kmt2e | 2.33E-211 | 0.3637745 | 0.244 | 0.102 | 6.29E-207 |
| Sgms1 | 3.21E-211 | 0.32897641 | 0.479 | 0.268 | 8.68E-207 |
| Tnrc6c | 1.15E-210 | 0.27297857 | 0.41 | 0.214 | 3.11E-206 |
| Tmem38b | 3.40E-210 | 0.69561828 | 0.12 | 0.027 | 9.18E-206 |
| Ankhd1 | 1.07E-209 | 0.43376097 | 0.207 | 0.079 | 2.90E-205 |
| Tcap | 1.74E-209 | 0.5721796 | 0.271 | 0.132 | 4.71E-205 |
| Ubr4 | 2.47E-209 | 0.52746346 | 0.19 | 0.069 | 6.67E-205 |
| Trim54 | 3.84E-209 | 0.34160589 | 0.269 | 0.118 | 1.04E-204 |
| Coro6 | 6.21E-208 | 0.58176143 | 0.144 | 0.041 | 1.68E-203 |
| Pcnt | 8.08E-208 | 0.3883151 | 0.282 | 0.127 | 2.18E-203 |
| Rapgef1 | 1.73E-207 | 0.39352973 | 0.301 | 0.143 | 4.68E-203 |
| Etfa | 8.21E-207 | 0.46214509 | 0.202 | 0.076 | 2.22E-202 |
| Tulp4 | 5.03E-206 | 0.47906261 | 0.192 | 0.071 | 1.36E-201 |
| Crebbp | 1.00E-205 | 0.29610722 | 0.332 | 0.161 | 2.71E-201 |
| Tmem233 | 1.01E-205 | -1.365818 | 0.158 | 0.263 | 2.73E-201 |
| Hibadh | 1.66E-205 | 0.37036854 | 0.216 | 0.085 | 4.49E-201 |
| Larp4b | 1.74E-205 | 0.39865587 | 0.236 | 0.099 | 4.70E-201 |
| A330023F24 | 3.98E-205 | 0.31955966 | 0.301 | 0.141 | 1.08E-200 |
| Kmt2a | 5.04E-205 | 0.35211799 | 0.269 | 0.12 | 1.36E-200 |
| Faf1 | 5.39E-205 | 0.32354972 | 0.335 | 0.164 | 1.46E-200 |
| Fam13c | 4.36E-204 | 0.76964767 | 0.101 | 0.018 | 1.18E-199 |
| Sgcg | 4.43E-204 | 0.43317928 | 0.215 | 0.085 | 1.20E-199 |
| Ccny | 4.96E-204 | 0.45668965 | 0.185 | 0.067 | 1.34E-199 |
| Atp9b | 4.31E-203 | 0.34577525 | 0.237 | 0.1 | 1.16E-198 |
| Ankrd10 | 2.90E-202 | 0.69979253 | 0.141 | 0.041 | 7.83E-198 |
| N4bp2l2 | 4.21E-202 | 0.36555241 | 0.259 | 0.115 | 1.14E-197 |
| Apex2 | 4.66E-202 | 0.35967323 | 0.303 | 0.144 | 1.26E-197 |
| Akap8l | 1.51E-201 | 0.4233394 | 0.237 | 0.102 | 4.08E-197 |
| Usp47 | 2.88E-201 | 0.31966887 | 0.252 | 0.109 | 7.79E-197 |
| Airn | 9.96E-201 | 0.35023387 | 0.606 | 0.383 | 2.69E-196 |
| Raf1 | 2.13E-200 | 0.37393302 | 0.239 | 0.102 | 5.76E-196 |
| Zfand3 | 2.70E-200 | 0.28992695 | 0.479 | 0.27 | 7.29E-196 |
| Tnrc6b | 2.34E-199 | 0.31287592 | 0.641 | 0.396 | 6.32E-195 |
| Arhgef12 | 2.98E-199 | 0.33745924 | 0.297 | 0.142 | 8.06E-195 |
| Nfix | 5.90E-199 | 0.32825546 | 0.258 | 0.114 | 1.60E-194 |
| Ppfibp2 | 6.52E-199 | 0.4545526 | 0.2 | 0.077 | 1.76E-194 |
| Hook3 | 4.90E-198 | 0.44163029 | 0.174 | 0.061 | 1.33E-193 |
| Pcgf3 | 7.04E-198 | 0.64999124 | 0.13 | 0.035 | 1.90E-193 |
| Rsrc1 | 1.64E-197 | 0.52374254 | 0.159 | 0.052 | 4.44E-193 |
| Asb5 | 1.92E-196 | 0.36437469 | 0.254 | 0.112 | 5.19E-192 |
| Acss2 | 3.39E-196 | 0.50942419 | 0.195 | 0.075 | 9.16E-192 |

| | | | | | |
|---|---|---|---|---|---|
| Picalm | 3.62E-196 | 0.31235451 | 0.256 | 0.114 | 9.79E-192 |
| Diaph2 | 1.10E-195 | 0.43546227 | 0.235 | 0.101 | 2.98E-191 |
| Dst | 3.62E-195 | 0.28393554 | 0.8 | 0.563 | 9.79E-191 |
| Setd5 | 4.89E-194 | 0.42597883 | 0.222 | 0.094 | 1.32E-189 |
| Zfp445 | 1.19E-193 | 0.4869421 | 0.171 | 0.06 | 3.21E-189 |
| Dnajc1 | 2.39E-193 | 0.36284496 | 0.27 | 0.125 | 6.47E-189 |
| Tgfbr2 | 2.97E-193 | 0.56101978 | 0.146 | 0.045 | 8.03E-189 |
| Gm49906 | 3.88E-193 | 0.27327393 | 0.467 | 0.261 | 1.05E-188 |
| 6230400D17 | 7.54E-193 | 0.60084175 | 0.12 | 0.03 | 2.04E-188 |
| Snx13 | 1.10E-192 | 0.37569599 | 0.2 | 0.078 | 2.97E-188 |
| Pibf1 | 1.46E-192 | 0.32542189 | 0.267 | 0.122 | 3.96E-188 |
| Asb11 | 1.53E-192 | 0.52450837 | 0.169 | 0.059 | 4.13E-188 |
| Kdm6a | 2.39E-192 | 0.31443698 | 0.229 | 0.096 | 6.47E-188 |
| Pbxip1 | 5.60E-192 | 0.45014838 | 0.223 | 0.095 | 1.52E-187 |
| Ube4b | 1.47E-191 | 0.46403269 | 0.195 | 0.076 | 3.98E-187 |
| R3hdm1 | 2.22E-191 | 0.36126036 | 0.216 | 0.088 | 5.99E-187 |
| Des | 2.96E-191 | 0.49825726 | 0.223 | 0.097 | 8.00E-187 |
| Slc20a2 | 4.77E-191 | 0.40748024 | 0.202 | 0.08 | 1.29E-186 |
| Dnm3 | 9.33E-191 | 0.67938885 | 0.126 | 0.034 | 2.52E-186 |
| Ryr1 | 1.65E-190 | -0.5433303 | 0.801 | 0.782 | 4.46E-186 |
| Psma3 | 1.88E-190 | 0.50248253 | 0.169 | 0.06 | 5.07E-186 |
| Phc3 | 1.99E-190 | 0.47211641 | 0.166 | 0.058 | 5.39E-186 |
| Rev3l | 2.17E-190 | 0.28434168 | 0.385 | 0.203 | 5.86E-186 |
| Taf1d | 3.49E-190 | 0.43051383 | 0.175 | 0.063 | 9.45E-186 |
| Slmap | 4.42E-190 | 0.28276065 | 0.291 | 0.139 | 1.20E-185 |
| Tet2 | 7.33E-190 | 0.41227307 | 0.188 | 0.072 | 1.98E-185 |
| Ube3a | 6.78E-189 | 0.35524058 | 0.235 | 0.102 | 1.83E-184 |
| Jpx | 2.99E-188 | 0.37399486 | 0.254 | 0.116 | 8.08E-184 |
| Mpp6 | 5.74E-188 | 0.41496906 | 0.172 | 0.062 | 1.55E-183 |
| Pcnx | 6.40E-188 | 0.47166809 | 0.154 | 0.051 | 1.73E-183 |
| Mef2d | 8.27E-188 | 0.29497699 | 0.264 | 0.121 | 2.24E-183 |
| Nr2c2 | 2.40E-187 | 0.4959826 | 0.139 | 0.042 | 6.50E-183 |
| Tbc1d4 | 9.39E-187 | 0.29378612 | 0.367 | 0.193 | 2.54E-182 |
| Ppp1cb | 1.54E-186 | 0.57230936 | 0.128 | 0.036 | 4.16E-182 |
| Gm49417 | 2.73E-186 | 0.60347279 | 0.192 | 0.077 | 7.38E-182 |
| Ube2h | 6.85E-186 | 0.2789693 | 0.416 | 0.229 | 1.85E-181 |
| Tbl1xr1 | 9.36E-186 | 0.30743828 | 0.245 | 0.109 | 2.53E-181 |
| Mon2 | 1.01E-185 | 0.37514729 | 0.2 | 0.08 | 2.74E-181 |
| Acsl1 | 1.52E-185 | 0.3895177 | 0.236 | 0.104 | 4.10E-181 |
| Nt5dc1 | 3.01E-185 | 0.43859992 | 0.15 | 0.049 | 8.15E-181 |
| Lamp2 | 6.42E-185 | 0.33350151 | 0.252 | 0.115 | 1.74E-180 |
| Usp25 | 9.69E-185 | 0.38412443 | 0.179 | 0.067 | 2.62E-180 |
| Rbm39 | 1.35E-184 | 0.34351004 | 0.528 | 0.326 | 3.64E-180 |

| | | | | | |
|---|---|---|---|---|---|
| Rnf216 | 2.28E-184 | 0.46156973 | 0.193 | 0.077 | 6.15E-180 |
| Mecp2 | 4.55E-184 | 0.36674318 | 0.215 | 0.091 | 1.23E-179 |
| Mtdh | 2.81E-183 | 0.26953308 | 0.284 | 0.137 | 7.59E-179 |
| Aox1 | 7.67E-183 | 0.50571164 | 0.193 | 0.077 | 2.07E-178 |
| Mllt3 | 1.33E-182 | 0.37100639 | 0.228 | 0.1 | 3.60E-178 |
| Gabrb1 | 2.35E-182 | 0.28830045 | 0.173 | 0.064 | 6.37E-178 |
| Dapk2 | 3.12E-182 | 0.52236441 | 0.198 | 0.081 | 8.43E-178 |
| Gtdc1 | 3.16E-182 | 0.4153256 | 0.198 | 0.08 | 8.54E-178 |
| Rbl2 | 4.26E-182 | 0.62887547 | 0.113 | 0.028 | 1.15E-177 |
| Mlxip | 7.12E-182 | 0.40546753 | 0.232 | 0.103 | 1.93E-177 |
| Nfe2l2 | 3.91E-180 | 0.48409092 | 0.139 | 0.043 | 1.06E-175 |
| Tmem135 | 4.26E-180 | 0.35734302 | 0.218 | 0.094 | 1.15E-175 |
| Rilpl1 | 7.18E-180 | 0.29968378 | 0.346 | 0.182 | 1.94E-175 |
| Gm47071 | 1.63E-179 | 0.65668524 | 0.111 | 0.028 | 4.41E-175 |
| Rmnd5a | 2.23E-179 | 0.47098165 | 0.143 | 0.046 | 6.03E-175 |
| Anapc5 | 6.23E-179 | 0.45949932 | 0.155 | 0.053 | 1.68E-174 |
| Psmd11 | 3.33E-178 | 0.41073718 | 0.18 | 0.069 | 9.02E-174 |
| Vwa8 | 5.42E-178 | 0.2717865 | 0.362 | 0.191 | 1.46E-173 |
| Camk2a | 8.17E-178 | -0.9461891 | 0.486 | 0.504 | 2.21E-173 |
| Txnl1 | 1.54E-177 | 0.38808259 | 0.179 | 0.068 | 4.16E-173 |
| Lrrfip2 | 1.39E-176 | 0.25635114 | 0.254 | 0.117 | 3.77E-172 |
| Nckap1 | 1.38E-175 | 0.43092377 | 0.155 | 0.055 | 3.72E-171 |
| Nek7 | 1.69E-175 | 0.36049082 | 0.182 | 0.071 | 4.56E-171 |
| Nsd3 | 1.98E-175 | 0.29788526 | 0.238 | 0.108 | 5.35E-171 |
| 4930402D18 | 7.29E-175 | 0.40995539 | 0.15 | 0.051 | 1.97E-170 |
| Zfp407 | 7.74E-175 | 0.26735829 | 0.261 | 0.122 | 2.09E-170 |
| Wbp1l | 1.72E-174 | 0.40213539 | 0.191 | 0.077 | 4.65E-170 |
| Cecr2 | 6.63E-174 | 0.34723202 | 0.199 | 0.082 | 1.79E-169 |
| Ide | 7.85E-174 | 0.41781671 | 0.167 | 0.062 | 2.12E-169 |
| Sqstm1 | 5.40E-173 | 0.57415934 | 0.135 | 0.043 | 1.46E-168 |
| Cul3 | 2.57E-172 | 0.39227505 | 0.159 | 0.058 | 6.95E-168 |
| Fam219a | 3.36E-172 | 0.48082161 | 0.165 | 0.062 | 9.08E-168 |
| Rbm33 | 5.40E-172 | 0.29382594 | 0.214 | 0.093 | 1.46E-167 |
| Mid1 | 1.18E-170 | 0.49792451 | 0.141 | 0.048 | 3.18E-166 |
| Col4a3bp | 1.22E-170 | 0.38336791 | 0.151 | 0.053 | 3.30E-166 |
| Atp1a2 | 1.88E-170 | -0.7311659 | 0.673 | 0.68 | 5.08E-166 |
| Ppp2ca | 7.71E-170 | 0.44665679 | 0.182 | 0.074 | 2.08E-165 |
| Kif5b | 2.95E-169 | 0.43290492 | 0.164 | 0.062 | 7.99E-165 |
| Fbxl20 | 5.81E-169 | 0.31177551 | 0.201 | 0.085 | 1.57E-164 |
| Psmd1 | 1.97E-168 | 0.32900073 | 0.2 | 0.084 | 5.34E-164 |
| Rasa3 | 2.32E-168 | 0.5873723 | 0.103 | 0.025 | 6.28E-164 |
| 4932438A13 | 2.83E-168 | 0.33679489 | 0.168 | 0.064 | 7.67E-164 |
| Asxl1 | 9.90E-168 | 0.36992742 | 0.173 | 0.068 | 2.68E-163 |

| | | | | | |
|---|---|---|---|---|---|
| Vps37a | 1.99E-167 | 0.44891525 | 0.129 | 0.04 | 5.38E-163 |
| Mrpl1 | 2.91E-167 | 0.5024556 | 0.108 | 0.028 | 7.87E-163 |
| Smg6 | 1.17E-165 | 0.26986795 | 0.275 | 0.137 | 3.15E-161 |
| Uvrag | 2.11E-165 | 0.40652351 | 0.152 | 0.055 | 5.70E-161 |
| Rab7 | 4.29E-165 | 0.27824662 | 0.315 | 0.165 | 1.16E-160 |
| Rnf13 | 5.61E-165 | 0.44317416 | 0.13 | 0.042 | 1.52E-160 |
| Ubc | 1.14E-164 | 0.42248314 | 0.134 | 0.044 | 3.08E-160 |
| Med13 | 1.32E-164 | 0.27609451 | 0.223 | 0.101 | 3.56E-160 |
| Clk1 | 2.62E-164 | 0.38649261 | 0.175 | 0.07 | 7.08E-160 |
| Jun | 3.29E-164 | 0.60783701 | 0.106 | 0.028 | 8.90E-160 |
| Myom2 | 3.51E-164 | 0.28385883 | 0.445 | 0.259 | 9.49E-160 |
| Susd6 | 3.89E-164 | 0.26705306 | 0.236 | 0.11 | 1.05E-159 |
| Flnc | 5.22E-164 | 0.43936673 | 0.256 | 0.127 | 1.41E-159 |
| Il6st | 2.94E-163 | 0.49661112 | 0.11 | 0.03 | 7.94E-159 |
| Cd2ap | 9.66E-163 | 0.33657477 | 0.191 | 0.081 | 2.61E-158 |
| Arid2 | 2.70E-162 | 0.28415338 | 0.17 | 0.066 | 7.31E-158 |
| Eef2k | 3.53E-162 | 0.33828028 | 0.286 | 0.146 | 9.55E-158 |
| Tax1bp1 | 5.60E-162 | 0.39853572 | 0.128 | 0.041 | 1.52E-157 |
| Acaca | 5.81E-162 | 0.39499204 | 0.195 | 0.084 | 1.57E-157 |
| Ggnbp2 | 1.13E-161 | 0.30142988 | 0.196 | 0.084 | 3.06E-157 |
| Pfkfb3 | 2.25E-161 | 0.27555808 | 0.383 | 0.222 | 6.08E-157 |
| Kpna3 | 5.01E-161 | 0.31892015 | 0.174 | 0.069 | 1.35E-156 |
| Epc2 | 6.19E-161 | 0.38212862 | 0.145 | 0.052 | 1.67E-156 |
| Tlk2 | 1.38E-160 | 0.34706151 | 0.187 | 0.079 | 3.72E-156 |
| Bmpr1b | 1.54E-160 | 0.35700515 | 0.242 | 0.116 | 4.17E-156 |
| Itsn2 | 1.66E-160 | 0.32314677 | 0.189 | 0.08 | 4.50E-156 |
| Rictor | 1.44E-159 | 0.38317803 | 0.151 | 0.056 | 3.90E-155 |
| Baz1b | 1.54E-159 | 0.26179352 | 0.228 | 0.105 | 4.17E-155 |
| Ranbp2 | 4.15E-159 | 0.37838285 | 0.169 | 0.068 | 1.12E-154 |
| Vldlr | 5.67E-159 | 0.40524307 | 0.165 | 0.065 | 1.53E-154 |
| Smyd3 | 5.82E-159 | 0.28948644 | 0.282 | 0.144 | 1.57E-154 |
| Rprd1b | 9.94E-159 | 0.43554772 | 0.125 | 0.04 | 2.69E-154 |
| Elp4 | 1.05E-157 | 0.2596425 | 0.215 | 0.097 | 2.85E-153 |
| Mreg | 2.11E-157 | 0.43014917 | 0.134 | 0.045 | 5.72E-153 |
| Trdn | 3.52E-157 | -0.2873938 | 0.88 | 0.9 | 9.52E-153 |
| Babam2 | 6.07E-157 | 0.25852563 | 0.301 | 0.157 | 1.64E-152 |
| Acss1 | 7.55E-157 | 0.55861205 | 0.123 | 0.039 | 2.04E-152 |
| Gab1 | 3.51E-156 | 0.27418166 | 0.204 | 0.09 | 9.50E-152 |
| Fam120c | 5.80E-156 | 0.38576191 | 0.15 | 0.056 | 1.57E-151 |
| Ubr1 | 1.00E-155 | 0.3801694 | 0.159 | 0.062 | 2.72E-151 |
| Phf3 | 1.49E-155 | 0.40992636 | 0.142 | 0.051 | 4.03E-151 |
| Ncor2 | 1.87E-155 | 0.4064439 | 0.125 | 0.041 | 5.06E-151 |
| Sec62 | 4.63E-155 | 0.49114674 | 0.104 | 0.028 | 1.25E-150 |

| | | | | | |
|---|---|---|---|---|---|
| Fbxw7 | 1.39E-154 | 0.31666914 | 0.162 | 0.064 | 3.76E-150 |
| Synj2 | 2.02E-154 | 0.49643215 | 0.114 | 0.034 | 5.47E-150 |
| Dtnbp1 | 2.22E-154 | 0.54479613 | 0.107 | 0.03 | 5.99E-150 |
| AW554918 | 9.15E-154 | 0.31221262 | 0.15 | 0.056 | 2.47E-149 |
| Maco1 | 9.51E-154 | 0.34911336 | 0.148 | 0.055 | 2.57E-149 |
| Mettl7a1 | 1.01E-153 | 0.41852446 | 0.148 | 0.055 | 2.74E-149 |
| Lnpep | 1.13E-153 | 0.27870989 | 0.208 | 0.094 | 3.06E-149 |
| Stag1 | 2.28E-153 | 0.25943162 | 0.221 | 0.103 | 6.16E-149 |
| Bptf | 7.80E-153 | 0.29663512 | 0.204 | 0.092 | 2.11E-148 |
| Tdrd3 | 8.52E-153 | 0.2970884 | 0.17 | 0.069 | 2.30E-148 |
| Acox1 | 5.89E-152 | 0.54016732 | 0.114 | 0.035 | 1.59E-147 |
| Pfkfb1 | 1.28E-151 | 0.30875778 | 0.219 | 0.102 | 3.46E-147 |
| Ctnna1 | 3.16E-151 | 0.38014457 | 0.155 | 0.061 | 8.55E-147 |
| Synrg | 4.96E-151 | 0.37789718 | 0.135 | 0.047 | 1.34E-146 |
| Aebp2 | 6.66E-151 | 0.46221223 | 0.128 | 0.044 | 1.80E-146 |
| Glyr1 | 2.64E-150 | 0.4280637 | 0.152 | 0.059 | 7.13E-146 |
| Rnf169 | 2.76E-150 | 0.44785211 | 0.109 | 0.032 | 7.46E-146 |
| Acvr1b | 5.25E-150 | 0.43494346 | 0.106 | 0.03 | 1.42E-145 |
| Rad23b | 9.73E-150 | 0.35248908 | 0.166 | 0.068 | 2.63E-145 |
| Bclaf1 | 1.07E-149 | 0.35465988 | 0.158 | 0.063 | 2.90E-145 |
| Slc7a6 | 2.42E-149 | 0.4355549 | 0.107 | 0.031 | 6.55E-145 |
| Ttll11 | 3.06E-149 | 0.30809957 | 0.224 | 0.107 | 8.26E-145 |
| Dcaf8 | 4.72E-149 | 0.27016703 | 0.26 | 0.131 | 1.28E-144 |
| 11-Sep | 3.23E-148 | 0.27493298 | 0.196 | 0.088 | 8.73E-144 |
| Hivep3 | 8.65E-148 | 0.56385461 | 0.129 | 0.045 | 2.34E-143 |
| Zfp292 | 2.18E-147 | 0.30468638 | 0.158 | 0.063 | 5.89E-143 |
| Scfd2 | 2.30E-147 | 0.30945712 | 0.143 | 0.053 | 6.21E-143 |
| Macrod2 | 4.85E-147 | 0.38206893 | 0.613 | 0.436 | 1.31E-142 |
| Acvr2a | 5.83E-147 | 0.30566747 | 0.139 | 0.051 | 1.58E-142 |
| Dusp13 | 1.11E-146 | 0.34185847 | 0.176 | 0.075 | 3.00E-142 |
| Dpyd | 1.49E-146 | 0.49065967 | 0.173 | 0.075 | 4.03E-142 |
| Fam120a | 4.49E-146 | 0.37653979 | 0.142 | 0.054 | 1.21E-141 |
| Aatf | 6.92E-146 | 0.4102924 | 0.12 | 0.04 | 1.87E-141 |
| Rheb | 2.87E-145 | 0.37073717 | 0.137 | 0.05 | 7.77E-141 |
| Cpeb1 | 6.83E-145 | 0.32672344 | 0.174 | 0.075 | 1.85E-140 |
| Herc4 | 1.23E-144 | 0.33310993 | 0.167 | 0.07 | 3.34E-140 |
| Ifrd1 | 1.32E-144 | 0.29156278 | 0.189 | 0.085 | 3.57E-140 |
| Cyld | 3.51E-144 | 0.40827989 | 0.102 | 0.029 | 9.50E-140 |
| Oga | 4.48E-144 | 0.4273171 | 0.126 | 0.044 | 1.21E-139 |
| Pten | 5.86E-144 | 0.32401814 | 0.157 | 0.063 | 1.58E-139 |
| Nudcd3 | 1.16E-143 | 0.33328612 | 0.155 | 0.062 | 3.13E-139 |
| Sel1l3 | 1.45E-143 | 0.36716284 | 0.131 | 0.047 | 3.93E-139 |
| Eprs | 1.50E-143 | 0.29935632 | 0.171 | 0.072 | 4.06E-139 |

| | | | | | |
|---|---|---|---|---|---|
| Me1 | 1.95E-143 | 0.41421597 | 0.119 | 0.04 | 5.28E-139 |
| Abi1 | 2.63E-143 | 0.35251474 | 0.145 | 0.056 | 7.11E-139 |
| Epg5 | 2.69E-143 | 0.34457996 | 0.132 | 0.047 | 7.28E-139 |
| Polr1a | 2.88E-143 | 0.49285005 | 0.102 | 0.029 | 7.80E-139 |
| Tnks | 1.06E-142 | 0.31836487 | 0.142 | 0.054 | 2.86E-138 |
| Abcc1 | 1.10E-142 | 0.2571596 | 0.165 | 0.069 | 2.96E-138 |
| Fus | 1.25E-142 | 0.42364056 | 0.129 | 0.047 | 3.38E-138 |
| Stt3b | 1.29E-142 | 0.457177 | 0.106 | 0.032 | 3.49E-138 |
| Kpna4 | 2.42E-142 | 0.30551317 | 0.161 | 0.066 | 6.53E-138 |
| Pou2f1 | 4.10E-142 | 0.2752399 | 0.17 | 0.072 | 1.11E-137 |
| Ddx5 | 4.86E-142 | 0.29276891 | 0.225 | 0.112 | 1.31E-137 |
| 7-Mar | 8.06E-141 | 0.3396876 | 0.142 | 0.055 | 2.18E-136 |
| Scarb2 | 8.80E-141 | 0.327227 | 0.144 | 0.056 | 2.38E-136 |
| Tmem87a | 1.78E-140 | 0.2721462 | 0.186 | 0.083 | 4.81E-136 |
| Wdr26 | 1.90E-140 | 0.31299256 | 0.17 | 0.074 | 5.13E-136 |
| Fgf13 | 6.91E-140 | 0.3538137 | 0.819 | 0.679 | 1.87E-135 |
| Agap1 | 1.33E-139 | 0.26835499 | 0.184 | 0.082 | 3.58E-135 |
| Sorbs3 | 1.82E-139 | 0.46215845 | 0.102 | 0.03 | 4.93E-135 |
| Clk4 | 2.13E-139 | 0.33900818 | 0.13 | 0.047 | 5.76E-135 |
| Zfp148 | 5.72E-139 | 0.26473048 | 0.208 | 0.099 | 1.55E-134 |
| Ece1 | 2.50E-138 | 0.35351998 | 0.188 | 0.086 | 6.76E-134 |
| Nploc4 | 7.70E-138 | 0.36630632 | 0.153 | 0.063 | 2.08E-133 |
| Slf1 | 7.92E-138 | 0.37311673 | 0.119 | 0.04 | 2.14E-133 |
| Sppl3 | 1.55E-137 | 0.28469654 | 0.17 | 0.073 | 4.18E-133 |
| BC005537 | 2.31E-137 | 0.46091551 | 0.101 | 0.03 | 6.26E-133 |
| Rabgap1 | 8.35E-137 | 0.26754283 | 0.167 | 0.071 | 2.26E-132 |
| Ypel2 | 2.22E-136 | 0.44384156 | 0.118 | 0.041 | 6.00E-132 |
| Top1 | 2.24E-136 | 0.39139041 | 0.12 | 0.042 | 6.07E-132 |
| Ube2d3 | 2.84E-136 | 0.28599863 | 0.162 | 0.069 | 7.69E-132 |
| Slc49a4 | 4.51E-136 | 0.3061948 | 0.135 | 0.051 | 1.22E-131 |
| Mtss1 | 4.98E-136 | 0.28512692 | 0.162 | 0.068 | 1.35E-131 |
| Arid1a | 5.77E-136 | 0.29355893 | 0.148 | 0.06 | 1.56E-131 |
| Fubp1 | 6.17E-136 | 0.37726129 | 0.113 | 0.038 | 1.67E-131 |
| Taf15 | 7.96E-136 | 0.33747461 | 0.153 | 0.063 | 2.15E-131 |
| Yy1 | 1.16E-135 | 0.33019277 | 0.128 | 0.047 | 3.15E-131 |
| Itch | 2.20E-135 | 0.25029158 | 0.191 | 0.088 | 5.96E-131 |
| Rffl | 2.44E-135 | 0.37202923 | 0.143 | 0.057 | 6.59E-131 |
| Ldlrad3 | 7.14E-135 | 0.50303207 | 0.147 | 0.061 | 1.93E-130 |
| Dennd4c | 4.21E-134 | 0.27127034 | 0.152 | 0.063 | 1.14E-129 |
| Map2k5 | 4.31E-134 | 0.25305372 | 0.171 | 0.075 | 1.17E-129 |
| Clec2d | 4.61E-134 | 0.32020005 | 0.228 | 0.119 | 1.25E-129 |
| Gpt2 | 5.54E-134 | 0.33141459 | 0.144 | 0.057 | 1.50E-129 |
| Ubap2l | 8.83E-134 | 0.32291232 | 0.151 | 0.062 | 2.39E-129 |

| | | | | | |
|---|---|---|---|---|---|
| Ccar1 | 4.07E-133 | 0.28015995 | 0.142 | 0.056 | 1.10E-128 |
| Zfyve1 | 7.51E-133 | 0.4055715 | 0.12 | 0.042 | 2.03E-128 |
| Rps6kb1 | 2.42E-132 | 0.28927004 | 0.142 | 0.057 | 6.54E-128 |
| Eif4g1 | 3.86E-132 | 0.29170061 | 0.125 | 0.046 | 1.04E-127 |
| Pnpla8 | 5.41E-132 | 0.35027521 | 0.107 | 0.035 | 1.46E-127 |
| Nfic | 6.66E-132 | 0.30649013 | 0.155 | 0.066 | 1.80E-127 |
| Tle1 | 1.46E-131 | 0.38143112 | 0.114 | 0.039 | 3.95E-127 |
| Aebp1 | 3.90E-131 | 0.5186533 | 0.117 | 0.042 | 1.05E-126 |
| Plekhm3 | 1.04E-130 | 0.2506781 | 0.163 | 0.071 | 2.81E-126 |
| 2210408I21R | 1.15E-130 | 0.36605833 | 0.132 | 0.051 | 3.10E-126 |
| Pde7b | 2.19E-130 | -0.9808206 | 0.305 | 0.369 | 5.93E-126 |
| Psmd14 | 3.69E-130 | 0.32800606 | 0.131 | 0.05 | 9.97E-126 |
| Hmgb1 | 6.59E-130 | 0.3476425 | 0.149 | 0.062 | 1.78E-125 |
| Agbl1 | 1.18E-129 | -0.5863388 | 0.238 | 0.338 | 3.18E-125 |
| Junos | 2.89E-129 | 0.43549236 | 0.11 | 0.037 | 7.82E-125 |
| Prdm2 | 3.06E-129 | 0.26741005 | 0.158 | 0.067 | 8.28E-125 |
| Pcmtd1 | 4.22E-129 | 0.33218741 | 0.11 | 0.037 | 1.14E-124 |
| Pdpk1 | 7.45E-129 | 0.28282709 | 0.189 | 0.089 | 2.01E-124 |
| Slc25a36 | 1.16E-128 | 0.27327232 | 0.146 | 0.06 | 3.15E-124 |
| Larp1 | 1.58E-128 | 0.26874192 | 0.166 | 0.073 | 4.27E-124 |
| Ttc3 | 2.49E-128 | 0.26862366 | 0.141 | 0.057 | 6.73E-124 |
| Abtb2 | 4.25E-128 | 0.35318372 | 0.129 | 0.049 | 1.15E-123 |
| Bzw2 | 2.34E-127 | 0.29138122 | 0.145 | 0.059 | 6.32E-123 |
| Tra2b | 3.66E-127 | 0.25951113 | 0.126 | 0.048 | 9.90E-123 |
| Zfp207 | 4.78E-127 | 0.31066799 | 0.128 | 0.049 | 1.29E-122 |
| Uggt1 | 4.90E-127 | 0.36664671 | 0.106 | 0.035 | 1.33E-122 |
| Vps54 | 5.85E-127 | 0.30496686 | 0.139 | 0.056 | 1.58E-122 |
| Parp4 | 3.15E-126 | 0.4154185 | 0.103 | 0.034 | 8.53E-122 |
| Gprasp1 | 5.73E-126 | 0.32758755 | 0.118 | 0.043 | 1.55E-121 |
| Rsbn1l | 7.93E-126 | 0.28667621 | 0.137 | 0.055 | 2.15E-121 |
| Armc8 | 1.78E-125 | 0.25390108 | 0.134 | 0.053 | 4.81E-121 |
| Bmp2k | 1.99E-125 | 0.34065465 | 0.105 | 0.035 | 5.38E-121 |
| Cdc37l1 | 4.84E-125 | 0.32771513 | 0.119 | 0.044 | 1.31E-120 |
| Jmy | 7.96E-125 | 0.34556714 | 0.116 | 0.042 | 2.15E-120 |
| Atf2 | 1.50E-124 | 0.33925327 | 0.124 | 0.047 | 4.05E-120 |
| Rbpj | 2.81E-124 | 0.37920176 | 0.13 | 0.052 | 7.61E-120 |
| Samd4 | 7.47E-124 | 0.29552183 | 0.76 | 0.565 | 2.02E-119 |
| Ptpn3 | 1.15E-123 | 0.34335187 | 0.166 | 0.075 | 3.10E-119 |
| Pfkfb4 | 1.39E-123 | 0.34309599 | 0.103 | 0.034 | 3.76E-119 |
| Zfp516 | 1.74E-123 | 0.25566869 | 0.13 | 0.051 | 4.70E-119 |
| Sptan1 | 2.83E-123 | 0.40682259 | 0.119 | 0.045 | 7.65E-119 |
| Scmh1 | 9.36E-123 | 0.2909594 | 0.147 | 0.063 | 2.53E-118 |
| Ptbp2 | 7.68E-122 | 0.27864865 | 0.108 | 0.037 | 2.08E-117 |

| | | | | | |
|---|---|---|---|---|---|
| 2610037D02I | 3.46E-121 | 0.30286583 | 0.124 | 0.048 | 9.35E-117 |
| Tasp1 | 6.05E-121 | 0.2721628 | 0.128 | 0.051 | 1.64E-116 |
| Psmc6 | 6.89E-121 | 0.34552181 | 0.112 | 0.04 | 1.86E-116 |
| Tsc1 | 1.53E-120 | 0.36711519 | 0.104 | 0.036 | 4.14E-116 |
| Xrcc4 | 2.95E-120 | 0.33388897 | 0.113 | 0.041 | 7.98E-116 |
| Zfp346 | 4.70E-120 | 0.38610156 | 0.114 | 0.042 | 1.27E-115 |
| Rspry1 | 9.67E-119 | 0.39372118 | 0.115 | 0.043 | 2.61E-114 |
| Rlf | 1.00E-118 | 0.26546349 | 0.144 | 0.062 | 2.70E-114 |
| Arhgap5 | 2.76E-118 | 0.25678753 | 0.143 | 0.061 | 7.46E-114 |
| Kpna1 | 3.45E-118 | 0.28428225 | 0.131 | 0.054 | 9.32E-114 |
| Spata5 | 3.53E-118 | 0.2651501 | 0.128 | 0.051 | 9.53E-114 |
| Eea1 | 3.79E-118 | 0.25417152 | 0.107 | 0.038 | 1.02E-113 |
| Ppfibp1 | 3.83E-117 | 0.2870181 | 0.122 | 0.048 | 1.03E-112 |
| Ylpm1 | 4.76E-117 | 0.31741733 | 0.121 | 0.048 | 1.29E-112 |
| 1810026B05I | 5.10E-117 | 0.30058915 | 0.117 | 0.045 | 1.38E-112 |
| Arntl | 7.39E-117 | 0.3194749 | 0.11 | 0.04 | 2.00E-112 |
| Naca | 1.10E-116 | 0.2946654 | 0.118 | 0.045 | 2.99E-112 |
| Dido1 | 1.28E-116 | 0.30789321 | 0.141 | 0.061 | 3.47E-112 |
| Cwf19l2 | 2.75E-116 | 0.25366595 | 0.1 | 0.034 | 7.44E-112 |
| Tgfbr3 | 2.91E-116 | 0.30220471 | 0.199 | 0.102 | 7.88E-112 |
| Sacm1l | 4.53E-116 | 0.26278724 | 0.142 | 0.061 | 1.23E-111 |
| Vps13a | 9.46E-116 | 0.2502574 | 0.133 | 0.055 | 2.56E-111 |
| Kdm5a | 1.38E-115 | 0.28019857 | 0.146 | 0.064 | 3.73E-111 |
| Ppard | 1.92E-115 | 0.27652207 | 0.14 | 0.06 | 5.20E-111 |
| Ubl3 | 2.92E-115 | 0.29394588 | 0.138 | 0.059 | 7.89E-111 |
| Slc27a1 | 4.17E-115 | 0.4040703 | 0.106 | 0.039 | 1.13E-110 |
| Amer1 | 5.24E-115 | 0.30937944 | 0.128 | 0.052 | 1.42E-110 |
| Ddx6 | 2.84E-113 | 0.322761 | 0.119 | 0.047 | 7.67E-109 |
| Psmb7 | 1.12E-112 | 0.2800064 | 0.107 | 0.039 | 3.02E-108 |
| Ngly1 | 1.41E-112 | 0.30813716 | 0.104 | 0.038 | 3.82E-108 |
| Uqcc1 | 7.35E-112 | 0.25426798 | 0.129 | 0.054 | 1.99E-107 |
| Myo10 | 8.58E-112 | 0.36886458 | 0.112 | 0.043 | 2.32E-107 |
| Nemf | 1.79E-111 | 0.31996993 | 0.105 | 0.038 | 4.84E-107 |
| Dgkd | 1.82E-111 | 0.25105949 | 0.112 | 0.043 | 4.92E-107 |
| Cdyl | 1.86E-111 | 0.25671245 | 0.116 | 0.046 | 5.03E-107 |
| Eef1a2 | 1.73E-110 | 0.50508475 | 0.128 | 0.056 | 4.69E-106 |
| Pygm | 2.69E-110 | -0.9828607 | 0.314 | 0.362 | 7.26E-106 |
| Pik3ca | 3.28E-110 | 0.28436272 | 0.12 | 0.048 | 8.88E-106 |
| Fhit | 6.27E-110 | 0.27250672 | 0.242 | 0.135 | 1.69E-105 |
| Zdhhc17 | 7.27E-110 | 0.29920253 | 0.101 | 0.036 | 1.97E-105 |
| Sptbn1 | 1.94E-109 | 0.41053653 | 0.126 | 0.054 | 5.25E-105 |
| Sec63 | 3.79E-109 | 0.27909103 | 0.108 | 0.041 | 1.02E-104 |
| Usp40 | 3.81E-109 | 0.25994985 | 0.11 | 0.042 | 1.03E-104 |

| | | | | | |
|---|---|---|---|---|---|
| Ctdsp2 | 4.28E-108 | 0.29835335 | 0.118 | 0.048 | 1.16E-103 |
| Arih2 | 7.13E-108 | 0.28379815 | 0.124 | 0.052 | 1.93E-103 |
| Abca8b | 7.90E-108 | 0.37822465 | 0.114 | 0.046 | 2.14E-103 |
| 2310001H17I | 1.18E-107 | 0.26914724 | 0.142 | 0.064 | 3.18E-103 |
| Asb15 | 1.36E-107 | 0.25307471 | 0.151 | 0.07 | 3.69E-103 |
| Ubxn4 | 3.86E-107 | 0.25870713 | 0.163 | 0.078 | 1.04E-102 |
| Pcx | 5.10E-107 | -1.2997838 | 0.121 | 0.188 | 1.38E-102 |
| Gm35330 | 5.17E-107 | 0.31073819 | 0.147 | 0.068 | 1.40E-102 |
| Sos2 | 6.16E-107 | 0.26790358 | 0.119 | 0.049 | 1.67E-102 |
| Inpp4b | 1.56E-106 | 0.32125897 | 0.145 | 0.067 | 4.22E-102 |
| Man1a2 | 1.36E-105 | 0.29505772 | 0.1 | 0.036 | 3.69E-101 |
| Tent2 | 6.35E-105 | 0.2554057 | 0.106 | 0.041 | 1.72E-100 |
| Tnni2 | 1.43E-103 | 0.29593409 | 0.145 | 0.069 | 3.87E-99 |
| Twf2 | 2.09E-103 | 0.33966068 | 0.115 | 0.047 | 5.65E-99 |
| Xpo4 | 2.34E-103 | 0.28102574 | 0.105 | 0.04 | 6.32E-99 |
| Dhx32 | 2.71E-103 | 0.25167876 | 0.106 | 0.041 | 7.33E-99 |
| Plcd4 | 4.54E-103 | -1.3046229 | 0.098 | 0.16 | 1.23E-98 |
| Prkag3 | 5.04E-103 | -1.0887534 | 0.216 | 0.279 | 1.36E-98 |
| Myo5a | 1.76E-102 | 0.32449928 | 0.115 | 0.047 | 4.76E-98 |
| Sox6 | 3.14E-102 | -0.4128059 | 0.887 | 0.851 | 8.49E-98 |
| Atp13a3 | 5.20E-102 | 0.38238279 | 0.117 | 0.05 | 1.41E-97 |
| Zfp652 | 6.01E-101 | 0.26985539 | 0.143 | 0.067 | 1.63E-96 |
| Faf2 | 7.20E-101 | 0.31745128 | 0.103 | 0.04 | 1.95E-96 |
| Pcmtd2 | 1.96E-100 | 0.2563792 | 0.11 | 0.044 | 5.30E-96 |
| Luc7l | 2.47E-100 | 0.30941291 | 0.105 | 0.041 | 6.69E-96 |
| Evi5 | 3.30E-100 | 0.25736032 | 0.124 | 0.054 | 8.92E-96 |
| Mxd4 | 2.39E-99 | 0.28994473 | 0.101 | 0.039 | 6.46E-95 |
| Agl | 1.05E-97 | -0.7816363 | 0.487 | 0.487 | 2.83E-93 |
| Sh3bgr | 1.55E-96 | -1.2884434 | 0.114 | 0.174 | 4.20E-92 |
| mt-Co1 | 9.97E-96 | -0.874376 | 0.127 | 0.196 | 2.70E-91 |
| Grb10 | 4.94E-92 | 0.25280135 | 0.108 | 0.046 | 1.34E-87 |
| Shprh | 5.48E-91 | 0.28430295 | 0.109 | 0.047 | 1.48E-86 |
| Ank2 | 1.54E-90 | -0.7789343 | 0.397 | 0.447 | 4.18E-86 |
| mt-Nd5 | 3.53E-90 | -0.9506599 | 0.095 | 0.157 | 9.54E-86 |
| Atrnl1 | 4.34E-89 | 0.26469659 | 0.151 | 0.077 | 1.17E-84 |
| Prdx1 | 2.70E-88 | 0.28872501 | 0.109 | 0.047 | 7.29E-84 |
| mt-Atp6 | 4.60E-86 | -0.7238533 | 0.167 | 0.238 | 1.25E-81 |
| Pkig | 1.37E-83 | 0.25323336 | 0.106 | 0.047 | 3.71E-79 |
| Rpl3l | 6.74E-82 | -1.1084521 | 0.13 | 0.185 | 1.82E-77 |
| Hadhb | 2.90E-78 | 0.25526264 | 0.1 | 0.045 | 7.83E-74 |
| Gab2 | 7.88E-77 | 0.2762541 | 0.103 | 0.047 | 2.13E-72 |
| Sorbs2 | 1.35E-73 | -0.9321757 | 0.213 | 0.266 | 3.64E-69 |
| Smoc2 | 1.67E-72 | 0.26950606 | 0.127 | 0.066 | 4.53E-68 |

| | | | | | |
|---|---|---|---|---|---|
| Ppara | 2.00E-69 | -1.1986793 | 0.153 | 0.205 | 5.41E-65 |
| Gm28653 | 5.76E-69 | -1.310684 | 0.133 | 0.188 | 1.56E-64 |
| Myoz3 | 5.92E-69 | -1.0764707 | 0.066 | 0.111 | 1.60E-64 |
| Myh1 | 1.72E-66 | 0.4271682 | 0.236 | 0.158 | 4.66E-62 |
| Ampd1 | 6.30E-60 | -0.8708409 | 0.301 | 0.327 | 1.70E-55 |
| Rhobtb1 | 9.85E-50 | -0.9702928 | 0.16 | 0.199 | 2.66E-45 |
| Kcnma1 | 3.55E-49 | -0.3636827 | 0.799 | 0.745 | 9.60E-45 |
| Ank3 | 8.88E-47 | -0.4012641 | 0.749 | 0.687 | 2.40E-42 |
| Klhl33 | 9.23E-47 | -1.0531233 | 0.137 | 0.176 | 2.50E-42 |
| Arhgap24 | 2.55E-45 | 0.25367386 | 0.1 | 0.057 | 6.88E-41 |
| Mlxipl | 5.60E-37 | -0.9472 | 0.184 | 0.212 | 1.52E-32 |
| Myh2 | 2.38E-34 | 0.38701556 | 0.162 | 0.115 | 6.44E-30 |
| Map2k6 | 5.37E-31 | -0.9474063 | 0.098 | 0.127 | 1.45E-26 |
| Ttc7b | 3.11E-30 | -0.3018742 | 0.215 | 0.148 | 8.40E-26 |
| Il31ra | 1.39E-29 | -0.7195637 | 0.181 | 0.21 | 3.76E-25 |
| Ckm | 9.25E-29 | -0.6115205 | 0.398 | 0.413 | 2.50E-24 |
| Kcnq5 | 6.63E-28 | -0.3477239 | 0.371 | 0.389 | 1.79E-23 |
| Camk2g | 8.24E-27 | -0.267953 | 0.179 | 0.124 | 2.23E-22 |
| 2310016D03 | 1.36E-26 | -0.8464443 | 0.075 | 0.101 | 3.68E-22 |
| Gphn | 2.00E-26 | -0.3138417 | 0.9 | 0.785 | 5.42E-22 |
| Ptp4a2 | 1.40E-25 | -0.2529907 | 0.179 | 0.126 | 3.79E-21 |
| Cdc42 | 2.16E-24 | -0.2520935 | 0.173 | 0.122 | 5.84E-20 |
| D430041D05 | 4.12E-24 | -0.8201419 | 0.105 | 0.131 | 1.11E-19 |
| Mical2 | 1.32E-23 | -0.250774 | 0.255 | 0.185 | 3.57E-19 |
| Ckmt2 | 1.49E-23 | -0.8624815 | 0.161 | 0.184 | 4.02E-19 |
| Cobl | 7.82E-23 | -0.2842035 | 0.147 | 0.103 | 2.11E-18 |
| Abcc5 | 4.37E-22 | -0.2626436 | 0.141 | 0.098 | 1.18E-17 |
| Ppargc1a | 8.41E-22 | -0.3693926 | 0.162 | 0.116 | 2.27E-17 |
| D5Ertd579e | 8.44E-22 | -0.3358839 | 0.258 | 0.188 | 2.28E-17 |
| Cuedc1 | 5.91E-20 | -0.2946924 | 0.292 | 0.214 | 1.60E-15 |
| St3gal3 | 7.39E-20 | -0.2990291 | 0.223 | 0.164 | 2.00E-15 |
| Arl6ip5 | 1.24E-19 | -0.8033501 | 0.203 | 0.215 | 3.35E-15 |
| Atxn1 | 1.38E-19 | -0.3712381 | 0.67 | 0.582 | 3.74E-15 |
| Itpr1 | 1.46E-19 | -0.2825999 | 0.163 | 0.118 | 3.95E-15 |
| Neurl1a | 2.30E-19 | -0.252536 | 0.103 | 0.071 | 6.23E-15 |
| Slc10a7 | 1.18E-18 | -0.2646858 | 0.137 | 0.098 | 3.19E-14 |
| Cacna1s | 1.72E-18 | -0.3294256 | 0.183 | 0.134 | 4.66E-14 |
| Best3 | 2.55E-18 | -0.8010415 | 0.082 | 0.102 | 6.90E-14 |
| Sdccag8 | 7.71E-18 | -0.2617712 | 0.205 | 0.153 | 2.08E-13 |
| Zc3h13 | 2.42E-17 | -0.2781553 | 0.108 | 0.077 | 6.54E-13 |
| Macf1 | 1.12E-16 | -0.3116956 | 0.271 | 0.205 | 3.03E-12 |
| Fhod3 | 3.21E-15 | -0.56282 | 0.331 | 0.317 | 8.69E-11 |
| Optn | 4.03E-15 | -0.2983977 | 0.131 | 0.097 | 1.09E-10 |

| | | | | | |
|---|---|---|---|---|---|
| Psd3 | 6.55E-15 | -0.3800592 | 0.394 | 0.388 | 1.77E-10 |
| Fnip1 | 1.14E-14 | -0.3764143 | 0.274 | 0.206 | 3.07E-10 |
| Slco5a1 | 1.95E-14 | -0.3797754 | 0.202 | 0.153 | 5.26E-10 |
| Smad3 | 3.81E-14 | -0.3245912 | 0.144 | 0.108 | 1.03E-09 |
| Macrod1 | 4.42E-14 | -0.3009151 | 0.283 | 0.216 | 1.19E-09 |
| Zfp423 | 5.56E-14 | -0.2925365 | 0.105 | 0.077 | 1.50E-09 |
| Alpk3 | 1.42E-13 | -0.5536797 | 0.548 | 0.474 | 3.83E-09 |
| Slc4a4 | 1.43E-13 | -0.6269498 | 0.277 | 0.273 | 3.87E-09 |
| Tom1l2 | 1.47E-13 | -0.5350217 | 0.453 | 0.404 | 3.96E-09 |
| 1700025G04 | 2.65E-13 | -0.3062147 | 0.271 | 0.207 | 7.16E-09 |
| Stxbp4 | 5.64E-13 | -0.3140192 | 0.157 | 0.12 | 1.53E-08 |
| Ahdc1 | 7.83E-13 | -0.3644705 | 0.134 | 0.101 | 2.12E-08 |
| Itgb6 | 3.30E-12 | -0.2989834 | 0.175 | 0.134 | 8.91E-08 |
| Slc25a12 | 4.68E-12 | -0.3365873 | 0.342 | 0.259 | 1.27E-07 |
| Actn2 | 5.94E-12 | -0.8691128 | 0.351 | 0.334 | 1.61E-07 |
| Casq1 | 6.73E-12 | -0.3209768 | 0.103 | 0.077 | 1.82E-07 |
| Pdzrn3 | 1.17E-11 | -0.3566406 | 0.524 | 0.484 | 3.16E-07 |
| Nav2 | 1.28E-11 | -0.3384508 | 0.258 | 0.2 | 3.46E-07 |
| Zbtb16 | 3.32E-11 | -0.2912967 | 0.75 | 0.647 | 8.97E-07 |
| St3gal6 | 5.10E-11 | -0.357385 | 0.171 | 0.132 | 1.38E-06 |
| Acvr1 | 6.03E-11 | -0.3667218 | 0.111 | 0.084 | 1.63E-06 |
| Cox10 | 6.54E-11 | -0.3457904 | 0.117 | 0.089 | 1.77E-06 |
| 1700012D14 | 8.20E-11 | -0.3070594 | 0.301 | 0.235 | 2.22E-06 |
| Eno3 | 1.64E-10 | -0.6751543 | 0.092 | 0.106 | 4.44E-06 |
| Rcan2 | 1.69E-10 | -0.2668508 | 0.148 | 0.117 | 4.57E-06 |
| Pfkm | 2.44E-10 | -0.4142301 | 0.163 | 0.126 | 6.60E-06 |
| Jph2 | 2.80E-10 | -0.295392 | 0.277 | 0.218 | 7.58E-06 |
| Qrich1 | 5.06E-10 | -0.2868994 | 0.142 | 0.111 | 1.37E-05 |
| Ccdc50 | 1.39E-09 | -0.670313 | 0.296 | 0.278 | 3.77E-05 |
| Mrtfa | 3.23E-09 | -0.3770303 | 0.129 | 0.101 | 8.74E-05 |
| Stau2 | 4.35E-09 | -0.3225698 | 0.238 | 0.189 | 0.00011752 |
| Large1 | 8.95E-09 | -0.3389825 | 0.386 | 0.302 | 0.00024207 |
| 5430431A17 | 1.55E-08 | -0.6734327 | 0.095 | 0.106 | 0.0004178 |
| Cmya5 | 2.06E-08 | -0.2874664 | 0.737 | 0.633 | 0.00055587 |
| Tcea3 | 2.64E-08 | -0.4134964 | 0.119 | 0.093 | 0.00071364 |
| Rabgef1 | 2.78E-08 | -0.4289259 | 0.161 | 0.128 | 0.00075093 |
| Musk | 8.81E-08 | -0.3572211 | 0.194 | 0.157 | 0.00238283 |
| Nexn | 1.50E-07 | -0.3344603 | 0.348 | 0.274 | 0.00406744 |
| mt-Rnr2 | 7.17E-07 | -0.2696529 | 0.78 | 0.617 | 0.019378 |
| Cbfb | 7.55E-07 | -0.3654382 | 0.124 | 0.1 | 0.02041061 |
| Msrb3 | 8.16E-07 | -0.3238866 | 0.334 | 0.266 | 0.02207098 |
| Dag1 | 9.38E-07 | -0.3238215 | 0.292 | 0.236 | 0.02536417 |
| Actn3 | 1.16E-06 | -0.60349 | 0.21 | 0.214 | 0.03141294 |

| | | | | | |
|---|---|---|---|---|---|
| Gm50194 | 1.40E-06 | -0.706566 | 0.163 | 0.165 | 0.03798241 |
